# Biosynthesis of modular signaling molecules requires functional diversification of carboxylesterases in *Pristionchus pacificus*

**DOI:** 10.1101/2025.03.28.645926

**Authors:** Pei Zhang, Penghieng Theam, Wenya Xu, Shudan Ye, Hanh Witte, Tingwei Liu, Ralf J. Sommer, Chuanfu Dong

**Affiliations:** The Key Laboratory of Cell Proliferation and Regulation Biology, Ministry of Education, Department of Biology, College of Life Sciences, Beijing Normal University, 100875 Beijing, China; Department for Integrative Evolutionary Biology, Max Planck Institute for Biology Tübingen, Max Planck Ring 9, 72076 Tübingen, Germany; Center for Biological Science and Technology, Advanced Institute of Natural Sciences, Beijing Normal University, 519087 Zhuhai, China; Department of Biology, Faculty of Arts and Sciences, Beijing Normal University, 519087 Zhuhai, China; Zhuhai-Macao Biotechnology Joint Laboratory, Beijing Normal University, 519087 Zhuhai, China

**Keywords:** *Pristionchus pacificus*, ascaroside, pheromone, biosynthesis, *Caenorhabditis elegans*, modular signaling molecules, carboxylesterases

## Abstract

The model nematode *Pristionchus pacificus* produces four types of complex ascaroside pheromones named UBAS, DASC, NPAR, and PASC. However, the exact biosynthetic pathways of these modular signaling molecules remain enigmatic. We have previously identified a carboxylesterase *Ppa*-UAR-1 for the biosynthesis of UBAS, enabling the attachment of ureidoisobutyric acid at the 4’-position of simple ascarosides. Here, we report three new carboxylesterases *Ppa*-UAR-5, *Ppa*-UAR-6 and *Ppa*-UAR-12 from *P. pacificus*. *Ppa*-UAR-5 functions downstream of *Ppa*-UAR-1 to furnish the biosynthesis of ubas#1 and ubas#2, whereas *Ppa*-UAR-12 specifically links two ascr#1 at the 4’-position to synthesize dasc#1. Finally, *Ppa*-UAR-6 is essential for the biosynthesis of npar#1-3 and part#9. The expression patterns of *Ppa-uar-6* and *Ppa-uar-12* in the intestinal and epidermal cells suggest pheromone biosynthesis to be restricted to specific tissues. These findings indicate that the expansion and functional diversification of carboxylesterases plays a crucial role in the evolution of complex pheromones in nematodes.

**SIGNIFICANCE:** Our study identified three new carboxylesterase genes (*Ppa-uar-5*, *Ppa-uar-12* and *Ppa-uar-6*) from *P. pacificus*. The encoded enzymes separately regulated the biosynthesis of three types of modular pheromones (UBAS, DASC, and NPAR-types) in worms’ intestinal cells and exhibited extremely high substrate-specificity. For example, we verified that *Ppa*-UAR-12 specifically linked one ascr#1 to the 4’-position of another ascr#1 to finally furnish the biosynthesis of ascaroside dimer dasc#1. These new discoveries largely clarify the delicate biosynthetic mechanisms of how diversification of carboxylesterases functions in *P. pacificus* to build up enormous complexity of modular signaling molecules. Such a biosynthetic mechanism of nematode pheromones might be widely conserved in the phylum Nematoda.

## INTRODUCTION

The necromenic roundworm *Pristionchus pacificus* is a model nematode for evolutionary developmental biology^1,2^. Most *Pristionchus* nematodes are found to be ecologically associated with scarab beetles^3–5^, normally arresting as dauer larvae (a developmentally stress-resistant larval stage) on living beetles. These nematodes resume normal development when bacteria proliferate on beetle carcasses and become available as food source. Like other nematodes of the family Diplogastridae, *P. pacificus* exhibits a mouth-form dimorphism with the so-called stenostomatous (St) or eurystomatous (Eu) forms, a striking example of phenotypic plasticity of feeding structures^6^. Specifically, St animals have a single tooth and feed on bacteria, whereas Eu animals are equipped with two sharp teeth enabling them to predate on other nematodes^7–12^. As genome editing by CRISPR/Cas9 engineering is reliably established in *P. pacificus*^13–16^, this powerful technique allows the in-depth investigation of the regulatory mechanism of mouth-form development and its associated predatory behavior. For example, the combination of forward and reverse genetic tools resulted in the identification of the major regulator of mouth-form plasticity, *eud-1*^17^ and the self-recognition locus *self-1* that prevents cannibalism against genetically identical kin^12^. In addition, *P. pacificus* excretes signaling molecules referred to as “nematode-derived modular metabolites”, a group of ascaroside pheromones initially characterized from *Caenorhabditis elegans*^18–32^, which are also found to influence the development of the mouth-form and dauer formation^33–35^.

Similar to *C. elegans*, *P. pacificus* produces a wide variety of signaling ascarosides ^33,36,37^, however, unlike *C. elegans* only three simple ascarosides—ascr#9 (asc-C5), ascr#12 (asc-C6) and ascr#1 (asc-C7)—have so far been identified (Figure 1A). These simple ascarosides harbor an ascarylose sugar linked to the aglycone of a short fatty acid side chain. These aglycones are derived from long chain fatty acids via several cycles of peroxisomal *β*-oxidation^38–44^. In addition, these simple ascarosides can act as core structures (Figure 1A) that are further decorated with additional building blocks to generate three types of complex modular ascarosides, named UBAS, DASC and PASC (Figure 1A). Indeed, each type of modular ascarosides incorporates different kinds of intermediates originating from various primary metabolic pathways. For example, UBAS ascarosides carry a common unit of pyrimidine metabolic pathway-derived ureidoisobutyric acid at the 4’-position, with some complex UBAS ascarosides, like ubas#1 and ubas#2, even separately containing an additional building block of oscr#9 (asc-ꞷC5) and ascr#12 (asc-C6) at the 2’-position. DASC ascarosides are normally comprised of two units of simple ascarosides, which are linked at the 2’- or 4’-position. In addition, PASC ascarosides harbor a succinate-based phenylethanolamine at their terminal carboxyl group of fatty acid side chain. Finally, *P. pacificus* produces a group of unique NPAR paratosides including part#9 and npar#1-3. Based on the structural skeleton of part#9 that is an epimer of ascr#9, npar#1-3 are integrated with additional groups of adenosine-based threonine at their C5 fatty acid side chains (Figure 1A). So far, only *P. pacificus* and its closely related species from the genus *Pristionchus* have been reported to produce these specialized metabolites^33,37^.

**Figure 1.**
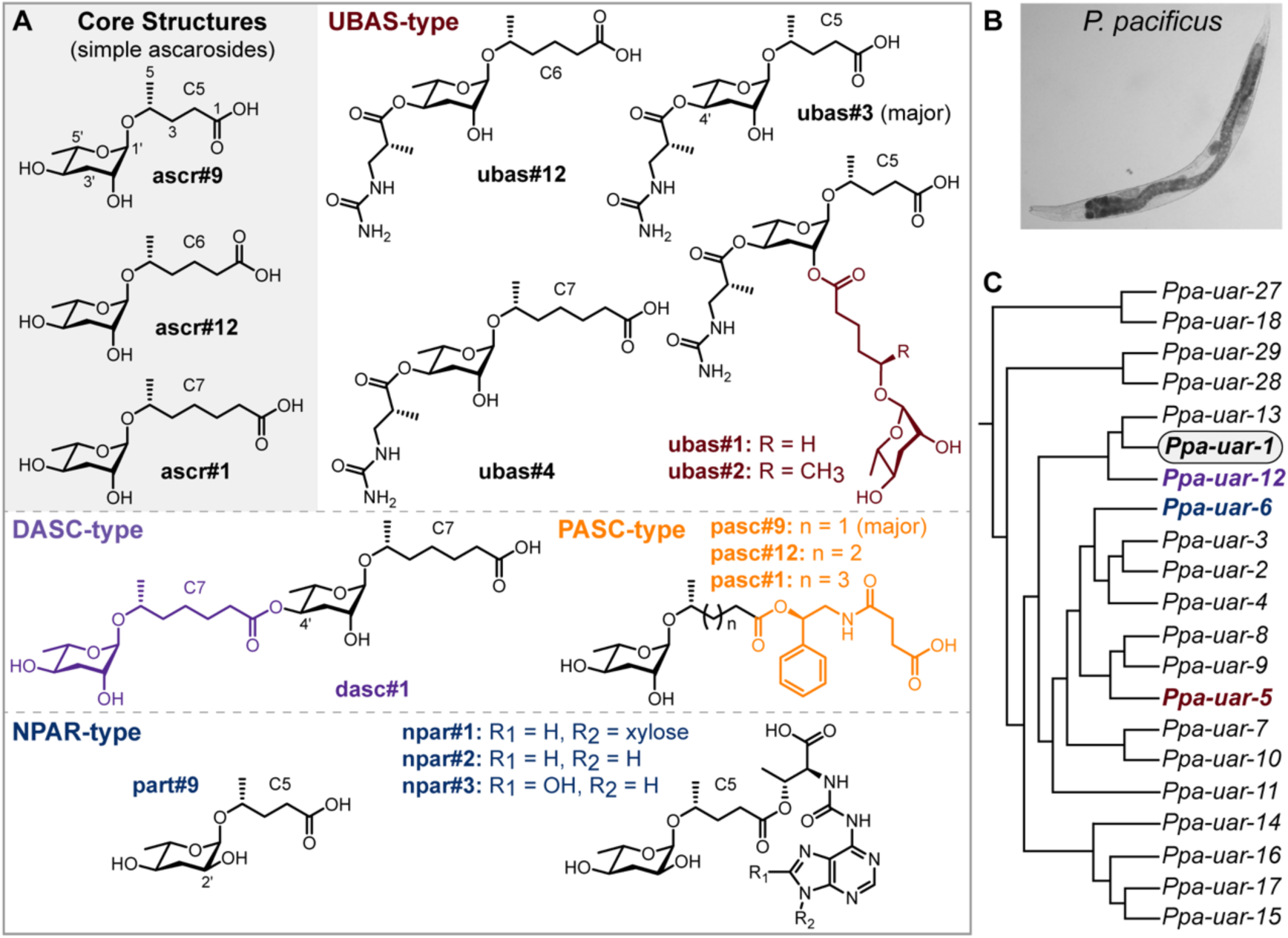
Pheromonal components and their biosynthetic genes from *P. pacificus*. (A) Chemical structures of simple ascarosides and four types of chemically modified modular pheromones including UBAS, DASC, PASC, and NPAR. (B) Confocal image of *P. pacificus* nematode. (C) 20 out of 76 candidate genes (Tables S1 and S2) with homology to *Ppa-uar-1*^49^ were knocked out by CRISPR/Cas9, from which *Ppa-uar-5*, *Ppa-uar-12*, and *Ppa-uar-6* genes were shown to regulate the biosynthesis of UBAS, DASC, and NPAR, respectively.

These decorated signaling molecules play critical roles in the regulation of various developmental processes in *P. pacificus* (Figure 1B). For example, the dimeric ascaroside dasc#1 is able to trigger the formation of the Eu mouth-form^33^, while UBAS, PASC and NPAR pheromones are capable of inducing dauer formation^7,33,34,36,45–48^. Given this complex biochemistry that is in part linked to the developmental decisions which respond to changing environmental influences, we seek to investigate the biosynthetic origins of these modular signaling molecules. While little is known about these biosynthetic pathways, we and co-workers have previously discovered a key gene *Ppa-uar-1* in the genome of *P. pacificus* by analyzing both metabolomes and transcriptomes of 264 *P. pacificus* wild isolates^49^. Differences in the metabolome of independent wild isolates were linked to the *Ppa-uar-1* gene through a genome-wide association study (GWAS) mapping. *Ppa-uar-1* mutants, analyzed by LC-MS (Liquid Chromatography Mass Spectrometry) of both worm culture medium (*exo*-metabolomes) and worm pellets (*endo*-metabolomes), showed the absence of UBAS ascarosides, which are characteristic of the attachment of a ureidoisobutyric acid unit at the 4’-position (Figure 1A). These results indicate for the first time, that the carboxylesterase *Ppa*-UAR-1 is capable of linking ureidoisobutyric acid to the 4’-position of simple ascarosides to further produce a group of modular UBAS ascarosides (Figure 1A)^49^. However, up to date, the downstream factors of the biosynthetic pathway of ubas#1 and ubas#2 have still been elusive, i.e., the exact mechanism of how additional units of oscr#9 and ascr#12 are incorporated into the 2’-position of ubas#3 remains unknown. In addition, the biosynthetic origins of the remaining types of modular signaling molecules including DASC, NPAR and PASC are completely uninvestigated. Therefore, it will be important to explore the mechanisms of how different building blocks are integrated into the core structures of simple ascarosides for the biosynthesis of these structurally complex pheromones.

Here, we report the discovery of three new carboxylesterase proteins *Ppa*-UAR-5, *Ppa*-UAR-12 and *Ppa*-UAR-6 in *P. pacificus*, which are required for the biosynthesis of UBAS, DASC and NPAR-type pheromones, respectively (Figure 1C). Knockout mutants of *Ppa-uar-5*, *Ppa-uar-12* and *Ppa-uar-6* led to the loss of specific sets of pheromones in both *exo*-metabolomes and *endo*-metabolomes. *Ppa-uar-6* and *Ppa-uar-12* were further observed to be specifically expressed in the intestine of *P. pacificus*, which highlights the importance of this tissue for pheromone biosynthesis. The discovery of these biosynthetic proteins provides new opportunities for uncovering the evolution and biological functions of signaling molecules in nematodes.

## RESULTS

### Knockout of 20 homologs of *Ppa-uar-1* by CRISPR/Cas9

To explore the biosynthetic origins of structurally complex pheromones in *P. pacificus*, we hypothesized that homologs of the carboxylesterase *Ppa*-UAR-1^49^ might play important roles in the formation of these signaling molecules. This hypothesis was based on similar inferences in *C. elegans*^50–51^ that have resulted in the identification of novel biosynthesis regulators after the original findings by Falcke et al.^49^ in *P. pacificus*. Bioinformatic analysis of the *P. pacificus* genome revealed 75 homologs of *Ppa-uar-1*^49^ and they were renamed as *Ppa-uar-x* (*x* marked from 2 to 76) (Tables S1 and S2). The amino acid sequences of the homologous carboxylesterase proteins were properly aligned and 20 of them were found to be most closely related to *Ppa*-UAR-1 in the phylogenetic tree (Figures 1C and S1). Thus, we were motivated to concentrate on these 20 *uar* genes, all of which were targeted for the creation of knockout mutants through CRISPR/Cas9 engineering (Figure 1C; Tables S1 and S3). We indeed succeeded in generating presumptive null mutants with a total of 57 mutant alleles in these 20 genes (Table S1). To study these mutants further, we cultured two or three mutant alleles per gene in S-medium for the collection of *exo*-metabolomes and *endo*-metabolomes (Method details), followed by LC-MS examination of their pheromonal compositions, which were compared with those of wild type *P. pacificus* worms. This workflow supported the discovery of three new biosynthetic genes *Ppa-uar-5*, *Ppa-uar-6*, and *Ppa-uar-12* (Figure 1C). Similar to *Ppa-uar-1* in *P. pacificus*^49^ and the *cest* genes in *C. elegans*^32,50,51^, these three genes also encode carboxylesterase proteins (esterase or hydrolase) belonging to the α/β-hydrolase-fold enzyme superfamily with a predicted C-terminal transmembrane domain (Figure S1)^32,50,51^. As described in greater detail below, *Ppa*-UAR-5, *Ppa*-UAR-12, and *Ppa*-UAR-6 were discovered to separately regulate the biosynthesis of UBAS, DASC, and NPAR-type pheromones (Figure 1C). Unfortunately, we did not identify any candidate gene that was involved in the biosynthesis of PASC-type pheromones, the most abundantly produced class of pheromones (Figure 1A). In the following paragraphs, the biosynthetic capacities for each newly identified carboxylesterase are described in detail.

### *Ppa*-UAR-5 functions downstream of *Ppa*-UAR-1 to furnish the biosynthesis of ubas#1 and ubas#2

First, we generated three new alleles of *Ppa-uar-1* mutants by CRISPR/Cas9 (Table S1) to add to the previously reported *Ppa-uar-1* mutants^49^. Indeed, these new mutants did not produce any trace of UBAS-type ascarosides. Given that both ubas#1 and ubas#2 carry a common building block of ubas#3, these results suggested that ubas#3 might be a biosynthetic precursor of ubas#1 and ubas#2^49^. However, how the remaining two building blocks oscr#9 (asc-ꞷC5)^39^ and ascr#12 (asc-C6) (Figure 1A) are incorporated into the 2’-position of ubas#3 for the biosynthesis of ubas#1 and uabs#2 was still elusive.

To elucidate the biosynthetic pathways of ubas#1 and ubas#2, all of the above-mentioned 57 mutants were maintained in 50 ml liquid culture, and the resulting *exo*-metabolomes and *endo*-metabolomes were freeze-dried, extracted by methanol, concentrated and aliquoted for LC-MS analysis (Method details). Out of these mutants, CRISPR/Cas9 targeting exon 4 of *Ppa-uar-5* produced three mutant alleles, including *tu2019* that carried an insertion of 68 bp along with 5 single nucleotide polymorphisms (SNPs). Additionally, the *Ppa-uar-5* alleles *tu2020* and *tu2021* carried deletions of 11 bp and 4 bp, respectively (Figure 2A). We found that all the three alleles of *Ppa-uar-5* did not produce ubas#2 anymore, while only trace amounts of ubas#1 remained detectable in the *exo*-metabolomes (Figures 2B and 2C). Previously, tiny amounts of ubas#1 were also detected in *Ppa-uar-1* mutants^49^. However, in contrast to the complete absence of ubas#3 in *Ppa-uar-1*^49^, ubas#3 was abundantly produced in *Ppa-uar-5* (Figures 2B and 2C). Quantitative analysis of the three major UBAS ascarosides in the *exo*-metabolomes clearly showed that *Ppa-uar-5* mutants markedly produced much larger amounts of ubas#3 than wild type worms (Figure 2F), indicating that ubas#3 should act as the biosynthetic precursor of ubas#1 and ubas#2. Furthermore, LC-MS examination of the *endo*-metabolomes of *Ppa-uar-5* mutants also showed that very large amounts of ubas#3 were produced, whereas ubas#1 and ubas#2 completely disappeared (Figures 2D and 2E). Together, these experiments suggest that the absence of ubas#1 and ubas#2 was a result of the perturbation or blocking of their biosynthetic pathway instead of the excretory process. Thus, *Ppa*-UAR-5 should act on the intermediate precursor ubas#3 and is involved in the biosynthesis of ubas#1 and ubas#2. Therefore, we have identified a new enzyme, *Ppa*-UAR-5, that acts downstream of *Ppa*-UAR-1 and is involved in linking oscr#9 and ascr#12 to the 2’-position of ubas#3 to finally furnish the biosynthesis of ubas#1 and ubas#2 (Figure 2G).

**Figure 2.**
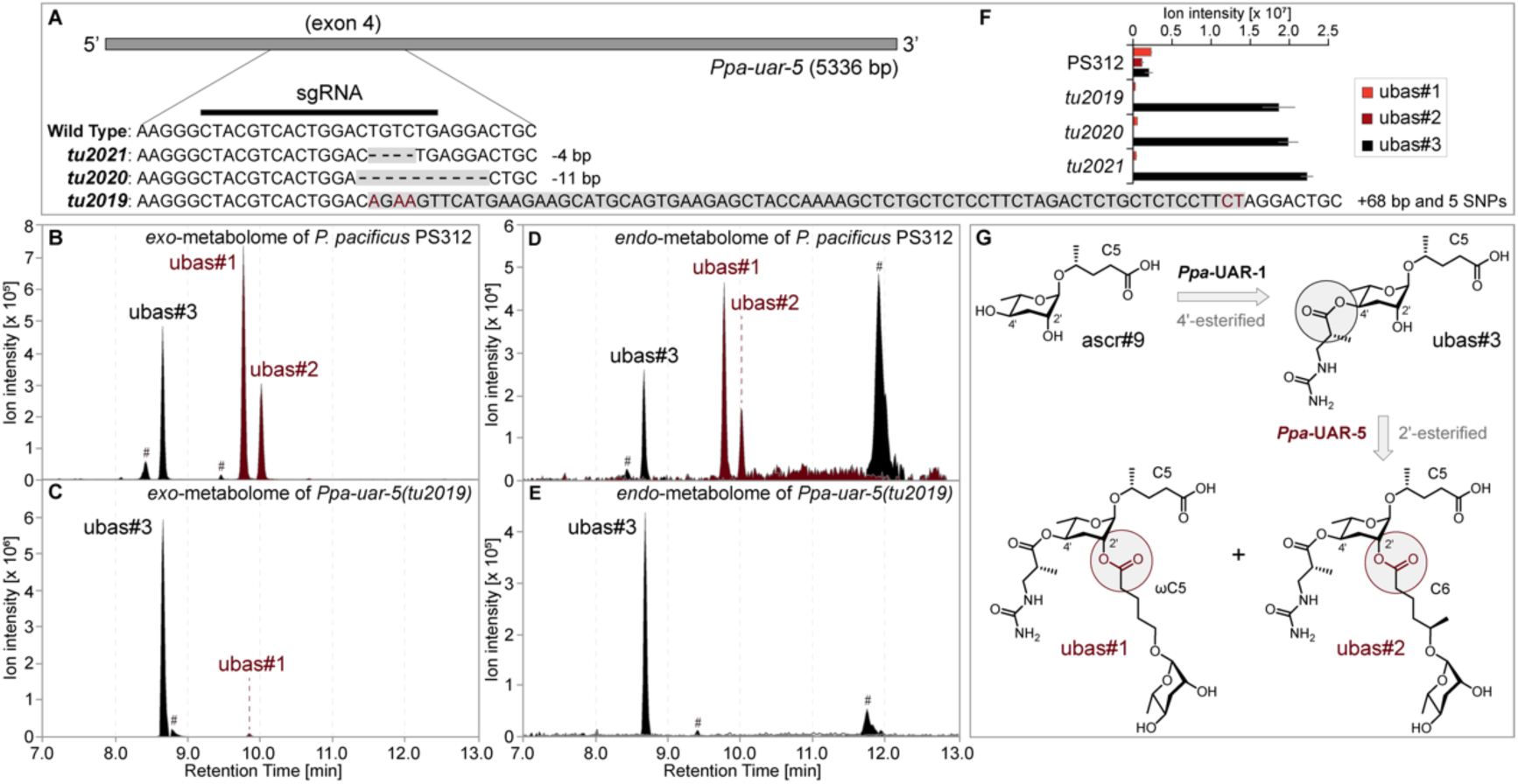
Biosynthetic investigation of UBAS-type pheromones. (A) Three mutant alleles of *Ppa-uar-5* were generated by CRISPR/Cas9. (B-E) The production of UBAS-type pheromones including ubas#1, ubas#2, and ubas#3 in the *exo*-metabolomes (B and C) and *endo*-metabolomes (D and E) of both wild type *P. pacificus* PS312 and mutants were examined by LC-MS. The pound signs (#) denote non-ascarosides. (F) Ion abundances of UBAS-type pheromones in the *exo*-metabolomes of wild type *P. pacificus* PS312 and three mutants were quantitatively analysed. Three biological replicates were performed for each allele. Error bars represent standard deviation (± SD). (G) Proposed biosynthetic pathway of UBAS-type pheromones.

### *Ppa*-UAR-12 regulates the biosynthesis of dasc#1

Utilizing two monomeric building blocks of simple ascarosides, many nematodes including *C. elegans*^52^ and its relatives^53^ are able to produce specific pheromonal content of dimeric ascarosides, each of which harbors an exclusive homo- or heterodimeric structure isomerized through a 2’- or 4’-ester linkage (Figures 1A and 3). Extensive chemical investigation of the *P. pacificus exo*-metabolomes also supports the identification of four dimeric ascarosides dasc#1, dasc#4, dasc#6, and dasc#9 (Figures S2 and S3), from which dasc#1 [4’-(asc-C7)-asc-C7] and dasc#4 [2’-(asc-C5)-asc-C5] represent the two most abundant components^33,37^. The homodimeric dasc#1 is constructed by connecting two identical monomeric building blocks of ascr#1 (asc-C7) via a 4’-ester linkage (Figure 1A), whereas another ascaroside dimer, dasc#4, carries two units of ascr#9 (asc-C5), which is instead linked at the 2’-position (Figure S2). However, how two simple ascarosides like ascr#1 and ascr#9 (Figure 1A) are biochemically processed to yield diverse dimeric ascarosides with different structures and activities is still mysterious. Here, we report a new gene, *Ppa-uar-12*, encoding the *Ppa*-UAR-12 enzyme which is capable of establishing a unique 4’-ester linkage for the biosynthesis of dasc#1.

**Figure 3.**
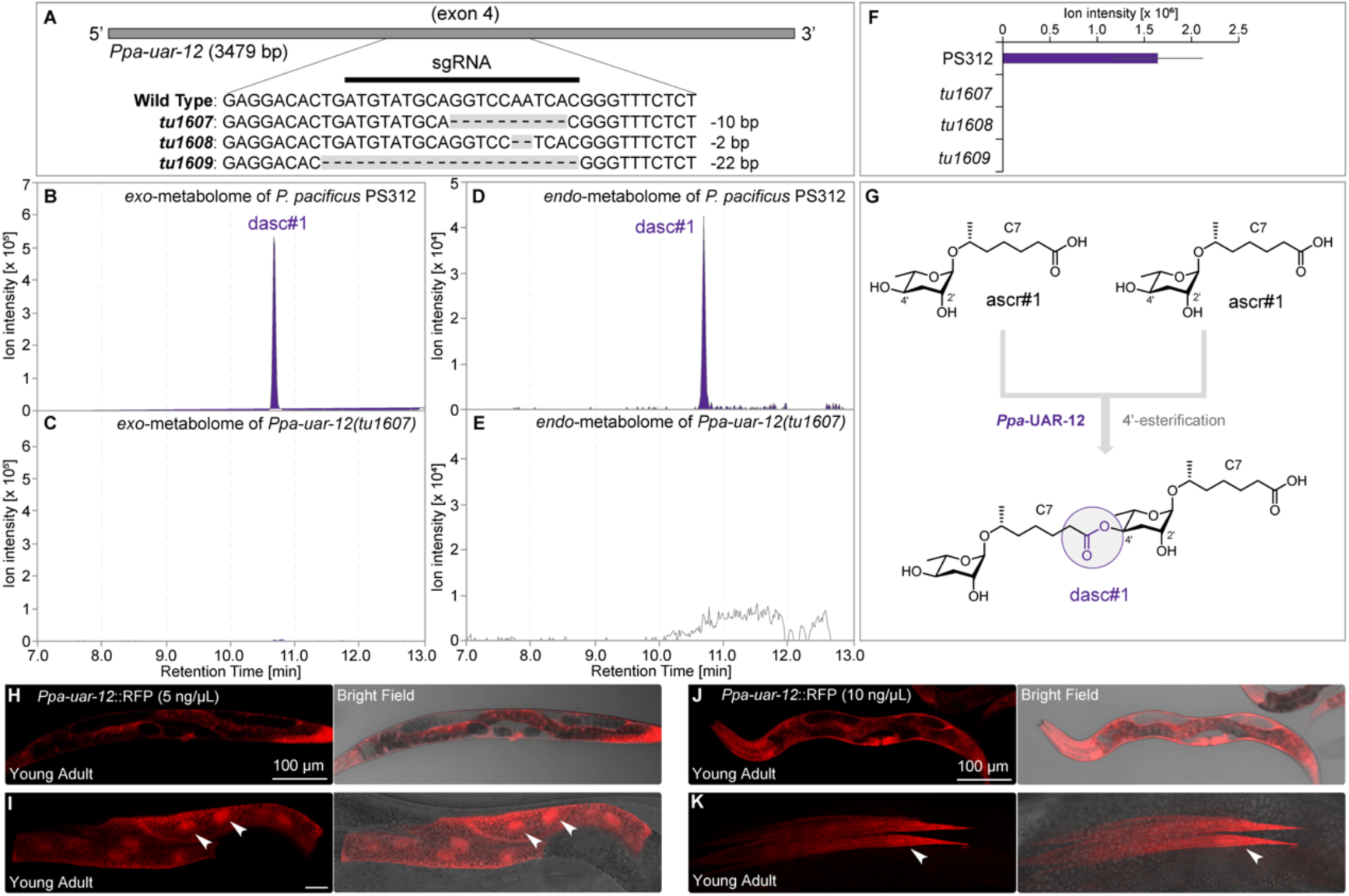
*Ppa*-UAR-12 regulates the biosynthesis of dimeric ascaroside dasc#1. (A) Three mutant alleles of *Ppa-uar-12* were generated by CRISPR/Cas9. (B-E) The production of dasc#1 in the *exo*-metabolomes (B and C) and *endo*-metabolomes (D and E) of both wild type *P. pacificus* PS312 and mutants were examined by LC-MS. (F) Ion abundances of dasc#1 in the *exo*-metabolomes of wild type *P. pacificus* PS312 and three mutants were quantitatively analysed. Three biological replicates were performed for each allele mutant strain. Error bars represent standard deviation (± SD). (G) Proposed biosynthetic pathway of dasc#1. (H-I) Confocal sections and overlaid Nomarski images of the reporter line expressing *Ppa-uar-12*::RFP at 5 ng/μl in the intestine. Scale bar is 100 μm. White arrows highlighted the nuclear localization of *Ppa-uar-12*::RFP. (J-K) Confocal sections and overlaid Nomarski images of the reporter line expressing *Ppa-uar-12*::RFP at 10 ng/μl in the epidermal cells. Scale bar is 100 μm. White arrows highlighted the expression of *Ppa-uar-12*::RFP in an individual epidermal cell.

When analyzing the pheromone production in *Ppa-uar-12* mutants, we found that all the three alleles of *Ppa-uar-12* mutants (*tu1607*, *tu1608* and *tu1609*), carrying deletions of 10 bp, 2 bp and 22 bp in exon 4, respectively (Figure 3A), did not produce any trace of dasc#1 in their *exo*-metabolomes (Figures 3B, 3C, and 3F). In addition, no trace of dasc#1 could be detected in the corresponding *endo*-metabolome samples as well (Figures 3D and 3E), indicating that the biosynthesis of dasc#1 was completely abolished in *Ppa-uar-12* mutants. In contrast, the production of other types of pheromones from *Ppa-uar-12* mutants was not affected, strongly suggesting that *Ppa*-UAR-12 regulates the biosynthesis of dasc#1 and has high substrate-specificity on the biosynthetic precursor ascr#1. For example, when we examined dasc#4, a previously identified 2’-linked homodimeric ascaroside^37^, we found that dasc#4 was present in both the *exo*- and *endo*-metabolomes of *Ppa-uar-12* mutants (Figures S2A-S2E). These observations suggest that *Ppa*-UAR-12 is only involved in the biosynthesis of the 4’-linked but not the 2’-linked ascaroside dimer, exhibiting extremely high substrate-specificity (Figures 3G and S2F). This was further supported by the detection of the other two 2’-linked minor dimeric ascarosides dasc#6 and dasc#9 in *Ppa-uar-12* (Figure S3). In the future, more efforts will be required to search for the yet uncharacterized putative enzyme(s) for the biosynthesis of the 2’-linked dimers like dasc#4, dasc#6, and dasc#9. Such proteins could possibly be uncovered among the remaining untargeted UAR candidates (Figure S1). Taken together, these results demonstrate that *Ppa*-UAR-12 enables the connection of two monomeric building blocks of ascr#1 and regulates the biosynthesis of dasc#1 via the formation of a specific 4’-ester linkage (Figure 3G).

*P. pacificus* produced dasc#1 has previously been reported to modulate its mouth-form dimorphism by inducing the Eu mouth morph^33^. When we investigated mouth-form plasticity under the standard laboratory growth conditions, we found that all three *Ppa-uar-12* mutants defective in dasc#1 production were still capable of forming the Eu mouth-form with ratios similar to the *P. pacificus* wild type animals (Figure S4). Thus, the abolishment of dasc#1 did not influence the mouth-form dimorphism of *P. pacificus* indicating that pheromone signaling and dasc#1 alone cannot override other aspects of mouth-form regulation, that resulted in the predominance of the Eu phenotype under the standard laboratory growth conditions.

Dissecting the expression patterns of biosynthetic genes may provide crucial insights into the cellular localization and the identification of the specific tissues responsible for synthesizing target pheromones. To investigate the expression pattern of *Ppa-uar-12*, a plasmid carrying the coding region of RFP codon-optimized for *P. pacificus,* under the control of the *Ppa-uar-12* promoter was generated^14^, and stable transgenic reporter lines were obtained after microinjection. Confocal imaging revealed that the biosynthetic gene *Ppa-uar-12* was mainly expressed in the intestine of the worms (Figure 3H). Besides the nuclei of intestinal cells (Figure 3I), some weak RFP fluorescence was also observed in the epidermal cells (Figure 3H). When we subsequently generated transgenic lines by injecting higher concentrations of the target DNA with a concentration of 10 ng/μl after linearization, we could further confirm that *Ppa-uar-12* was indeed expressed in the epidermal cells of *P. pacificus* (Figures 3J and 3K). Together, we identified a new biosynthetic enzyme *Ppa*-UAR-12 from *P. pacificus* and revealed its cellular localization in the intestinal and epidermal cells for the biosynthesis of the dimeric ascaroside dasc#1.

### *Ppa*-UAR-6 is essential for the biosynthesis of part#9 and npar#1-3

In 2012, a group of NPAR-type pheromones including part#9 and npar#1-3 (Figure 1A) was identified from *P. pacificus*^33^. All of these paratosides harbor an unusual core structure of paratose probably derived from the common ascarylose sugar via 2’-epimerization^33,54^, and exhibit the ability to induce dauer formation^33,36^. So far, only some closely-related species of *P. pacificus* from the *pacificus*-clade were found to produce NPAR-type pheromones^37^. Also, we have not found any clues about the biosynthetic origins of these signaling molecules over the last decade. Here, we identify a new gene *Ppa-uar-6* in *P. pacificus*, which encodes the *Ppa*-UAR-6 enzyme that is essential for the biosynthesis of both part#9 and npar#1-3.

Three *Ppa-uar-6* mutant alleles were generated by CRISPR/Cas9 engineering, with *tu2012* and *tu2013* displaying insertions of 11 bp and 7 bp in exon 3, respectively, whereas the third allele *tu2011* carried a 5 bp deletion (Figure 4A). LC-MS profiling of the NPAR-type pheromones from *Ppa-uar-6* mutant-derived *exo*-metabolomes showed that part#9 and npar#1-3 were all lost (Figures 4B, 4C, and 4F). Similarly, part#9 and npar#1-3 were also fully abolished in the corresponding *endo*-metabolome samples (Figures 4D and 4E). These results represent the first evidence demonstrating that the biosynthesis of NPAR-type pheromones could be completely blocked by mutating a specific gene, in this case *Ppa-uar-6*. The production of other pheromones in the mutant worms was not severely affected, except for the accumulating amounts of ascr#9 (Figure S5). These observations suggest that ascr#9 may probably play a role as a biosynthetic precursor of part#9 and npar#1-3. Therefore, we hypothesized that *Ppa*-UAR-6 creates an ester link on ascr#9, resulting in the production of “nasc#9” intermediates, by connecting tRNA metabolism-derived t^6^A^33,55,56^ to the C5 fatty acid side chain (Figure 4G). However, as “nasc#9” has not been detected in *P. pacificus*, we hypothesize that following a series of biochemical transformations including 2’-epimerization, hydrolysis of ribofuranose and glycosylation with xylose, the potential intermediate precursor of “nasc#9” is probably completely transformed to yield npar#1. Further cleavage of the xylose sugar from npar#1 by a yet unidentified enzyme will give rise to npar#2. Supposedly, npar#2 is further hydroxylated to form npar#3 which might be finally degraded by removing the group of adenosine-based threonine to synthesize the end product part#9 (Figure 4G). While the discovery of *Ppa-uar-6* supports a pathway suggesting part#9 is likely derived from ascr#9 via a series of streamlined biochemical transformations of npar#1-3 downstream of the peroxisomal *β*-oxidation pathway (Figures 4G and 5), the confirmation of these hypotheses awaits future analysis.

**Figure 4.**
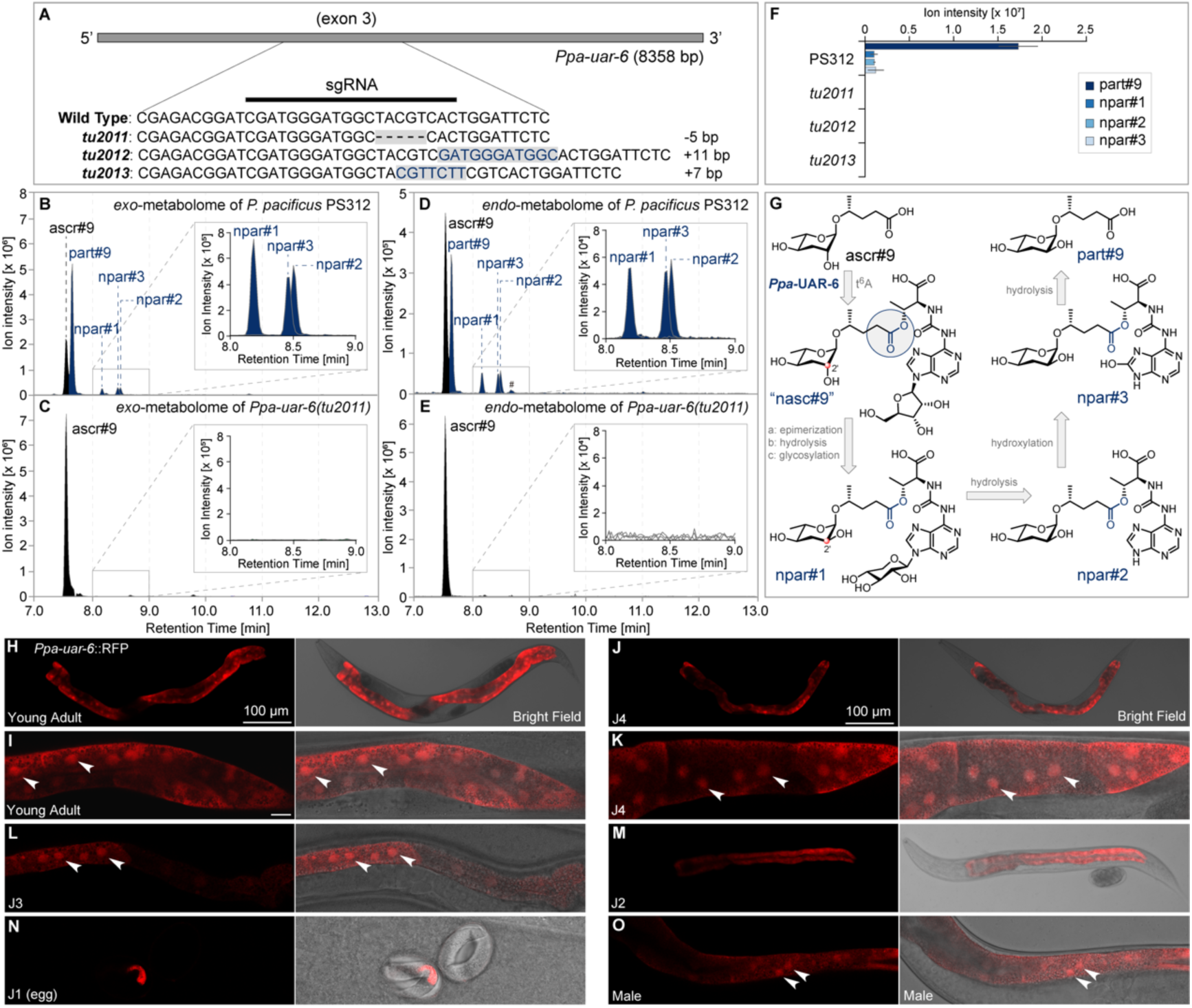
Biosynthetic investigation of NPAR-type pheromones. (A) Three mutant alleles of *Ppa-uar-6* were generated by CRISPR/Cas9. (B-E) The production of NPAR-type pheromones in the *exo*-metabolomes (B and C) and *endo*-metabolomes (D and E) of both wild type *P. pacificus* PS312 and mutants were examined by LC-MS. The pound sign (#) denotes non-ascarosides. Inserted ion chromatograms of npar#1-3 were magnified 10 times. (F) Ion abundances of NPAR-type pheromones in the *exo*-metabolomes of wild type *P. pacificus* PS312 and mutants were quantitatively analysed. Three biological replicates were performed for each mutant allele. Error bars represent standard deviation (± SD). (G) Proposed biosynthetic pathway of NPAR-type pheromones. (H-N) Section and Nomarski images of a transgenic worm in all developmental stages (H and I, Young adult; J and K, J4 larval stage; L, J3 larval stage; M, J2 larval stage; N, J1 larval stage) expressing *Ppa-uar-6*::RFP. Scale bar is 100 μm. White arrows highlighted the nuclear localization of *Ppa-uar-6*::RFP. (O) Section and Nomarski images of a male worm expressing *Ppa-uar-6*::RFP in the nucleus of intestinal cells. Scale bar is 100 μm. White arrows highlighted the nuclear localization of *Ppa-uar-6*::RFP.

**Figure 5.**
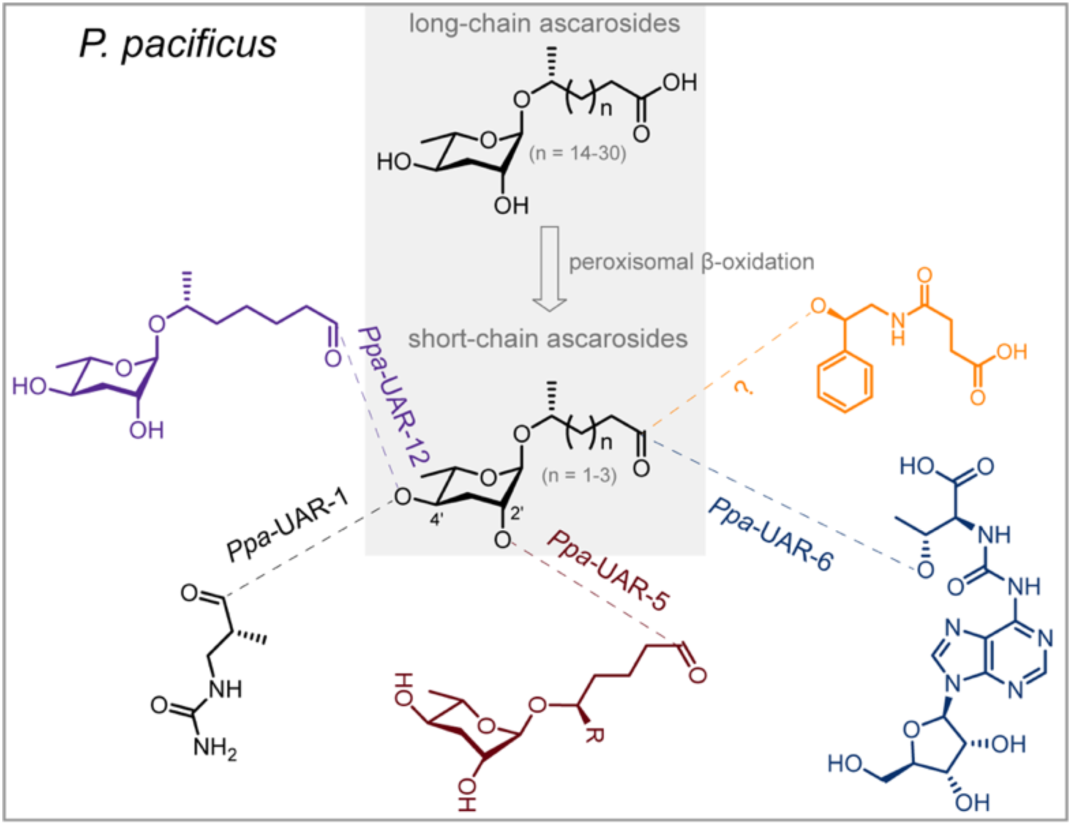
A group of UAR enzymes regulates the biosynthesis of modular signaling molecules from *P. pacificus*. *Ppa*-UAR-1 was previously found to specifically link ureidoisobutyric acid to the 4’-position of simple ascarosides for the biosynthesis of UBAS pheromones^49^. In this study, *Ppa*-UAR-5, *Ppa*-UAR-12 and *Ppa*-UAR-6 were discovered to be involved in the biosynthesis of UBAS, DASC, and NPAR-type pheromones, respectively. The UAR enzyme required for the biosynthesis of PASC ascarosides is yet uncharacterized.

Finally, we generated *Ppa-uar-6*::RFP transgenic lines through microinjection. We found the expression of *Ppa-uar-6* in the intestine of young adult hermaphrodites (Figures 4H and 4I). Notably, earlier developmental stages from J1 to J4 worms (Figures 4J-4N) together with males (Figure 4O) were also shown to have an identical intestinal expression pattern. Thus, the intestine of *P. pacificus* serves as the primary site for the biosynthesis of NPAR-type pheromones. Taken together, our work indicates that the complex *P. pacificus* pheromones discussed above are synthesized and pooled in the intestinal or epidermal cells.

## DISCUSSION

The biosynthesis of plant and microbe-derived secondary metabolites has been intensively studied using many different bacteria and plant species^57–65^. In contrast, the biosynthesis of nematode pheromones is still challenging^66^ and has thus far only been investigated in *C. elegans*^50–51^ and *C. briggsae*^67^, together with some preliminary studies in *P. pacificus*^49^. Here, we seek to investigate the biosynthetic origins of four types of modular signaling molecules in *P. pacificus*, each of which exhibits considerable complex chemical structures (Figure 1A). Previously, only one gene, *Ppa-uar-1*, had been discovered to be involved in the biosynthesis of UBAS ascarosides from *P. pacificus*^49^. In this study, LC-MS analysis of CRISPR/Cas9-generated mutants of 20 homologs of *Ppa-uar-1* facilitated the discovery of three carboxylesterase proteins *Ppa*-UAR-5, *Ppa*-UAR-12 and *Ppa*-UAR-6, which function downstream of the peroxisomal *β*-oxidation pathway to regulate the biosynthesis of UBAS, DASC and NPAR-type pheromones, respectively (Figure 5). These continuous efforts provide a step forward in uncovering the delicate biosynthetic regulatory mechanisms for each type of modular signaling molecules in *P. pacificus*.

In the future, the application of a work-flow similar to the one used here—genome-wide bioinformatics of genes encoding certain enzyme classes, followed by the systematic generation of gene knockouts, and subsequent LC-MS analysis of mutants—will possibly accelerate the discovery of biosynthetic genes for modular signaling molecules from other *Pristionchus* and more distantly related nematode species. Given that the CRISPR/Cas9 technique is widely applicable to knock out genes in *Pristionchus* species (e.g., *P. mayeri*)^68^, it will be increasingly feasible to investigate the detailed biosynthetic mechanisms of modular ascarosides in other nematodes. For example, it is very likely that *Ppa*-UAR-5 is also involved in the biosynthesis of the recently identified UPAS-type pheromones (Figure S6), considering that UPAS pheromones share very similar structures with UBAS^37^. Similarly, the recent discovery of novel ascarosides from other *Caenorhabditis* species like *C. briggsae*^53,54,69–71^, also offers great opportunities to explore their detailed biosynthetic mechanisms^32,51,67^. Therefore, the work-flow described in this study highlights a promising strategy to identify enzymes involved in the regulation of other complex pheromones in nematodes and potentially beyond.

Similar to the discovery of the *uar* genes in *P. pacificus*, more than 40 homologs of *Ppa-uar-1* were identified in the genome of *C. elegans*, which are referred to as *Cel-cest* genes. Indeed, five important candidate genes were ascertained to regulate the biosynthesis of different types of modular ascarosides, through the formation of an ester or amide bond^32,50,51^. For example, the *C. elegans* sex pheromone ascr#8^72^ is formed by CEST-2.2 through the introduction of a folate metabolic pathway-derived intermediate to the terminal carboxyl group of ascr#7 (asc-△C7)^51^. Thus, it will be highly interesting to study the biosynthetic mechanisms and capacities of UAR and CEST proteins in a broader evolutionary context.

Along with the CEST enzymes from *C. elegans*^50,51^, all the newly identified proteins *Ppa*-UAR-5, *Ppa*-UAR-6, and *Ppa*-UAR-12, as well as *Ppa*-UAR-1, exhibit high specificity for their substrates. For example, the esterase proteins *Ppa*-UAR-1 and *Ppa*-UAR-5 exhibit high substrate-specificity on ascr#9 and ubas#3, respectively, by selectively targeting their 4’- and 2’-position for the biochemical reactions (Figure 2G). Similarly, *Ppa*-UAR-12 establishes a 4’-but not a 2’-linkage for the biosynthesis of dimeric ascarosides. However, to test the selectivity and specificity of these enzymes in more detail will require additional efforts to purify carboxylesterase proteins from heterogeneous system to reconstitute the biosynthesis of pheromones *in vitro*.

Based on our experiments and other studies from different laboratories^44,50,51,73^, the confirmed expressions of the carboxylesterase genes in the intestine and in several epidermal cells further support the notion that these excretory tissues serve as the primary sites for the biosynthesis of nematode signaling molecules. However, the nuclear localization of the *uar* genes in the intestine is puzzling. While we speculate that this might have to do with the endo-duplication of the genome in intestinal cells, future studies using translational reporter lines may help to uncover their exact roles in pheromone biosynthesis.

Lastly, it is important to note that many biosynthetic steps for the generation of modular signaling molecules in *P. pacificus* remain unclear. Like *C. elegans*, the biosynthetic pathways are not fully understood in *P. pacificus*. For example, many other biosynthetic components are likely required for *Ppa*-UAR-6 to function as a multi-step biosynthetic pathway (Figure 4G). One hypothesis would be that additional carboxylesterases are involved in these processes. Indeed, prioritizing *Ppa*-UAR proteins based on sequence similarity to *Ppa*-UAR-1 might not be a perfect predictor of protein function. Thus, the remaining *uar* genes should be knocked out by CRISPR/Cas9 to generate more mutants in order to search for other unidentified biosynthetic genes of part#9 and npar#1-3 (Figures S1 and 4G). In addition to UAR enzymes, it is currently unknown if other classes of enzymes are potentially involved in the biosynthesis of NPAR-type pheromones. For example, the mechanism by which the 2’-epimerization of part#9 is formed will be an interesting question to be addressed, with a potential epimerase taking part in this reaction.

In summary, using a work-flow including i) bioinformatic tools for genome-wide carboxylesterase identification, ii) CRISPR/Cas9 gene knockouts and iii) chemical analysis of multiple mutant lines, three key biosynthetic genes *Ppa-uar-5*, *Ppa-uar-12*, and *Ppa-uar-6* were discovered in *P. pacificus*. The identified carboxylesterases are involved in the biosynthesis of three types of modular signaling molecules, including UBAS, DASC and NPAR-type pheromones. This discovery largely illuminates the biosynthetic mechanisms of the complex, modular signaling molecules in *P. pacificus* and will be helpful for uncovering their biological activities, most of which remain currently unknown.

### Limitations of the study

This study identified three new carboxylesterase enzymes from *P. pacificus*, which separately regulated the biosynthesis of three types of modular pheromones. However, the precise mechanisms of these biosynthetic pathways remain to be characterized. We failed to express carboxylesterase genes in heterologous system for the purpose of purifying proteins for the *in vitro* biochemical test. In the future, more efforts will be required to reconstitute the biosynthesis of pheromones *in vitro*.

## RESOURCE AVAILABILITY

### Lead contact

Requests for additional information on data, resources, and reagents should be directed to and will be fulfilled by the lead contact, Chuanfu Dong.

### Materials availability

All materials are available as indicted in the key resources table. All materials generated in this work are available from the lead contact upon request.

### Data and code availability

- All data are publicly available as of the date of publication, and software used for this work is indicated within the key resources table and method details.
- This paper does not report original code.
- Any additional information required to reanalyze the data reported in this paper is available from the lead contact upon request.

## ACKNOWLEDGMENTS

We thank Dr. Frank C. Schroeder for valuable discussions about the biosynthetic pathway of NPAR-type pheromones. We also thank Dr. Ziduan Han and Dr. Shuai Sun for their assistance with microinjection. LC-MS data and confocal microscopic images were collected by using Waters LCMS, Agilent LC-HR(ESI)-qTOF-MS and Zeiss LSM 980. Mutants were generated by using the CRISPR/Cas9 genome editing microinjection system. All the instruments were accessed from the Instrumentation and Service Center for Science and Technology (ISCST), Beijing Normal University, Zhuhai. We are grateful to Dr. Wu Wen, Dr. Yuan Gao, Dr. Xianxin Dong and Dr. Pingzhou Du at ISCST for technical supports and data acquisition. This work was funded by the Max Planck Society, National Natural Science Foundation of China (Grants NO. 22107011, 22407016 & 22477010) and Department of Education of Guangdong Province (Grant NO. 2021KQNCX269).

## AUTHOR CONTRIBUTIONS

P.Z., P.T., and W.X. performed the experiments. P.Z., P.T., and W.X. analyzed the data. S.Y., H.W. and T.L. assisted with microinjection and worm culture. R.J.S. and C.D. supervised the project. P.Z., P.T., W.X., R.J.S. and C.D. wrote the manuscript and all authors commented. R.J.S. and C.D. provided the funding. P.Z., P.T., and W.X. contributed equally.

## DECLARATION OF INTERESTS

The authors declare no competing interests.

## SUPPLEMENTAL INFORMATION

Document S1. Figures S1-S6 and Tables S1-S4.

## STAR★METHODS

### KEY RESOURCES TABLE

| REAGENT<br>RESOURCE or | SOURCE | IDENTIFIER |
| --- | --- | --- |
| Nematode strains |  |  |
| *Pristionchus pacificus* | Sommer<sup>1</sup> | PS312 |
| *Ppa-uar-1* | This study | *tu1948, tu1949, tu1950* |
| *Ppa-uar-2* | This study | *tu2022, tu2023, tu2024* |
| *Ppa-uar-3* | This study | *tu2014, tu2015, tu2016* |
| *Ppa-uar-4* | This study | *tu2068, tu2069, tu2070* |
| *Ppa-uar-5* | This study | *tu2019, tu2020, tu2021* |
| *Ppa-uar-6* | This study | *tu2011, tu2012, tu2013* |
| *Ppa-uar-7* | This study | *tu1993, tu1994, tu1995* |
| *Ppa-uar-8* | This study | *tu1934, tu1935, tu1936* |
| *Ppa-uar-9* | This study | *tu2017, tu2018* |
| *Ppa-uar-10* | This study | *tu1919, tu1920, tu1921* |
| *Ppa-uar-11* | This study | *tu1991, tu1992* |
| *Ppa-uar-12* | This study | *tu1607, tu1608, tu1609* |
| *Ppa-uar-13* | This study | *tu1563, tu1564* |
| *Ppa-uar-14* | This study | *tu1937, tu1938, tu1939* |
| *Ppa-uar-15* | This study | *tu1930, tu1931, tu1932* |
| *Ppa-uar-16* | This study | *tu1975, tu1976, tu1977* |
| *Ppa-uar-17* | This study | *tu1911, tu1912, tu1913* |
| *Ppa-uar-27* | This study | *tu2071, tu2072, tu2073* |
| *Ppa-uar-28* | This study | *tu1972, tu1973, tu1974* |
| *Ppa-uar-29* | This study | *tu1924, tu1925, tu1926* |
| Chemicals |  |  |
| ascr#9 | Bose et al.<sup>33</sup> | N/A |
| ascr#12 | Bose et al.<sup>33</sup> | N/A |
| ascr#1 | Bose et al.<sup>33</sup> | N/A |
| ubas#1 | Bose et al.<sup>33</sup> | N/A |
| ubas#2 | Bose et al.<sup>33</sup> | N/A |
| ubas#3 | Falcke et al.<sup>49</sup> | N/A |
| ubas#4 | Falcke et al.<sup>49</sup> | N/A |
| ubas#12 | Falcke et al.<sup>49</sup> | N/A |
| dasc#1 | Bose et al.<sup>33</sup> | N/A |
| dasc#4 | Dong et al.<sup>37</sup> | N/A |
| dasc#6 | Dong et al.<sup>37</sup> | N/A |
| dasc#9 | Dong et al.<sup>37</sup> | N/A |
| pasc#1 | Bose et al.<sup>33</sup> | N/A |
| pasc#9 | Bose et al.<sup>33</sup> | N/A |
| pasc#12 | Bose et al.<sup>33</sup> | N/A |
| part#9 | Bose et al.<sup>33</sup> | N/A |
| npar#1 | Bose et al.<sup>33</sup> | N/A |
| npar#2 | Bose et al.<sup>33</sup> | N/A |
| npar#3 | Yim et al.<sup>36</sup> | N/A |
| Oligonucleotides |  |  |
| For details of sgRNA and primers used in this study, see Table S3. | This study | N/A |
| Software and algorithms |  |  |
| SnapGene (version 8.0) | GSL Biotech | https://www.snapgene.com/ |
| ChemDraw (version 14.0) | PerkinElmer, Revvity | https://revvitysignals.com/products/research/chemdraw |
| MassHunter Quantitative Analysis (version 10.0) | Agilent | https://www.agilent.com.cn/zhcn/product/software-informatics/mass-spectrometry-software/dataanalysis/quantitative-analysis |
| R (version 4.3.1) | Comprehensive R Archive Network (CRAN) | https://www.R-project.org |
| Other |  |  |
| LC-MS | Agilent | 1290 Infinity II HPLC instrument coupled to 6546 LC/Q-TOF mass spectrometer |
| Fermenter | ZhiChu | ZQZY-CF8V |
| Microinjection system | Eppendorf | https://www.eppendorf.com/cn-zh/ |

### EXPERIMENTAL MODEL AND STUDY PARTICIPANT DETAILS

#### Nematode culture

All of mutant nematodes used in this study were listed in the key resources table and Table S1. Protocols for nematode growth medium (NGM) agar plates and maintenance of nematodes were described previously^37^. All strains were maintained at 20 °C on NGM agar plates seeded with *E. coli* OP50.

#### Method details

##### Reagents

Purified or synthetic ascarosides were used as standard controls for chemical analyses^33,36,37,49^. This study did not generate new chemicals or reagents.

##### Nomenclature of ascarosides from P. pacificus

*P. pacificus* produced ascarosides or paratosides were named using four letter codes “SMIDs” (<u>S</u>mall <u>M</u>olecule <u>ID</u>entifiers), for example, “nasc#9” and ubas#1^33^. The SMID database (www.smid-db.org) is maintained by Frank C. Schroeder and Lukas Mueller in collaboration with WormBase (www.wormbase.org). Structure-based abbreviations “(head group-)asc-(ω)(Δ)C#(terminal group)” were also used to describe common ascarosides, NPAR and UBAS-type ascarosides. For those DASC ascarosides consisting of two simple ascaroside units, the first ascaroside was placed in parentheses to differentiate it from the second ascaroside. For example, 4’-linked dimeric ascaroside dasc#1 was named as dasc#1 [4’-(asc-C7)-asc-C7]; 2’-linked dimeric ascaroside dasc#4 was named as dasc#4 [2’-(asc-C5)-asc-C5].

##### Protein sequence alignment and phylogenetic reconstruction

The amino acid sequence of *Ppa-*UAR-1 annotated by Falcke et al.^49^ was used for BLASTP search (*e*-value < 0.00001) against *P. pacificus* protein annotations (version 3) on www.pristionchus.org. Seventy-five protein sequences were obtained as the top BLASTP hits (Table S4). These sequences were aligned using the MUSCLE program (version 3.8.31)^74^. The resulting multiple sequence alignment was used for a maximum likelihood tree reconstruction using the phangorn package in R (version 4.3.1, model = “WAG+G(4)+I”, optNni = TRUE)^75^. The best tree model was determined using the pml_bb function. 100 bootstrap replicates were calculated for the midpoint rooted tree and nodes with the standard support values above 50 were visualized on the generated tree. R version 4.3.1 (www.R-project.org) was used to analyze the phylogenetic relationship of 76 carboxylesterase proteins.

##### CRISPR/Cas9-mediated knockout of twenty esterase genes

CRISPR/Cas9 knockouts were generated using previously published protocols in *P. pacificus*^13,14^. CRISPR RNAs (crRNAs) were designed to target the first, second, third or fourth exon of the gene. crRNAs and tracrRNA (Cat. No. 1072534) were synthesized by Integrated DNA Technologies (IDT). crRNAs were fused to tracrRNA (IDT) at 95 °C for 5 min, after which the Cas9 endonuclease (Cat. No. 1081058) purchased from IDT was further added. The CRISPR/Cas9 complex was prepared by mixing 0.5 mg/ml Cas9 nuclease, 0.1 mg/ml tracrRNA, and 0.056 mg/ml crRNA in the TE buffer followed by a 10 min incubation at 37 °C. The *Ppa-eft-3*::RFP plasmid (20 ng/μl) was added after the incubation as a co-injection marker^14^. Specific crRNA and primers applied for knocking out each candidate gene were listed in Table S3. Microinjections were performed following standard practice using an Eppendorf microinjection system. Injected P0s were removed ca. 16 hours post injection. Eggs from each injected P0 animal were allowed to hatch and further transferred onto single NGM agar plates. These F1 animals were genotyped via Sanger sequencing to establish heterozygous null mutant worms. A similar protocol was used to screen homozygous worms in the subsequent F2 generation.

##### Reporter line of Ppa-uar-12

To construct the transcriptional reporter line for *Ppa-uar-12*, a 3.0 kb *Ppa-uar-12* promoter fragment was amplified from *P. pacificus* genomic DNA using 2 × Phanta Flash Master Mix (Vazyme, P520). This fragment was cloned into the *Ppa-eft-3*::RFP plasmid vector (a gift from Ziduan Han, Northwest A&F University, China) upstream of RFP to substitute the *Ppa-eft-3* promoter using ClonExpress Ultra One Step Cloning Kit (Vazyme, C115) through homologous recombination. DNAs were prepared from multiple independent isolates, then verified by restriction digestion and sequencing. Plasmid DNA and genomic DNA were cut by PstI and injected as two fragments into the gonads of young adult *P. pacificus* worms at concentrations of 5 ng/μl and 30 ng/μl, respectively, to establish intergenerationally inherited worms that could consistently express RFP fluorescence. Normally, constructs were injected at 5 ng/ml or 10 ng/ml, along with *Ppa-eft-3*::RFP plasmid as marker. The expression pattern was confirmed in more than three independent lines. Single worms were imaged using a Zeiss LSM 980 confocal microscope. Stable lines of *Ppa-uar-12* were imaged and properly frozen at −80 °C for further usage. Worms recovered from −80 °C were confirmed to consistently express RFP fluorescence.

##### Reporter line of Ppa-uar-6

Reporter lines of *Ppa-uar-6* were constructed using the identical method as described for *Ppa-uar-12*. Constructs were injected at 10 ng/ml along with *Ppa-eft-3*::RFP plasmid as marker. The expression pattern was confirmed in more than three independent lines. Single worms were imaged using a Zeiss LSM 980 confocal microscope. Stable lines of *Ppa-uar-6* were also imaged and frozen at −80 °C for further usage. Worms recovered from −80 °C were confirmed to consistently express RFP fluorescence.

##### Preparation of bacterial food source for growing nematodes in liquid culture

LB medium (4 l) inoculated with a single colony of *E. coli* OP50 was incubated in a shaker at 37 °C with 170 rpm. After one night, *E. coli* OP50 pellets were collected by centrifugation (Beckman Coulter, Avanti JE 369003, JA10 rotor, 5000 rpm, 10 min, 4 °C). The supernatants were removed and resulted *E. coli* OP50 pellets were transferred into 50 ml falcon tubes for further centrifugation (Cence, H2050R, Swing rotor 0302891036, 4000 rpm, 10 min, 4 °C). *E. coli* OP50 pellets were equally split into six sterilized plastic tubes (50 ml) and stored in the refrigerator at 4 °C for further usage. All experiments were performed in a clean bench to avoid any potential contamination. Each tube of *E. coli* OP50 pellet was diluted by adding 10 ml M9 buffer. On the first day, 2.5 ml *E. coli* OP50 pellets were added into the nematode liquid cultures (50 ml) as their bacterial food source.

##### Collection of exo-metabolome (culture medium) and endo-metabolome (worm pellets) from mutant worms grown in 50 ml liquid culture

Bleached worms were initially cultured on five NGM plates (9 cm diameter) that were coated with 700 μl *E. coli* OP50. Using 6 ml M9 buffer to wash worms into a sterilized glass flask containing 50 ml S-medium. On the first day, prepared *E. coli* OP50 was diluted in 10 ml M9 buffer, and 2.5 ml of diluted *E. coli* OP50 pellets were added into each flask as bacterial food source. Worm cultures were incubated at 22 °C and 140 rpm for one week. Worms were harvested through centrifugation (Eppendorf, Centrifuge 5810R, FA-45-6-30 rotor, 8000 rpm, 10 min, 4 °C) to collect the culture medium (*exo*-metabolome) as well as the worm pellets (*endo*-metabolome). The resulting *exo*-metabolome samples were frozen at −80°C for one night and further dried into solid powder by lyophilizer. The solid powder was then extracted three times with 100 ml methanol each. Crude extracts were combined, filtered and dried at 40 °C under reduced pressure to remove organic solvents. Samples were finally dissolved in 1 ml methanol and 100 μl aliquots were applied for LC-MS analysis. The *endo*-metabolome samples were also frozen at −80°C for one night and further dried by lyophilization into a solid powder. The dried powder was extracted three times by adding 5 ml methanol each time. Crude extracts were combined, filtered and dried at 40 °C under reduced pressure to remove methanol. Samples were finally dissolved in 0.5 ml methanol and 100 μl aliquots were applied for LC-MS analysis. All 57 mutant strains were successfully grown in S-medium for chemical analysis. *Exo*-metabolome and *endo*-metabolome samples of wild type *P. pacificus* PS312 were prepared as standard controls. Three biological replicates were performed for each strain.

##### Chemical analysis of pheromones in exo-metabolomes and endo-metabolomes

*Exo*-metabolome and *endo*-metabolome samples were separately dissolved in 1 ml and 0.5 ml methanol. Samples were further filtered and 100 µl aliquots were applied for LC-MS analysis. LC-HR(ESI^-/+^)-MS analysis was performed using an Agilent 1290 Infinity II Series HPLC system connected to an Agilent 6546 high resolution qTOF spectrometer equipped with an electrospray ionization operating in negative ion mode. The HPLC instrument was equipped with an Agilent Eclipse XDB-C18 column (250 × 3 mm, 5 µm particle diameter). A mixture of water (H_2_O + 0.1% Formic acid) and acetonitrile (ACN + 0.1% Formic acid) was used to elute the LC-MS system with a flow rate of 0.4 ml/min (an injection volume of 2 µl). The LC-MS system was equilibrated with 3% acetonitrile for 3 min, followed by gradient elution (3%-100% acetonitrile) over 15 min, and finally washed with 100% acetonitrile for 2 min, after which the acetonitrile content of the solvent was decreased to 3% within the next 2 min. After every third sample, a blank MeOH run was performed using the same method. The *exo*- and *endo*-metabolomes of mutants were analyzed by LC-MS under the condition of Electrospray ionization (ESI) with Agilent 6546 high resolution qTOF system operating with capillary voltage 3500 V, dry gas flow of 12.0 l/min and dry temperature 275 °C. LC-MS analysis was conducted using the single MS mode by scanning from m/z 50 to 1700. Agilent Qualitative Analysis 10.0 software was used to calibrate, process and analyze the mass data of mutants and wild type *P. pacificus* PS312.

##### Mouth-form phenotyping

The mouth-form dimorphism of *tu1607*, *tu1608* and *tu1609* mutants were screened by a Zeiss LSM 980 microscope using standard protocols^33^. Wild type *P. pacificus* PS312 was used as control. All nematodes grown on NGM agar plates were kept in non-starved conditions and crowding of populations was properly avoided. Fresh *E. coli* OP50 culture was applied as food source. Young adult worms were picked for mouth-form screening (Figure S4). Intermediate mouth-form were not observed for any strains.

#### Quantification and statistical analysis

At least three biological replicates for all the experiments indicated in this study were performed. All the experiments of LC-MS analysis shown in each figure were conducted using the same mass spectrometer and method to ensure accurate data comparison. No samples or data points were excluded from the reported analyses. The quantification and statistical details of experiments can be found in the figure legends.

## Notes

### Competing Interest Statement

The authors have declared no competing interest.

