## Supplemental Information for "Biosynthesis of modular signaling molecules requires functional diversification of carboxylesterases in *Pristionchus pacificus*"

| Contents | | pages |
|---|---|---|
| **Supplementary Figures** | | |
| Figure S1 | Phylogenetic relationship of carboxylesterase enzymes from *P. pacificus*. | 3 |
| Figure S2 | The production of dimeric ascaroside dasc#4 in *Ppa-uar-12* mutants. | 4 |
| Figure S3 | The production of minor dimeric ascarosides dasc#6 [2’-(asc-C6)-asc-C5] and dasc#9 [2’-(asc-C7)-asc-C5] in *Ppa-uar-12* mutants. | 5 |
| Figure S4 | Confocal images of mouth-form of both wild type and *Ppa-uar-12* mutants. | 6 |
| Figure S5 | Quantitative analysis of the production of ascr#9 (asc-C5) in both wild type and *Ppa-uar-6* mutants. | 7 |
| Figure S6 | Proposed biosynthetic pathways of UBAS and UPAS-type pheromones. | 8 |
| **Supplementary Tables** | | |
| Table S1 | List of carboxylesterase genes were knocked out by CRISPR/Cas9 for biosynthetic investigation. | 9 |
| Table S2 | List of the remaining carboxylesterase genes were not mutated for biosynthetic investigation. | 10 |
| Table S3 | Primers and sgRNA were used for knocking out carboxylesterase genes. | 11 |
| Table S4 | Amino acid sequences of carboxylesterases from *P. pacificus*. | 12 |

#### Supplementary Figures

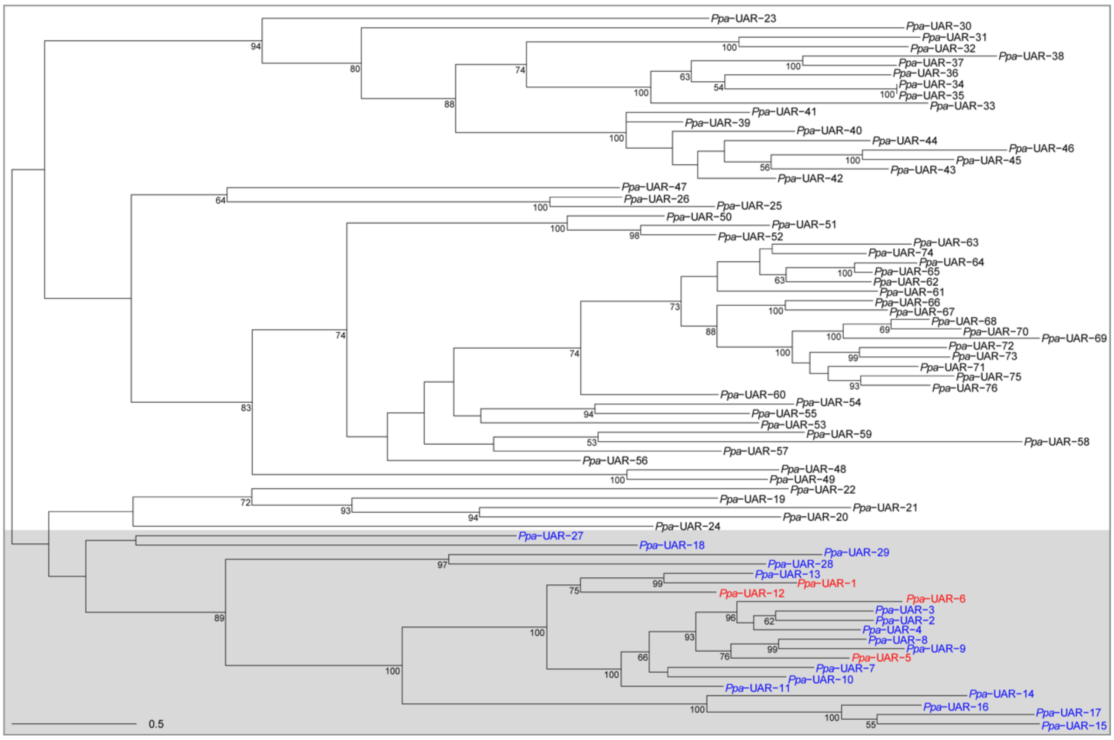


**Figure S1. Phylogenetic relationship of carboxylesterase enzymes from *P. pacificus*.** 75 homologous proteins of *Ppa*-UAR-1 were discovered from *P. pacificus* (Table S4). Their amino acid sequences were aligned with *Ppa*-UAR-1, from which twenty proteins (marked in blue) share high similarities with *Ppa*-UAR-1 in the phylogenetic tree (four functional biosynthetic enzymes were marked in red). These 20 encoding genes were selectively knocked out by CRISPR/Cas9.

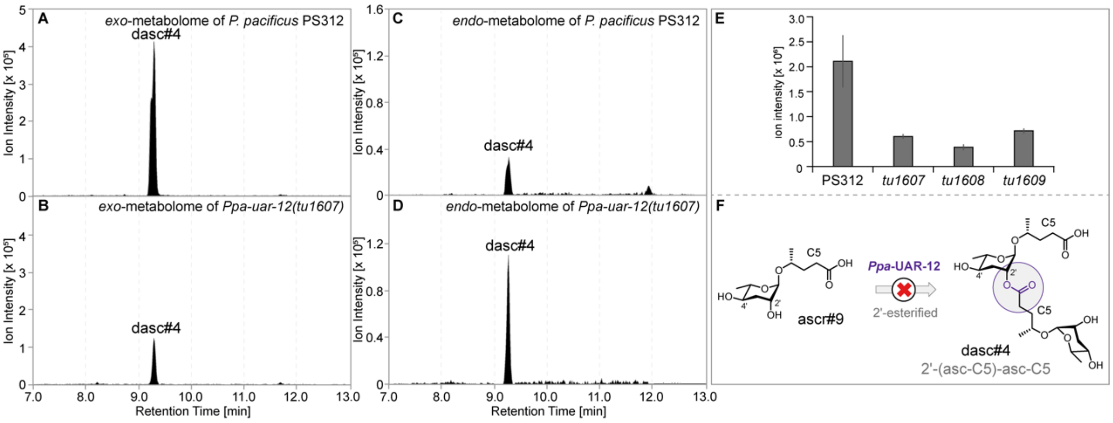


**Figure S2. The production of dimeric ascaroside dasc#4 in *Ppa-uar-12* mutants.** (**A-D**) The production of dasc#4 [2'-(asc-C5)-asc-C5] in the *exo*-metabolomes (**A** and **B**) and *endo*-metabolomes (**C** and **D**) of both wild type *P. pacificus* PS312 and mutants were examined by LC-MS. (**E**) Ion abundances of dasc#4 in the *exo*-metabolomes of wild type *P. pacificus* PS312 and three mutants were quantitatively analysed. Three biological replicates were performed for each allele mutant strain. Error bars represent standard deviation (± SD). (**F**) Carboxylesterase *Ppa*-UAR-12 is not involved in the biosynthesis of the 2'-linked dimeric ascaroside dasc#4.

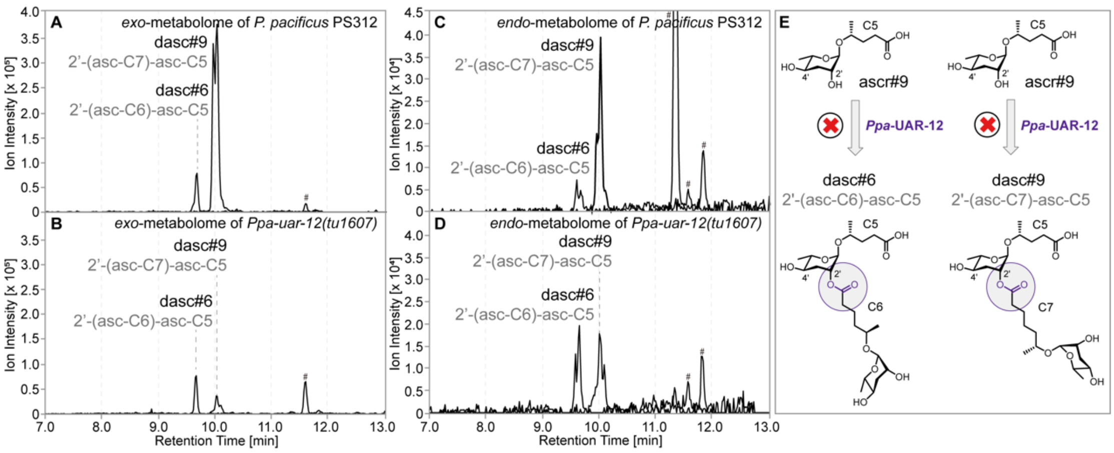


**Figure S3. The production of minor dimeric ascarosides dasc#6 [2'-(asc-C6)-asc-C5] and dasc#9 [2'-(asc-C7)-asc-C5] in *Ppa-uar-12* mutants.** (**A**-**D**) The production of the 2'-linked dimeric ascarosides dasc#6 and dasc#9 in the *exo*-metabolomes (**A** and **B**) and *endo*-metabolomes (**C** and **D**) of both wild type *P. pacificus* PS312 and mutants were examined by LC-MS. The pound signs (#) denote non-ascarosides. (**E**) Carboxylesterase *Ppa*-UAR-12 is not involved in the biosynthesis of the 2'-linked dimeric ascarosides dasc#6 and dasc#9.

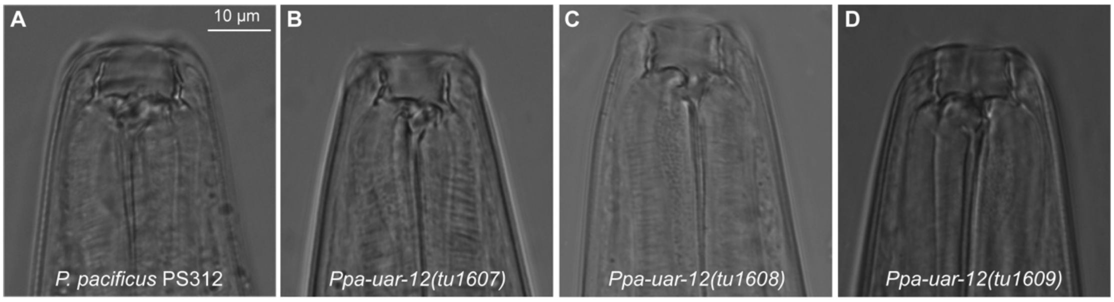


**Figure S4. Confocal images of mouth-form of both wild type and *Ppa-uar-12* mutants.** (**A**-**D**) The mouth-form of wild type *P. pacificus* PS312 (**A**) and *Ppa-uar-12* mutant animals (**B**: *tu1607*; **C**: *tu1608*; **D**: *tu1069*) was Eu under normal culture conditions. Scale bar is 10 μm.

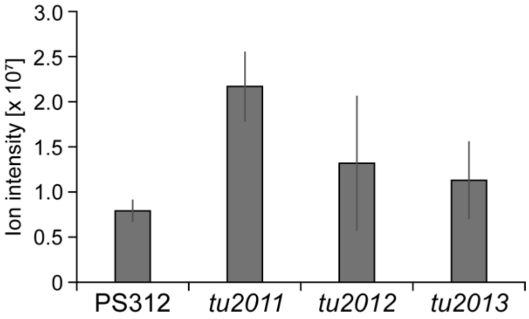


**Figure S5. Quantitative analysis of the production of ascr#9 (asc-C5) in both wild type and *Ppa-uar-6* mutants**. Three biological replicates were performed for both wild type *P. pacificus* PS312 and *Ppa-uar-6* mutants. Error bars represent standard deviation (± SD).

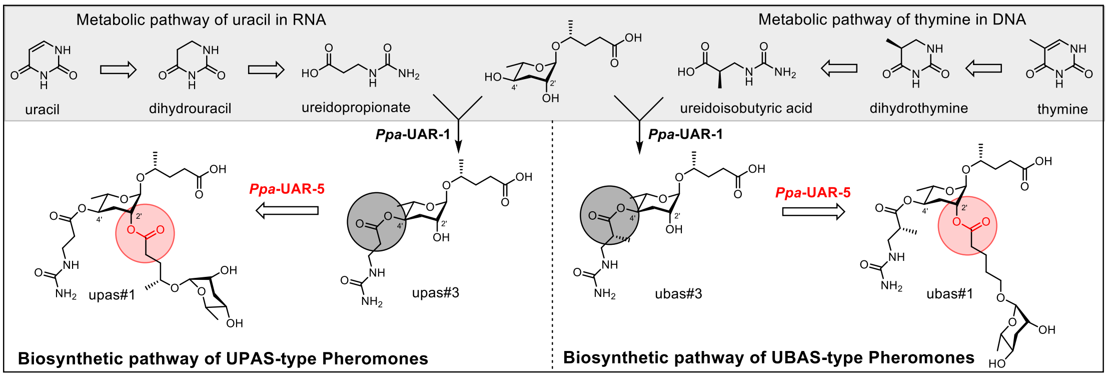


**Figure S6. Proposed biosynthetic pathways of UBAS and UPAS-type pheromones.** *Ppa*-UAR-1 functions upstream of *Ppa*-UAR-5 to synthesize intermediate precursor upas#3 or ubas#3, by linking pyrimidine (uracil and thymine) metabolism-derived intermediates (ureidopropionate and ureidoisobutyric acid) to the 4'-position of ascr#9. *Ppa*-UAR-5 further integrates another building block of ascr#9 or oscr#9 to the 2'-position of upas#3 or ubas#3 to finally synthesize upas#1 or ubas#1.

### Supplementary Tables

#### Table S1. List of carboxylesterase genes were knocked out by CRISPR/Cas9 for biosynthetic investigation.

| Name | Esterase or hydrolase genes | Allele 1 | Allele 2 | Allele 3 |
|---|---|---|---|---|
| ***Ppa-uar-1*** | Iso_D.223.8 | *tu1948* (8 bp deletion, exon 2) | *tu1949* (7 bp insertion and 4 SNPs, exon 2) | *tu1950* (29 bp insertion and 1 SNP, exon 2) |
| ***Ppa-uar-2*** | ppa_stranded_DN9008_c0_g1_i1 | *tu2022* (10 bp deletion and 1 SNP, exon 4) | *tu2023* (19 bp insertion, exon 4) | *tu2024* (8 bp deletion, exon 4) |
| ***Ppa-uar-3*** | ppa_stranded_DN30926_c0_g3_i15 | *tu2014* (1 bp insertion and 4 SNPs, exon 4) | *tu2015* (4 bp insertion, exon 4) | *tu2016* (5 bp deletion, exon 4) |
| ***Ppa-uar-4*** | Iso_D.224.1 | *tu2068* (4 bp deletion, exon 4) | *tu2069* (11 bp deletion, exon 4) | *tu2070* (20 bp deletion, exon 4) |
| ***Ppa-uar-5*** | PPA25673 | *tu2019* (68 bp insertion and 5 SNPs, exon 4) | *tu2020* (11 bp deletion, exon 4) | *tu2021* (4 bp deletion, exon 4) |
| ***Ppa-uar-6*** | ppa_stranded_DN30149_c0_g1_i4 | *tu2011* (5 bp deletion, exon 3) | *tu2012* (11 bp insertion, exon 3) | *tu2013* (7 bp insertion, exon 3) |
| ***Ppa-uar-7*** | ppa_stranded_DN20512_c0_g1_i2 | *tu1993* (14 bp deletion, exon 3) | *tu1994* (4 bp deletion, exon 3, “variant#1”) | *tu1995* (4 bp deletion, exon 3, “variant#2”) |
| ***Ppa-uar-8*** | ppa_stranded_DN28083_c0_g1_i4 | *tu1934* (1 bp deletion, exon 3) | *tu1935* (11 bp deletion and 2 SNPs, exon 3) | *tu1936* (41 bp insertion and 5 SNPs, exon 3) |
| ***Ppa-uar-9*** | PPA42050 | *tu2017* (20 bp deletion and 1 SNP, exon 7) | *tu2018* (4 bp deletion and 1 SNP, exon 7) | |
| ***Ppa-uar-10*** | Iso_D.1590.1 | *tu1919* (2 bp deletion, exon 3) | *tu1920* (26 bp insertion and 2 SNPs, exon 3) | *tu1921* (5 bp deletion, exon 3) |
| ***Ppa-uar-11*** | PPA17279 | *tu1991* (5 bp insertion, exon 5) | *tu1992* (1 bp deletion, exon 5) | |
| ***Ppa-uar-12*** | PPA21718 | *tu1607* (10 bp deletion, exon 4) | *tu1608* (2 bp deletion, exon 4) | *tu1609* (22 bp deletion, exon 4) |
| ***Ppa-uar-13*** | PPA21991 | *tu1563* (5 bp deletion, exon 3) | *tu1564* (55 bp deletion, exon 3) | |
| ***Ppa-uar-14*** | PPA39247 | *tu1937* (11 bp insertion and 6 SNPs, exon 4) | *tu1938* (4 bp deletion, exon 4) | *tu1939* (11 bp deletion and 8 SNPs, exon 4) |
| ***Ppa-uar-15*** | ppa_stranded_DN31291_c3_ g5_ i5 | *tu1930* (8 bp deletion, exon 3) | *tu1931* (1 bp deletion and 1 SNP, exon 3) | *tu1932* (1 bp insertion, exon 3) |
| ***Ppa-uar-16*** | ppa_stranded_DN29519_c1_g1_i1 | *tu1975* (11 bp deletion and 3 SNPs, exon 5) | *tu1976* (2 bp insertion and 9 SNPs, exon 5) | *tu1977* (14 bp deletion and 2 SNPs, exon 5) |
| ***Ppa-uar-17*** | PPA23744 | *tu1911* (31 bp insertion and 1 SNP, exon 1) | *tu1912* (10 bp deletion, exon 1) | *tu1913* (4 bp deletion, exon 1) |
| ***Ppa-uar-18*** | PPA23795 | Embryonic lethality (exon 1) | | |
| ***Ppa-uar-22*** | PPA23754 | In-frame mutation (exon 3) | | |
| ***Ppa-uar-24*** | PPA23779 | Embryonic lethality (exon 3) | | |
| ***Ppa-uar-27*** | ppa_stranded_DN23720_c0_g1_i2 | *tu2071* (28 bp deletion, exon 4) | *tu2072* (10 bp insertion and 5 SNPs, exon 4) | *tu2073* (7 bp deletion, exon 4) |
| ***Ppa-uar-28*** | ppa_stranded_DN25492_c0_g3_i2 | *tu1972* (5 bp deletion, exon 4) | *tu1973* (37 bp insertion and 5 SNPs, exon 4) | *tu1974* (8 bp deletion, exon 4) |
| ***Ppa-uar-29*** | Contig35-snapTAU.30 | *tu1924* (1 bp insertion and 1 SNP, exon 2) | *tu1925* (4 bp deletion, exon 2) | *tu1926* (39 bp deletion, exon 2) |

**Table S2. List of the remaining carboxylesterase genes were not mutated for biosynthetic investigation.**

| Name | Esterase or hydrolase genes | Name | Esterase or hydrolase genes | Name | Esterase or hydrolase genes |
|---|---|---|---|---|---|
| ***Ppa-uar-19*** | PPA15738 | ***Ppa-uar-41*** | ppa_stranded_DN16831_c0_g1_i1 | ***Ppa-uar-60*** | ppa_stranded_DN30573_c0_g1_i2 |
| ***Ppa-uar-20*** | ppa_stranded_DN26620_c0_g1_i2 | ***Ppa-uar-42*** | ppa_stranded_DN27477_c0_g1_i1 | ***Ppa-uar-61*** | PPA04333 |
| ***Ppa-uar-21*** | PPA25312 | ***Ppa-uar-43*** | ppa_stranded_DN18687_c0_g1_i1 | ***Ppa-uar-62*** | PPA37960 |
| ***Ppa-uar-22*** | PPA23754 | ***Ppa-uar-44*** | Contig58-snapTAU.64 | ***Ppa-uar-63*** | PPA39767 |
| ***Ppa-uar-23*** | ppa_stranded_DN25900_c0_g1_i1 | ***Ppa-uar-45*** | ppa_stranded_DN29180_c0_g1_i2 | ***Ppa-uar-64*** | PPA41660 |
| ***Ppa-uar-24*** | PPA23779 | ***Ppa-uar-46*** | PPA36339 | ***Ppa-uar-65*** | PPA10010 |
| ***Ppa-uar-25*** | Iso_D.2531.1 | ***Ppa-uar-47*** | ppa_stranded_DN28328_c1_g1_i1 | ***Ppa-uar-66*** | ppa_stranded_DN25758_c0_g1_i1 |
| ***Ppa-uar-26*** | ppa_stranded_DN13491_c0_g1_i1 | ***Ppa-uar-48*** | PPA35183 | ***Ppa-uar-67*** | ppa_stranded_DN9111_c0_g2_i1 |
| ***Ppa-uar-30*** | PPA39427 | ***Ppa-uar-49*** | PPA10555 | ***Ppa-uar-68*** | ppa_stranded_DN31013_c0_g1_i1 |
| ***Ppa-uar-31*** | Contig58-snapTAU.108 | ***Ppa-uar-50*** | PPA03821 | ***Ppa-uar-69*** | PPA03767 |
| ***Ppa-uar-32*** | PPA02785 | ***Ppa-uar-51*** | PPA13063 | ***Ppa-uar-70*** | PPA41108 |
| ***Ppa-uar-33*** | PPA07019 | ***Ppa-uar-52*** | PPA22191 | ***Ppa-uar-71*** | PPA36944 |
| ***Ppa-uar-34*** | PPA40913 | ***Ppa-uar-53*** | PPA28522 | ***Ppa-uar-72*** | PPA34289 |
| ***Ppa-uar-35*** | PPA40915 | ***Ppa-uar-54*** | PPA00898 | ***Ppa-uar-73*** | ppa_stranded_DN30123_c0_g1_i5 |
| ***Ppa-uar-36*** | Iso_D.10131.1 | ***Ppa-uar-55*** | Contig34-snap.12 | ***Ppa-uar-74*** | PPA40739 |
| ***Ppa-uar-37*** | ppa_stranded_DN18279_c0_g1_i1 | ***Ppa-uar-56*** | PPA22060 | ***Ppa-uar-75*** | PPA42366 |
| ***Ppa-uar-38*** | PPA10777 | ***Ppa-uar-57*** | PPA30526 | ***Ppa-uar-76*** | PPA26246 |
| ***Ppa-uar-39*** | Contig58-snapTAU.63 | ***Ppa-uar-58*** | PPA22915 | | |
| ***Ppa-uar-40*** | Iso_D.6227.1 | ***Ppa-uar-59*** | ppa_stranded_DN26305_c0_g1_i1 | | |

**Table S3. Primers and sgRNA were used for knocking out carboxylesterase genes**.

| Name | Esterase or hydrolase genes | Sequence of gRNA | Forward primer | Reverse primer |
|---|---|---|---|---|
| ***Ppa-uar-1*** | Iso_D.223.8 | CCATGGAATGGGACGCTAGA | GTGATCAGCACTGGATATGGTTCG | CGATTACCTCTCGACATCTCTCGC |
| ***Ppa-uar-2*** | ppa_stranded_DN9008_c0_g1_i1 | GTCTATATCCATGGAGGCTC | GTGCTCAAGTCGACTCATCATGGG | GAACGAAGCGCAGACTCTCCACG |
| ***Ppa-uar-3*** | ppa_stranded_DN30926_c0_g3_i15 | CCTGGGTCGGATACGTAACT | CATCGAGTACCAAGCAGGGAATGG | CTTTGTGCACATCACCTTGTCTCGG |
| ***Ppa-uar-4*** | Iso_D.224.1 | CCAGTGGTAGTGTACATCCA | TCAGAAGACTGCCTAACACTGAACG | AGGCAATACGTGCTCGTCACC |
| ***Ppa-uar-5*** | PPA25673 | CTACGTCACTGGACTGTCTG | CCAATCGTCCAAACGTCATACGGC | CTTGCCTCGCGTAGTTTGTCGC |
| ***Ppa-uar-6*** | ppa_stranded_DN30149_c0_g1_i4 | CGATGGGATGGCTACGTCAC | GAAGCGAGTAATGGATTCATCGG | GATAATTCACAACAAGCGTATCATCGG |
| ***Ppa-uar-7*** | ppa_stranded_DN20512_c0_g1_i2 | GGGCCATTCAGAGGACTGCC | GATAGTGAGGAGAATCTAGTCGAGG | GACGATAGTCCATTGTGACCAGG |
| ***Ppa-uar-8*** | ppa_stranded_DN28083_c0_g1_i4 | CAGGTTCCCTCACACTGGGA | CCTCATCAGCCCTGGAATCAC | CTGTCTGCACTCTGTTGACGTG |
| ***Ppa-uar-9*** | PPA42050 | GAACAAATCGCACGTAGTCG | AGCTACAATCGTGCTGGCTCTCG | CTGAGGCTGCATTGATGATCTCACTG |
| ***Ppa-uar-10*** | Iso_D.1590.1 | CCATTCTACTGGCGCGACGA | CTCGAATCTGTTCTTGCATGGTAC | GGAGATGCACTCATAGCAATGG |
| ***Ppa-uar-11*** | PPA17279 | CGCAAGGAGATTCACAATTT | GCCAATATTGCTATTCACGGTAAGC | CGATCCACTCATGGCTATCGC |
| ***Ppa-uar-12*** | PPA21718 | ATGTATGCAGGTCCAATCAC | CGGATCAATTCGTGGGTATGAATACAAGAAC | CGATAGGCTGTAGTGACCACGACG |
| ***Ppa-uar-13*** | PPA21991 | GCCTATCGCCTTGCAGCGTT | GGCATTTTCGAAGCTACGATGAAG | CCACTCGTTGACTCCTCTCTACTGTC |
| ***Ppa-uar-14*** | PPA39247 | GACTTCATTCCCACCAATTT | TACGTGAGCGCTGAATGCAGG | GACCCATTAGAGTGAGGGCATCTG |
| ***Ppa-uar-15*** | ppa_stranded_DN31291_c3_ g5_ i5 | TGCGATGGAAACTGCCTGAG | GGATGACTGGATAGCGAAGACC | TGCGAAGACGTTGATGTAGAGG |
| ***Ppa-uar-16*** | ppa_stranded_DN29519_c1_g1_i1 | CAACGGACGGTACAAGAGTT | CACTCGCCGATAAGAATGCTCC | GTTGAGTTGCTCATACAAGCTGCTG |
| ***Ppa-uar-17*** | PPA23744 | TTGATAGTTAACACCAGCTA | CTTATCTCGAGACTGGTTATCAAC | GTGGAGTTACTCATACAAGCTGC |
| ***Ppa-uar-18*** | PPA23795 | AAGCAACTTCAAATCTTCCG | CCTTCCTCTTTCCTCATCTC | CCAATTCTTGATGCCATCGAG |
| ***Ppa-uar-27*** | ppa_stranded_DN23720_c0_g1_i2 | GTCGCATCCAGAGAGAAGTC | GCAACGATTATGCAATCTTCAAGG | GATGCTCCACCGTACTGGAAC |
| ***Ppa-uar-28*** | ppa_stranded_DN25492_c0_g3_i2 | CGGACGGGCTACGGATTTCT | GGGACTGTGAGTCTCAAGATCAGG | GTTGCCACAGTACAGCGATGC |
| ***Ppa-uar-29*** | Contig35-snapTAU.30 | AAATCACGAGTAACTCTATT | TTGCATCCAGGTCAGTTACGAC | CATGAAGCTGCGATGTTCACATC |
| ***Ppa-uar-22*** | PPA23754 | ATGCATTCAAACTGGACTAC | CATTATGGAAGTGCTGTCTCATCC | GGACGAATATGTTCATGTAGAGGC |
| ***Ppa-uar-24*** | PPA23779 | CCCTGGTACGAATCCGTATT | GAACAACAACGACTCCGAGACC | CACCATGGAAGAATACTATCACTGG |

#### Table S4. Amino acid sequences of carboxylesterases from *P. pacificus*.

| Esterases | Amino acid sequence |
|---|---|
| ***Ppa*-UAR-1** | MRLLVLSVSLFFITASYSTSDYYFPVISTGYGSIRGYAFTAHDGTEAQIFKKIPFASAPIGDLRWRKPQPHRPWNGTLDGTFFGPACTQRTNKYDGPVTGFSEDCLHVNVYTSERCRESNSTCPVA<br>FIIHGGAGLYESTMKFPDEKLVRNFVSQGIVVVTTAYRLSAFGAMDLGDENALPANLGLHDIVAALNFTRNEIGHFGGSKEKDHSTRPVRGRSLRADDRVFTGDIEAGREKRLIDGIIVMSGNGGLE<br>FREKAVERSHSVAKQLNCTGTAREIVECMRLHDTESIVAASYKVNGPHILSQKAPFGITMSGELFPITNEKELREDPNPIRLMIGTTIEEIGGGVAGVDKINRILGIENGEECYDKYLNDVESGAFNP<br>GYIEASQEMIMTAHIYAKYVAQIGGEAYLYEYDYPVHGGHTDDAYHVLGIHEYEMDENERWLSRAYPRYFSNFIKGERLAKDWSKVTPLLMNYYSVNRSRTDGVFPHTKYGYKNNLVKYYDELVK<br>YDQVLSAAKMQAANAPIEYKSLNFDAGFQGFLESLSIFDAIIFVCVVLGVLSLLCCLCNCFTSLCCCCCRGGYSSI |
| ***Ppa*-UAR-2** | IFSRMINQCILLLIPFVSSLSKYPIAHTPYGSVRGYEFEASNGFVGEIYKKIPFAAPPIGNKRWKKPGPPEPWNHTIDGTFSGPACAQVDSSWAGYPTGISEDCLTLNVYTSKQCRESNAACAVVVYI<br>HGGSGLFSSSIHFPDDHLVSKYATEGVILVTISYRLGVFGVMALGDEHALPANLAVHDVVESLRFVQQTIHAFGGDKDQVSIMGHSIGSTIVLFLMYSPEVNKAGKPPLFSRAIAISASMNFENVEK<br>QVNRTHGVAAHLGCKGTAHEIITCLLPFSTEQMLRAASEVGGDALGFSATHLAGIVKAGELFPAHEMNEIRENQEDHMTEIAPAPPTKLMIGTQLNEFRQDEFLWWYEANVLNGTVIEAIDVLGVR<br>NYKDCADKYTDDVKSGRFKAEYDTLSQCLILTASLFASEQVRAGGEVYLYQYDYSNHSDHADDLYFLMGVHDYPMDDNEKWLSRVYPVYFTNFIKGLPLAPDWKPLDPKLMNYYSINKSFEEGI<br>VPQMKFGYHQPITDYYLDLQKFDKNLSNVKQLKATAPIMFNSLVLPSSESVNLRDIIFYSVFTVSIVLFIMGVIQCSRNSRRRAGYIVLKSDV |
| ***Ppa*-UAR-3** | MIFEAKESKQKLPMVSIPGFLSSTWSSVMQYLLSLSLRRAHNTSSLVHRMLLFLLFFLPIASALIKYPIAHTPYGSVRGIEYQAGNGFVGEIFKKVPYASPPVVERRWKKPEPPEPWNYTIDGTFAG<br>PACAQVNSTWVGYVTGLSEDCLTLNIFTSKKCRESNGTCPVVVYIHGGSALYSSALHFTDESQVSNFPRQGVILVTMEYRVGIFGVAALGDEHALPANLAVHDVIQSIRFVRSSIHAFGGDKEQITV<br>MGHSIGAGMVLLLLFSPAINKAGEPPLFARAVASSGVMNFEPVVKQVNRTHGVATKLGCEGTAQEIVTCMRGLSTDEIIRAAAEVGGNDLFSETHLSGMTIAGELFPFRDVREMRKQQKDHMNGI<br>LPAPPTKLLIGTLFNEFRPSAYFFYKETGFLDLSLNEAIDSLGVRNYNECMTKYFDDWKTGQFKPGYDTLSQGLFVTAAIFGSAQVRAGGEVYLYSLDYPKNTNHADDMYILMGIHEYSMDSNQE<br>WLSRVYPVYFTNFIKGLPLAPDWEPLNPELMNYYSINKSFTDGIVPHMKLGYHQPILEYYANLTTFDENLTKSIQMVANAPVQFKSAPIPRVEPVNTPSIPSDIIVTSLYFALMLLIVVIMCQCCRSAR<br>KGAEDIPLKMVA |
| ***Ppa*-UAR-4** | CHARVNAKRRLIVSVQCLIMSTLARSTLLLLFVLPIASSLTNYPIVTTSYGAVRGYEFEASNGFVGEIFKKIPFATPPIGVQRWKKPVPPERWNYTVDGTFAGPACAQINSTWAGYVTGQSEDCLTLN<br>VYTSRNCRVSNGSCPVVVYIHGGSALFSSSIHFPDDVLISKYATKGIVLVTISYRVGVFGVMALGDEHVLPANLAIHDVVESLRFLRTEVKAFGGNKDQITVMGHSTGASAALILVFSPTINKAGEPP<br>LFARAISVSGTMNFEDEPKQVKRSHAVATHLGCEGTAQEIVDCLSLISTDDLLRAATEVGGGDLLSVNHLTGITLAGELMPVRDAYELRQQQEEHMKGIAPARPTSLLMGTLVNEFRGDVLQAYGQ<br>EEEDRSLHEAIDTLGVRNLQECTHKYYDDVRSGYFKPGYDKLSQCVYLTASTFGATQARAGAEVYLYQVDYPKHTNHADDLYYLMGVHNFPMDENEEWINHVYPEYITNFIRGLPLAPDWEPLD<br>PNLMNYYSVNKCFTEGIVPQTKFAYHQPISDYYAELMRYDQNLTKIERLAASAPIQYKLLALPSSDSFTLRDLFIYSTIFGLILLIILTTCKCGRNKRPTNEQIPLKPKEFI |
| ***Ppa*-UAR-5** | MFTALKSRANMCYSVFKITFGFLQNEGFFNESSSIGVLIPSSDDRFLKCASRSFTHCQQQKRTQSDHFDSRNCLMKILLVLLIFPIASSLSNFPIVQTSYGSVRGYEYRTKSGFVGEIFKKIPFAAPP<br>TSARRWTKPEPPEPWNHTIDGTFFGPGCAQVRSSWKGYVTGLSEDCLTLNVYTSRECRLSNSTCPVVVYVHGGSALFSGALMFPDEVLATNYARQGVILITIAYRVGVFGVMALGDENALPANL<br>AMHDIVQSLRFVRQEIHAFGGDTDKVTLMGHSTGATIVVALVFSPNVNKAGETPLFSRAIAMSSTMNLESEEKQVNRSHVLAAMLGCAGTAREIVDCLRPLSTDAIISAAQAVGGPDMFSPTHLSG<br>ITLAGELMPIHNMRDLRENQEEHMRSLGKASPTKLLLGSILKEFKMGRGVVDQELDRLNVGTNQALDVLGVRNKEECLMKYFNDLNAGAFNSVYDTLSQAFLMTTAWFASAQARTSAEDENED<br>WLSRVYPVYFTNFIKGVPLAPDWSPLDPDSMNYYSINKSISDGVVPRMKYGYHQHLNDYYQEMMKFDETVTNNKRVKVNTPIEYKNAVLLSENSSNFRDIVYYCSIICVISFTAFKMFDLLKRRRT<br>HSRLNGPSLEKLFEQIDTMG |
| ***Ppa*-UAR-6** | MQEIFFLLLGLPLVACLTKFPIVRTSYGAVRGYEYEASNGFIGEIYKKIPFAAPPIGARRWKKPAPPEAWNYTLDGTFAGPACAQIETDRWDGYVTGFSENCLTLNIYTSKECRQSNASCPVVVYLH<br>GGSLIYSSAVHYPDDTLVVNYPTQGVIMVTIAYRVGVFGVMALGDENALPANLAIHDIMESLRFLRREVHVFGGDKDQISVMGHSIGANLALFMTYSPAVNKAEEPPLFARAISISASMNLFTEEKQ<br>VSRSHAVATHLGVMSGHSCGNCAVPASIQHR |
| ***Ppa*-UAR-7** | VEMTILFLISLVLPITSSLSNFPIVHTSYGSVRGYEYQAKNGFVAEIFKKIPYASPPIDARRWKKPAPPEAWNYTIDGTFFGPGCAQYPPRFITGHSEDCLALNIYTSRACRESKASCPVAFFIHGGNA<br>VTGGTMNYPDEALVSNFASQGIVLVTMDYRLGIFGVLALGDENVVPANLAIHDVMESLRFTRKEIHNFGGDKEQITVMGHSTGATMVIVIAFSKGINQSEESSLFSRAIAMSPSAVFNKEEAQVAMS<br>HFAANALGCSGSAQEIIDCLTLMSTDEIIDEMTEKVESEAEVRDIALAGELLPFRNVKELHENQEPTTLLIGSTLDELGYAPRSPKDHACFMIGARNIDQCLEKYERDVALGTFEPVYNVVSQAYFML<br>NWQIAKAQVKAGGEVYLYQYDYPLNAVHAQDLFYILGSHPRELDANEEWLSHVYPIYFSNFIKGLSLAPDWKPLDADLMNYYSINKSLTDGVEPEMKIGYHQTIADYYNNMIEFDENLTKIIQMQLN<br>APIQYKDLILQATDTFSITDIILYSAVLAFIFFAVFKLNRFCLNTRSDEKSPLILAHGRNKRS |
| ***Ppa*-UAR-8** | FRQPPHCLMINAAAICLTFLHVASSLSTYPIVRTSYGAVRGYEYQAKNGFVGEVYKKIPFAAPPTGERRWNKPAPHQPWNHTLDGTFFGPACAQVPSHWEGYVTGTSEDCLTLNVYTSRTCRES<br>NATCPVVVFFHGGSTLTGGTTVFPDETLVTNFAKRGIIMVTAAYRLGVFGVMTLGDEHVLPANLALHDAVASLRFVRSEIHSFGGDRDQVTIMGQSAGATIVVALVFSPFINLVDNDILFSRVIAMSS<br>CLILKSEESQVNISHAVAAKLGCSGTAREIIDCMRPLSTDEIINAASAVGGPDMSSSVSHLRDITLAGDFFPFLSVRELRKNQKRHMEQLSSPTKLLLGTMLNEFKVEPLVSNINASVNEVTEVLGLF<br>NNEECVQKYNSDVASGAFDPGYDTLSQALFVSNALYASVHARTGAETYLYQYEYAAYSMHGYDMYYVLGSHAHPMDENEEWLSRVYPLYFSNFIRGMPPAPDWERFEPDLMNYYSINKSFAVG<br>ASPKMKYGYHQNLNEYYSGMVQFDQNLTDFKEMVCIVCQRQTIVEILFAPIQYKKLVLHSTTPINIRHLLFYGTLVGIFIIAMHRSRNFRRITDVFVL |
| ***Ppa*-UAR-9** | MCSLVCLLLLPKALSLITYPIVHTSYGTVRGYEYQAMNGFVGEIYKNIPNSSHILRKIPYASPPIGRRLTALHALRFHHTGRAMSQAPPRIASHSTFTRQESVGSGNTVFPDDKLVTNYPKQGVIMVT<br>VSYRLGVLGVMTLGDENALPANLALHGDHNGPVNWSYNRAGSRFLSSQSPLFSRAIAMSAAMNLEREAKQVNKSHVVAAKLGCYGTAQEIIDCMRLLSTDEIINAASAVGGSDMFSVSHLSGITL |

| | |
|---|---|
| | AGELLPFHSVRELRENQKKHMALFSSPTKLLLGTMLTEFNAGPLRGLVNNINTGINQVTDVLGVINKEECVQKYYNDVDSGMFDPGYDTLSQTLFVTTLLFASSQAQTGAEVYLYQFDYPAHAWH<br>ADDVYYALGSQSHPVDENEEWLGRVYPVHFTNFIRSLPPAPDWELFDSELMNYYSINKSFSDGVSPGMRYGYHQDLTDYYDALVQFDENLTGFKQTVLNAPIQYKKTALYSSKPINIRDLFLIGIVA<br>GIVLIAMSSSNLIMYFLILLLGLPSLTHSCLRMREPQPGIPFVCPPFDDYTDCNDDYQDLCMPATRSTTEVTCGTKPYTFFAVIGVMAILGSEPYSVACVKKPV |
| ***Ppa*-UAR-10** | MVTVAYRLGAFGAMSLGDENVLPSNLFLHDVLEAVHFTRREIHNFGGDKDQITIMGHSTGATMVMMLSLSPGINKDADETLFARSIAMSASPILKKKEKQVERSNAVASVLGCNGTAQEIVDCLLP<br>FSTDQIVQAAFEAGEDNAEKMLLDITMTGELVPIRSMRELSMNRMRSKLLLGTTQNELGFIDVTKETDYVNVIIGVQNEEECSQKYMEDRKSGKFDPGYNLISQAAFMLTWTIANGQAMNGGEVY<br>LYQFDHPTYSFHGDDTFYVVQSRENLAGENEEWLNRVYPQYFSNFIKGLPLPQDWKHVDPALMNYYSIYKSSDGVEPTMKLDYHKTIADYYTDMIKFDENLTKGRDNQMLMFLFLFFVFPFVSS<br>LSNFPIIRTSYGSVRGYEYRAKNGFIAEIFKKIPFAAPPIGAKRWKKPVPPEAWNYTIDGTFFGPACAQHPSHWAGYVTGFSEDCLTLNIYTSTKCREAGSSCPVVVYIHGGFALFDGTMMFPDETI<br>VTNFASKGIIMVTVAYRLGVFGVLALGNENVLPANIAIHDVLESIRFIRKEIHNFGGDKDQISVMGHSTGSTVIFILAFSPGIENTKESPLFARAIAMSGTVNFNSEEKQIERSHAVTTRLGCSGTAQEII<br>DCMMLYNTCEIIQAAKDEEGDMKFMSPTQLADITLAGELMQFRSGKELSENKKPIKLMVGTIMNEFDPPRDFYRNASIETKQKLMCSKKNKVCEILGLRNYEECSDKYYNDCISGKFEPGMTVAS<br>QSTFMTNWLFANAHAMSGGEAYLYQFDYIAHAQHTDDAYYVMGFHEHPKDVNEEWLSRVYPAYFANFIKGVPLAPDWKPVDPELMNYYSVNKSFTDGVSPEMKLGYHRILSDYYLGLVEFDEKI<br>SKLKQTVFNAPVQYKDLSILSDEPFEFRDLIVYLTIIGIILFIAFKLYQCYDERQNRGERSPFLQA |
| ***Ppa*-UAR-11** | MVLLSLLLLLPIASSLSNFPIVSTSYGKVRGYEYRAKNGFVGEIFKFWCRMCATFFSVERICDRILRGLSESEHLHFERSTSSCPVILYIHGGVALFDGTMMFADEALITNFASVIMVTTEYRLGVFGV<br>MALGDENVLPANIAIHDVLEALRFTRKEIHNFGGDKDQISIMGHSTGATIVLTMSFSPGINKPGEPPLFARAIAMSGSTNYEDEEKQVKRSHDVAAKLGCEGSAQEIVDCMLPISTEKILDVAFKTGG<br>LDVFSKTQLNDLTMAGELMPIENVKELREKQKVTKLMLGTTLYEMEMKPFNTSSTVNNEVNLILGVKNEEECTQKYFADKKSGKFVSQYSGQSQAFFVTKWLFSEAQTKAGGEVYLYQYDYPAH<br>AIHTDDVSYVMGFHDHAKDVNEEWLSRTYPIYFANFAKGLPPAPDWTPVVPELMNYYSVNKSFTDGVSPEMKTGYHRDIIDYYKGIVQYDDNLSKIKKKLLNAPVQYKELIVLTTERVNIRDILFYSII<br>LGVLLFLVFKSYQCWRERQRSAEESPLLIDRLNNNE |
| ***Ppa*-UAR-12** | MLFILLFFLQALSSSSISNFPIVSTSFGSIRGYEYKNDAAFTAQIFKKLPFASPPIGDLRWRKPQSPSKWNTTIDGTFFGPACAQRTLMYAGPITGFSEDCLHVNVYTNEKCRTSSSCSVIFIIHGGLGI<br>YESTMKFPDETLVTNFVSQDIVVVTTAYRVGAFGIMALGDENALPANLAMHDVLAALKFTRDEIHAFGGDKESITIMGHSTGAQIALNFAFSPGISPPGVPRLFSSVISMSGPLGLEKEEKQVERSH<br>LVANLLKCSGTAREIIACMKKKTTDEILDAAFDVQEGDIFNERGLIGLMMGGELMPIRSARELRENNQPIRMMIGTTLYETKNFGLYNSTEVDEDKVNRVLGIDNDEECFEKYLDDTKSREFVTEHS<br>TDSQALFMTAYSLAKEQSESGGEVRMAKEMLPMPGIEPGPPG |
| ***Ppa*-UAR-13** | MLLSLSLFTFVLLNLPSVFAEFPVVTTSYGTLRGYVYTADDGTEAHIFKKIPFAEPPIGELRWKKPVATEKWNGTRDSTFFGPACAQKTIVYKGPVTGFSEDCLHLNVYTSQKCRESKTCNVVFVIH<br>GGVGIFEATMKFPDEALVRNFISQDIVVVTTAYRLAAFGAMSFGDENVLPSNLAMHDILEALRFTRKEIGNFGGDNEKVSDATMNKLSTYILITVLGHSEGGHYALMLAFSPGISKPGEKRLFESVISI<br>SGPSGLESQEQTVERSQRVAKELGCVGSAREIMDCFRRFDTQTIVNATFKVHGPNILSSQGPIGITMAGELFPLFKELRERRDPVRLMTGTTIYEMPGPSAKDKVNRVLGIVNREECYEKYTRDT<br>ESGEFTPGYNEVSQDIVMTAHLYAKYQAEIGGEAYLYEYDYPIHGDHTDDVYFLLGFHEYELDDNQKWLSRVYPRYFTNFIKGRRPAADWAQVTPRLMNYYSINKSLTDDVYPHTKYEYQNNIIQY<br>YDGIVKYDNLLFLVKTKVLSAPVQYKSLNFNGDFQDFFESLTMIDASMPSTVRSFSVLLGLVGVLSALPIESSTTFPTETTTITWVANEQIRKHINLINEFERIREEMNDASVESTTSSDSTMIPMADL<br>DDGIEAAQMLIHKFFAIKKDLETKKQEYAMMTTTTIATTTPGRNFIRKLFDKIKFFD |
| ***Ppa*-UAR-14** | YERIARNGKTIRIIKLDSNCVFILQSIPFAAPPIGELRWKLPQDPNKWDGILEGTKYSAACLSNHTDTQKMDEDCLYLNVYVSAECRGECPIILMFHGGAMNFNSAVYYNDDHLIENFASKGIMLVVP<br>AFRLGFMGIFNLGDDFIPTNLGLHDALHALQFVSKEAVNFGGNKDALTLMGHSYGGTIAGVLAYSSLRREKVNISRLIMMSPSFQFTSLQTVQTLSFRLVEKSGCSGRSSSLILSCLKSKDAAELLR<br>YQSEIQKEDESISWDAGFMFILMSSPLLPFSSFTQMFDLPPGVDVVIGSTAREMDVPVKLQTHPGDLVNAANLIELEKIHEILVENKNGTLHTPAIELIFLTVHKVVRSVIHGGNKAFVYSYQQPSSH<br>NYHSDDFSFIAGVSLFEQDNNEKDIAKFYPELFSNFTLIGIPSPDWMSMNESGSYFSIVVDANSKRTDPLQKCHDSRPKMLDGYERESINYWENEAPKIDIFEIGNSFEDIIKSKSITNVELSEKFES<br>CIDRQATNEYRGIYASN |
| ***Ppa*-UAR-15** | FSMIIRLIAALFIIGSAIAQDDWIAKTTYGSIQGYETTSNNGTRVKVFKAVPFAAPPVGDLRWKLPEWPKKWNDVKDGTKYSAACMSNSTTTKSPQEWVDEDCLYINVFAPAHCSAEHKCPIITFYH<br>GGGMNYDSAVYYDDKALIETYPANGVILIVPAFRLGFTGQFSLGDNQELVPSNLGVYDALHALKYVKQEAAAFGGDSDAITVMGHSGGGGLSILLAFSPVRKLQVSIDRFVMMSPGLAFDPPGNN<br>EKLSKEMVERAGCSSISKKGTLACMRKKSGPELLAIQRAMEEENAPSRSIFGGIVMCPPLFPFKSIPELLENPPKASILIGTTTKEMDLTPSENAMHGANVGTFIGAKNVKELDEIYDSMTRSDKNS<br>TFSCFQSIRNHFFQGTVKF |
| ***Ppa*-UAR-16** | FRMILRIFPILIVISLSIAEEDLIVNTSYGSIQGYESTSTDGTRVRIFKSVPFASPPIGDLRWKLPVQPKKWEGVKDGKKYSAACMSNSTTSKSPQPWVDEDCLYLNVFIHGECSASNKKCPIVVYYH<br>GGGMNFDSATYFNDTALKDTYASNGVMLVIPAFRLGFTGQFTLGDNEELVPSNLGVHDALHALKYVKKEAAAFGGDTDAITVMGHSFGGGLAIMLAFSPLRKLQVPIHQFIIMSPGFTFDPPETNE<br>KLSKEMVKRAGCSNMSKKLVFSCMKTKSGPELLAIQRTLEEENFPSVGATFGGVLLRAPLFPFDSISDVMNNPPNANILIGSTINEMDTLTNATGRTGALIGAKNVDELDQLYGNLSLSGKINLTHA<br>PQTESLYVSVYLTSKAVMENGFKAFLYSFDQPTHNHHTDDLSYLMGIHYFEKDDNEKEIAKFYPELFLNFTKFGTPSPNWEPTNHRSRHYSIKVDLPSGRMPEMHDFYEKETIDFWVQKAPKIDK<br>MITVTKAAGAKIEWEKITGEFRDLIIEEDSAIGKKNETIVPVVLIHSSYFPSFLMVCAVFLLGIILGRYLLGDSESHEERMWILGPDGRGMRIVSGKAPAYAEQMF |
| ***Ppa*-UAR-17** | MLTTNQTRPDNLISRLVINTIYRVFSSPRTQLIDKHYRGILFFSSPLVLFFALGVSVFSPNRYFRMVVRFIVALSLIGSVIGQDDLIVNTSYGSIQGYESTSSDGTKIRIFKSVPFASPPIGDLRWKLPIEP<br>KKWKGIKDGTKYSAACMSNSTTSKSPQPWVDEDCLYINVFIHGKCSKTNKCPVVVYYHGGAMNYDSATYFNDTALIETYASNGVMLVIPAFRLGFTGQFSLGDNQELIPNNLGIYDTLHALKYLKE<br>EADSFGGDPDGITVMGHSFGGVLSILLAVSPVRKHEVSINQFIMMSPGFYMDPPEINENLSMEMVKRAGCSNISKKLILSCMLKKTGPELLAIQRKMEEEDGSVVFGGAVMSPPLFPFKSLPELLK<br>NSPRANIMIGSTAKELDYPPKPDTPHGSNVGALVGAKNLDELDELYYNLTQSGKNNQTHSSDTQGLYLSVYIASRANMEKGNKVFLYSFDQSTHNMHTDDLSYLMGIHGFEKDENEKEIAKFYPQ<br>LFLNFTKYGEPSPNWKPMNLKDRYYSIEVNKTSGRQWDKISGEFRDLILEDDSNSIKNETFIPVVFVQMRMIILLTFILCLVATSIAQDDLTVKTSYGSILGYETLSSTGIRVKIFKSVPFASPPIGDLRW<br>KLPVKPKKWEGIKDGTKYSAACMSNSTTTSSPQLVRLSNRIHLLLYVLQWVDEDCLYLNLFIHGECSVRTFIIKQNKCPIVVFFHGGDMNYDSATYFNDTALIETYASNGVMLVIPAFRLGFTGQFSL<br>GDNQELIPSNLGVHDALQALKYLKDEADAFGGNSDAITLMGHSFGGALSVFLGFSPLRKLQVPIHQIALMSVGIDFRPADVVENLSKEMVKRAGCSNISKKLTLSCMLKKSGSELLAIQRRMEEEW |

| | |
|---|---|
| | FGGALFGGVVMSPPLLPFKSVPDMFKNAPNSNILVGGTIVEMDGPPNPLLPHGAGVGKLIGAKNVAELDELYYNLTQSGKTNQIHSPDSQNIYVGVYVTARAVMEKGNKAFVYSFDQRTHNIHTD<br>DLSYLMGIHGFEKDENEKEIAKFYPQLFINFTKYGVPSPNWKPMNVKDRYYSIEVNLTSGRHEILKPHFIQVYEFRMPEMRDYYEKETIDFWVRKAPKIDKMITVTRAAGAKMRWDKVTRGFRDLI<br>LEEDSTPAKNETLIPVALVHMSYFPSFLLICSVFLLGVLLGRFLLGDSTSQDERMWILGPDGRGIRIDPAKPPAYAEQMF |
| ***Ppa*-UAR-18** | MFYLLLLFFLPLSSSQNLEKSRSVWVEQGLVRGKIYKIGDKQLQIFRGIPYAEPPVGDLRFRKPTKKSRWHQELAAVEYGSPCLQFMDFHRNDRFASESMKRESEDCLYLNVFSPYDGDDESSLS<br>PILVWIHGGSFLAGSGDTGIDMEVVAKNIVFKGVTLVTLNYRLGPLGFMNYNMDGKVEGNFGIYDIIMALEWIQANIKQLNGDPSRITLMGESAGGAAVSLLSVSPLTRDLFHRSIVMSGSSAAGWA<br>VHTHGTPMWSIENMISYLRCERAISEEDANEIVGEEFSEEEIGRKHCNYQEEKIACLSPDMNTQEMLSCLTKSLNFSSSLFRRALATELGVSKVIVDGHLLSSSGADFISENARIPLLTGVAKSEWS<br>HKKPQFYNLPRLTNLTKEEVEQAVFKVIDTSFHQSVKDKLTNSTLLLLSNASFVRYMDDPTGQYATANVVTALQKVMEADIEFVAPCHREISAYQSKGIPVFAYSFDYLPESPIYEEEKKIFSLFGKQ<br>PIDIVRKDANFADHKLEAFHGLDHAYIFSQGYTSNFEIRPFTKRDDVMSKMLTNMWTNFVKTGDPSTARFKWPITTPEETQYVSIQLPPKVIQGDIHFPSPKFWNVEADLISRFVSKGTDRLTIDPAS<br>ELTNEERVELSAYRRAWWALWVLVGVLAFLLWSFVIYIALKKSRAPRSKPYDNIVVAAR |
| ***Ppa*-UAR-19** | MDCQEPSTSYCEYPVPPSEDEPEPEACRRVPRGSPARRSTVAVTIESIALLLALLVQIAHANDVIYLRDGSPLFGEELTAPNGKLVTQFFGIPFAEPPVGNLRFRKPVPKAPWRNILNATVLPNSCIQ<br>SLDTNFGEFFGATMWNANTKQSEDCLYLNVFVPGRLDQTKRLAVMVWVFGGGFWSGSSTLDVYDGKIFPTEENIILVTLNYRVSIFGFLYLGTPEAPGNMGLWDQHLALKWVHANIDQFGGDNT<br>RITLFGESAGAASVNMQMLSEKSTPYFHRAIIQSGSATAPWAMETTERTVVVYNVMKCEGEHTNMAFHEMDINKIYKCFMSASHETLRDNEWAPVREFVDFPWVPVVDGDFLIESAQTSLRNGH<br>FKTTQLLAGSNLDESIYFIVYQLGEIFPLKKFFTGRDFIQDRQTWIKGLMDLLPREMVRSKVAVSSIVHEYEPADLPVKPSDWINALDKALGDLQFTCNVNEMALAHTRHGGDTFYYYFSHRASMQ<br>TWPAWMGVLHGYEINFIFGEPYNRISYNYTREEQELSSRFMRYWANFARTGDPNKNEDGSYTTDPWPKYNAASMQYMNLTVESDYTAGAGRLGHGPRRKECQFWKAYLPNLMTALDEVELIE<br>SDEESSSSTTVPSTTVPVEPTVSSSTLPYPMAADTRSSEASLVTPIVVALSLYGLVLLTAVCLVTAPRFFNRRKTENLPECTCEEEESLGPTIYMTDIPDPIVHWKQEMHRWSQEYMLDWQRHFEQ<br>YKKYQNYRRRDSDEGTCDL |
| ***Ppa*-UAR-20** | MDRMIGNGIVLLLYSLLSPIYSRAILNDGFVVHTSLGTIRGVEQKFEDKVIGAYLGVPYGRPPTGNKRFQRPQMAERTSTDFVADKLARSCFFTQDHTFPDFPGAEMWNPPNSQDEDCLNMNIWV<br>PREHDGSVLVWIYGGGFFSGSPSLDLYDGSVLAALHNTIVVNINYRLGPFGFLYLGEESPVSGNMGLLDQQLALKWIHHNIAAFGGDPKRVTLFGESAGSASTTAHLIAPGSRHYFHKIIAKSGAIIN<br>SWATRSKQEMLDVSFNLVRRLNCSNSDHELILKCLAHIPSHIVQREADGITGGIHLPMSFAFVPVEEDNHFFKGNVFDKIRRRDFKKDVSALVGTVKDEGTYWLPYYMSDGQFGFKFNHTISPED<br>PFNSARINREQYKKSLEAFLPYFGNSPLVKHALMNAYQDVPDKQSFNKKDEEKLRDGVARFLGDYFFTCGLTQFADILAESINGPLYMYYFTMRSSANPWPRWMGVMHGYEIEYAFGVPLRRPQ<br>MYRKDRLEVEQSFSKKIMQMWINFANSGSPSSFWPRYNRIDKRSLELNENIVTGGYYVGVDVHGEGCRLINEGKTFSRAKDCRANRFDLPSSSSLSIGLSSITVVILSIINILRVL |
| ***Ppa*-UAR-21** | MILLFFLIPTFVTIRAEGEVEVQLKLGTIRGKSESYLDKTVHSFLGVPFAEKPLGENRFRPPRVKKSWNETLDTRKLSPACYQGRDPYEKENKPFWGAEMWNANTPISEDCLYMNIWAPAQAFNLT<br>VMVWLFGGGFWYGSPSLALYDGRALAIQGNVVVVNVNYRVGAFGFLYYDHEDVPGNMGLLDQQLALYWIKDHIFNLGGDPSRIVLFGESAGASSIVAHMIAPSSRHLFTNGILQSGALDNRWAM<br>DSPSHALSKSIDLASRLNCTIDGQSGLPFTMDQTVSCMRDLPAQKLVDELWNFNVNFLEFPFVIVSRDRNFFKHKDGFKALREGDFRDDVNIMIGINHDEGNFWNIYNLAGYFDKEVQPELNRTA<br>FHDCIETAFKVQPTLVRNAAKFVNQMVGDYFFTCDSLWMADMLSEKRNGKSKSGKVFIYHFAQTTSANPWPEWTGVMHGYEIEFVFGAPLANPSSYKKEKVDREQKFSKKVMHAQKRVECDL<br>WRHAKDIEYAAYDKSLVVSSTSLNSIISSFILLSVILF |
| ***Ppa*-UAR-22** | MNSPLLSLLFFFSLHLTSGQELSRTVSTSWGVVVGELVSPGTNDLPPVTQYLGIPYGSAPVGSDRFNMVKSAPKWTHSTKETKTLFASCIQTGLPELSETKAMRLMSSQRFDFVHRILPKLKVQS<br>EDCLYMNIFVPERLQSMSKLSVLVIVHGDEYGWGSGGPFNGTTLASYGQIIVVTINYRLGPFGFLGRCDSSSCRGNEGMSDIVSALTMLNSIVIDFGGDHTSVTLLGWGSGASLVSLLMASPITNP<br>KERLFKRAILLDGTALSPWAMSEDPQGAFLDLAAKLNCVTEESKKSPSALYNGGMDQVIKCMQDHSPQNISNAAQTIHAPSFLSSFAPIIDGQFVPNKPSLLFSPQYGNLFRDIDLMVGTASFPSH<br>HMMGDEELKNGISAEKRNKMIRTLVRNLYKFHRNEIMAAILNEYTDWTNPKESSRALRDSVLSILSDSLFASPLIRTLRAHSSDETSRNSKTFFFHFGHETRSWSAEQANSGIRGSFSGDHIPYILG<br>YPLSRKDNDEKLYYGFDSDDRGLSRVMMNYVSNFVKSGDPQMPTKMSKQSQMEETFARVAWPQYNQMNREAYLEIGVNRDNPAVVKNYYRNAHVGFWSGLVPQLHAAGKEGVTVPEEHHF<br>LPDYFKTDSFYGDVRPFGGRSNIPIPPPPMPPTPPTMVKPSSPSPSSSSDQSPSSSTIASSSYSTMLSLTVAVGCGLLALNICVFFGMYRTCMNNNRGKKKLQLQYQTYSANHGQNPDVQYSIN<br>SPLALPPPPPSSNHSQSLMDDVLPPTINGIICGSLPRSNGNSHYSRGTVSSHTYTTTFPHHHHHNHEQACDKHERDRQEPLLSASQKTSIAGIRAGVSPTCPRHGRTAIANAAASRGNSLASGAT<br>LGGSTLDRDPMRSLPPSSIPIEEIQSFIAHSFAQFLHSIESINQWISPIESMSSNSVVSLVVYGVVAAFGVIINVLVLISFVRGRLASSQYSSIYILALQTLIVDVIFITIHLFYWIPKAINPDFLSVEGYDSFI<br>LHLVDSIGTYAWFHNALSHVFISSNRFVVLVFFERSFFTRRRVIIMAIVHHFLALTITLLTQFAIPCCRLSFDFSIFSYFNVPNGDLPNYSDIFFTVPVNTLSTVTSVVCYTTIFITVRRERTATKSMQISD<br>AKRKSEYSYALQFASLAGVFSFCWISFRVFPFIIPLSDPNLVWVFGLTTVFTMVNCFMNALVYLVNNKDVQRNLSSLYGYEPTLIQPTTMHTVSSVGMR |
| ***Ppa*-UAR-23** | LFQMSFLIVLFLVPSCLLAQNQRYVTVTTPQGAMQGAKLDYGTNTNDLYYGSAYSFLGVPYAQPPTGQLRFSEPVPMQPSNQLYDATYLRGRCPQAGVSPGPVSEDCLYLNIFTTQVGNTTSLN<br>AVMIFIGGEGGFTRGGANEAEIPGTVRNLASRGVVVVTVQYRLGALGFFNLANSGVPANLGMQDQVMALQWIQENIRSFGGDPNRMTLCGHADGACAVAAHALSPMSNDLFQQAILQSGSIYT<br>CYQDTPQPATTPMPTLQRDPGLAYQPLNQQQQGQQQQQQQQPYQQQQQQYVPPYTTTPTPRQQTVADEDPSLQLAMAMCNVSSDQLRAGYSNARLRTCLQSLTVDVFVQFQGYRTSKWVI<br>VRDNSFMPGSPQSLSQNARRIPMIIGTVQDEDADYVFRMIADGSARGQSEQQLFDGWFVDFAKKNKINGTDASQVKNIIENNYGITQQPQQNQQFYNPPQNGNNGQNQNYQIVSQSNANTYN<br>PNNGNNQMQPTNNGYQQQTYSNTQFSQSTGGNNQQTMQYLQVISKISSDANGVSQSVSQVNLVQQQGADTRLFQFTHVSELGRSNVPNTGDWKPVFRGQDQYFLFMSQTVWTTGQPTAA<br>DMRVANEMGQKWTDFAKTGQVQNWQSTPPGQYNYCNLNAQPTMQQNYAPTARAVFNDQVYPIVQQATNTAPFIPSPVPSPPLAANQNANVMVHNSNGSSSWSVSFSIPASSLLPFR |
| ***Ppa*-UAR-24** | MNQLKEPNRTSLSYHWLVPSIDPSDSSLSFPPFSCLNGETEMIVSIVLSSVIFSLGASQLPEPTRPIDIWNPFGTTTTPRPPGGGIGENEVIIRLNIGDIVGKKVKLSDLPWTPDEDPLEQIPPFRDRL<br>EPNPLPKNNSVTVFTFLGVPYAEPPTGERRFKPPQQLINFPGTNPYLAFRWGAACPQDVENEPKFTFRELYPFDVSEDCLYLNIFTPDVSKSSGRIYPVIVFFHGGNFQTGSANEWPGHALSSRG<br>VVVVTVNYRLGAFGFMSLGDKTGNFGLQDQRQALIWVRDYISSFGGDPQAVTIVGHDAGGVSVGLHMLSPRSKNLFRSASSMSGAEVSYHSLIGKPALAYNNTIKLGRYLGCTQTLPQNVWDCI<br>LTRSTDDIIRATTSIPIEYNRYLFMPTIDGENVITNPLWALNNIPTGGALYPSPVPYLTGMNAQDGTEVILEDRYLAEYNQFNNVDEEYMKSFALEYSFRHNYTMNKEAIAEAIISRYTFWPDRANEW<br>MKREKFIELTTDCYYTAPIALSAHLHSAAGSRTFMYVNNYNFSRDNDALRFIPSWMAVCRECDLYLMFGYAFLPVDLRPQILKNATFTDTDRNASQLFSTLFRRFAYHQNPNLLYDGSWAVHEPRG |

| | |
|---|---|
| | HWYMNFNYSKQDEMTAPGKIERDYRYEDVAFWNEYIPQLVNYMTTTFPPEDVEDRRVNVTLKWIIGILVLVVIALIVIAGATGYKVCEGSLYEEAEMHKLVNVKSDAPSQSSMYSPRGLPKRGRIPPSSNLSSL |
| ***Ppa*-UAR-25** | MVDKWAPGRIAAKRFGDSCNSFTSSHVPGTTYSEDCLFINIITPKIGKANLPVFFYIHGGFYEAGSSTYFGYKKLAEKVASEGIVVVTINYRLGPFGFLSLGDSSAPGNLGLWDQTLALRFARELLPSFGGDINRITVGGHSAGSASASALQFSPHSNTLFAQSILLSGSSLAEFAQSEMVVEESKELLREFGCPVTSPQLGLDCLRQRTPEEIIEAVEMIGTSRRHPNVVKYNPRIDGDFFPLPITELSKTAPKKRVLAGATDQESALFVMNEQMAWIAAIALGKDEQESFSREDLLDFIERVVVTNEEHGSSARAFRQLLVDFYAGTDDGQDSMVYLQRYSDLNSDLQFLIPLYQEVRLNLRNNWPTFLYVLQHNTRSFPREGFAVQGMYHGDELEHFTDSYVLHPLKPSEQEDHTFGKHFVASMANFIKNGDPSSPSFSWSPISKARPFQHASLAVDMIFKTNAYRSDALRLWLDTIPQSVSGDLLRKSRLPGTEGLIVHTEL |
| ***Ppa*-UAR-26** | FPAMRLLFLTLVLPSLALCSTVDTPFGEIEGFEHITEQGIPTHVFLGVRYGQAPDVHGRFKKPEMVEKWAPFTHSAKHFGAACIPFTETFLIPDENYSEDCLFMNIITPTLAGKDLPVFVFIHGGGYEIGASNHYGYKRLAERVAAEGIVVVTINYRLGPFGFFSLGDSISPGNMGLWDQTLALRFLHEVLPSFGGDSSRITVGGHSAGSCSASALQFSPHSNTLFSQAILMSGSTLSEFALSQSVVEESKKLAEELNCLDKSTTKTLDCLQERSAHEIIQAVDKIGTTRRHTNILKFHPRFDGEFFPHAIERLAKESPRKRTMSGATDQESAIFVLFEELSFLTGLALSKPNWATYSRENLIDFIETVVVTDKEHHDSASAFRRHLIDFYVEKDEKKGDAQFYLQRFSDLASDLQFLIPLYHEIQIKLANNWPSFLYILQHSAKDSPRDELPVTGAFHGDEMAHFTDSNGFYPVKDGDDEFESFGVNFATALVNFIKNGDPSSPAFSWPPVTKERPFQHASITSKMTLKKEAYRSDAARLWLESVPQSVSGELLRKTRLPGAEALIVHTEL |
| ***Ppa*-UAR-27** | MRIPFILLISFHCAHSSIRHVHPIELTIQSGSIRGEYLRQKGNDYAIFKGIPYAAPPVGSLRFQMPEPPARWRGVMNATQYSALCVQKREGLPSHPERSQAHFSEDCLYLNVFAPTQFTNDTYPVVVFIHGGKFQYGGASDLSQQAILDNFVSRKIVFVSMNYRLGPLGFVSTGDSSLPGNIGLWDMLLALKWTKTNAQVFGGDPDNILVMGNGAGAAAASILALSPKAEGLFHRVLLMSGSALSPGAVRETAVNATWNLDTKLHCRSFNSSQLIDCFRKKIKDEILNFDRPDPDEYEEFVPIVDGPGGIIPEPPERTLKRTKLKHALMIGTTKDESSLYIYEKNINVSAIVTDADAEKYTLRLTNTTEKISNKQLVHQACIHEYINTKVDPAYSETILQHSVLKMFSHFWYDAPSSKLAEYYLNRSQPVYLYSFDHISENFYDIDRAFHGVDAMHLFNIEPKFLHKRKDMNWKLDERVTEIFSELISNFARAGSPTPMNSGYAFNWTSVEINELNYLSITDSPQMEVGFRWQGHVFWNRYIAQLEDVDVGNLHKVAALEKSLGDFQLATWMLLFSTLFFFAILVGLACYCTRKESEDDDL |
| ***Ppa*-UAR-28** | MLSLLALTIFCPFLIAGLSFDDAAGTVSLKISQGELLGKRINSYDNRTGYGFLGIPFAAPPVGPLRFAKPVAAAAWDGVRNVTEKGVACPAVAELTYKRPEGGPMGEDCLHSYVYASKNCLLKGGCPIMWVLHGGKYNFESPVVFVDDVIVHNFASDGQDVVVIIPAYRLENFGFLNIAPGLNTSAPPNTAVWDLLLALQWVHREAFHFGGDAKKITVFGHSAGSQFADVISTSPHFNGLFRGMSMMSGADTYYEGGIQANSLASWVTAKLLGCANDATNWKDIVVVEGVIECMRSKTWEEIIAANKQLNSYAEDYHGPHADGPGGILPLPPSTLSRTRLPVPIMIGTTSAEFHDTKYALTADGTAGDLNKVAELCEGVGYGTGYIHPDVMTKRCLDYYMQGKNVMSLQQDTMFFAPTFLTARDMAKGAVKNKVYLYSFTYTGVGAAFKKYVKLDPEDHPSHSEDYVYILGMHRGNFSAKDYEIEKIYSGMVLNFVKTGYPNPGKGQPAWKPFTKLGRDYYQIDFDDNLRMPGPKKGYQDGAIKLWVDDAEKYGGGINPPQDYPAHADRFNPMDMVTVMEAHATNINVAHDKTLETELSNYDDWTTYINNRRFKHLQGLKNFRSMAVEQLDLTQVAAPENNGISYEPERAIFRMPVEILIICISAVLIFLLLRVCLHFVGVCQERKGYEEF |
| ***Ppa*-UAR-29** | MAVVHGGKFNFESPAVFPEDVIVDNPDILAGLKWIQREIHHFGGDKSRVTLFGHSSGASLVDTLSLSPRSIGLFSQLVIASAASASEINKNVNIAASWAIVKATGCAPSHYDYHSGSMEKTELALRCLRSIPYNKIIDAQRSLYNSTEDFYGPSIDGDILPSSNWNLIPIRPLYPTLIGTVNAESRIGRYMLTPDAKSVNSTLIDEYCRHVGYGIWFERREEVYKACFDKYSKGSPSAFDNRWGLDMEDRPAHSEEYIYILGLHRANFSAKDYEIQEIFSKVFADFVNKGDPTPEGRKWSLYEENERNYFEIDFPTNLISSASFPGPSTKFMSRANDFWIDVIEKK |
| ***Ppa*-UAR-30** | MGTDIAMRSLILVSLCFLLARAWEEISTDVEVDTPRGRLKGRHVNFGSNTTDFYYGEADIFLGIPYVLPPQRFKRPVANCQYTNDGSTLTVQQYGPSCYQAGKGCPDENSIPMSEDCLTLNVFTPDVTSNYKYPVMVYIHGGGLMYGCAAEYPYDGAVVNMVNRGVVVVTIQYRLSTLGFFTTYTEDFPSNLGMLDQVEALRWVKDQISHFGGDPFRITIFGQSAGGASVSAHTFSPLSQDLFQQAIMESGVALTTFEGSLGYDNLSRKRAQEICKVSQADFDNGKWDKMSDCLYKSDPKDLVKLDTVNIIGWKISAGDDFMPDIPNHLAPFRNNIPVLIGSMHDEWAYYDMQLMNLGIAQYDNYSREMFEFEYDVLGGFLAERKVDMLKILEDVYAGVEAPDYDHNLWMRAMSGAFTSAAFTSFIQRDFMDYLANGNKRMWLYEMTYPKAIGRIYDLPNFPEAVFHTAEVSYLWKRKKQFNEAVSTGYINQDDYDLSEWFGATWTNFAKYGQPYLNDEWPNIPTTGPETYMEITGPKPVININNGYKTRDNIIWNQVLVALHGDFFPAQFSTPRFSKEDFDKIKANASLQDGVCSMNDPTQQPPTMKSTISSMMPTTEFISSTLSYFSTTTTSSSVSTDVEVDTPRGRLKGRHVNFGSNTSDFYYGEADIFLGIPYVLPPQRFKRSVANCQYTNDGSTLTVQQYGPACYQAGNGCPDENAPMSEDCLTLNVFTPDVNSNYKYPVMLYIHGGGLMFGCAGEYPYDGAIANLVNRGVVVVTFQYRLSTLGFFTTYTEDFPSNLGMLDQVEALRWVKDQISHFGGDPFRITLFGQSAGAASVSAHTYSPLSQDLFQQAITESCVALATFEGSLGYDNLSRKRAQELCHVSQVDFDTGKWDKMSDCLYKMDPKDLVKLDAINVIGWKISSGDDFMPDIPNHLAPFRNNIPILMGSMHDEWSYYDMTMMSLGIAQYDNYTREMFEFEFNILGSFLGDRKKDMLKILEDVYAGVEAPDDDHNLWMRAMSGVAFTSAAFTSFMQRDFQDYLANGNKRMWLYLMTYPKAIGRLYDLPGFPDAVFHNAELAYLWKMKKQFNEAVSTGYVNNDDYDLSEWFGATWTNFAKYGRPCLNDDWPNIPTSGPEIYMDITGPKPVIRNNGFMSTDNIIWNQVLVALHGDFFPAQFSTPRFSKEDFDKIKASASLQDGVCAMNDPTQQLPITTMKSTITTMLTTTEFISSTLSYFSTTTTSSSLLFGWNSIIFVLLVLIR |
| ***Ppa*-UAR-31** | MSLRRRRGQPRCPQGDDARGFGYQTSEDCLYLNVFTPNISDGRKRAVMFYVPGGAYQLGGADIYDYKGAVRNLVSRDVVVVVTQYRLSTIGFFTTFSPEFPANRAIFDVLMALRWTQNEIANFGGDPDRVTLFGHSAGAVITDALSYSPLAKGLFQQVLIQSGPIIDTFKCDIRRSGSNSSIPTGYGRNATADEDRVVKLCNITDMRVPVRNATLRRLQHCFNRLTGEAMVAVDNGASLYGVTLDGVIFPLGPTANLARLAPNYAIMQGDTPDEYAFSIDEVAAGDISTVNETTLDYYLKKNVPYLSKDQLATYKRAVLSFYDPNRTLAVDDHLGWTKMVSRVISDHNFFGPARKELQWYKTNGNSRNVFLYNYDYFNSFCRNNFEVEGWMPSIHCSELCFLWFYPYEWIKAQEEGKVRPEDLRVADNLGVAWTNFAKYGNPGWSPMRDRFEYVTISDTVSDVQLDWGEAANKLYNEILPSLLKMDLPPFHIDPAVQNQIERQAPSILKTWYNATCPTWKPLYDGELTVYPLKSQETMNAKYGRYRITPRQASGNNLGFLKCFIITLAASLQM |
| ***Ppa*-UAR-32** | MLLLLFLFFLIQSLAAQDASPELTVTAQTSAGPVRGFQVDYGSDKSRLYYGSASVFLGIPFAKAPLGELRFKLPQALCRFEGDVGSTEYKPRCPQSHDYHGFDYITSEDCLYLNVITPNITDGRKRAVMFYIPGGVYVLGGADIYHYKGAVRNLASRDVVVVVIQYRISTIGFFTTYTPEFPANRGIFDVLLALRWTNNEIANFGGDPNRITIFGHSAGAHIADALATPDFFDNDTTPEVEHAEKLCNLTLSSSSMHDSIPLLQECFSDIPGEEMIKADNAGKSYSMSLDGVIFPIAPIAELARIRPNYVVMLGDTTDEFVFSMNEILDGNISTVGESMMEKYLKEKFGGQSKDQFALIKKAVLSFYTPNGAPNKDDHLAWTKLISRAITDKDYYAPARKELQWFKNNNNSNIVFLYDFDYFNSFCANSYPIGDWRPSYHCSELCFLWFYSYEWERAETEGRIKQSDLEVANNLGQAWTNFAKYGNPGWSPMRDQFDFVTIADTVSDTQADWGGPANRLYNEIIPAMLKTDLNRFLVDPRMKNQIEKHAPSIMNSWLAATCPTWKPLTTTTLTVFPLMTEEATVPRGHRNHTSQPDSSSYSCMFCSVVVLLATLYLSL |

| | |
|---|---|
| ***Ppa*-UAR-33** | MLLLLLLLQLLQLSHQQQPSSFTTRVTTSRGVVRGFRVDYGSNRSQLYYGTADVFLGIPFAQPPVGEARFKLPRPICRYTSEVGELSYKPRCLQQSPPSPQPVRTDEDCLYLNVFTPETTVARRY<br>PVMVWVHGGAFVSGAAEQFHYKGAIRNLVSNGVVVVTIQYRLGMLGFFTTYTPEFPANRGLFDQLLALRWVKEEIGAFGGDPERITAFGESAGAVSISGLSLSPLARGLFNRMAITSGSVYLSFQ<br>APNDVRGSIERDRAAQLCNITGDPMVHPALRVQLSNCLQSMPASGLISKESPPLLASSVKPTNIGWSPSRDGVMFPSDPEVLACSRPPMDALIGDMKEEYAFFMSEYMNGNISYIGATSLREALL<br>NRGKGYLTQEQLSLISDRMIRSYQTQKLAANDNLGWFKVIVDAMTGEMISSIMKRDVEWLKSSRSRVFLFTFDHPSTVCNIMPSRLQGFNGNLGRISASDLAVANNAGRVWTNFAKFGNAGWTE<br>SGSAYSFMKFDDKLNTMGNDWRGTADRLYNKEFPRLIGDYPPTKLSPIVEEMLKGQGAKVLDQWNRNLAMCHFPTRGKNILDLVLCDSPIIRSVKPDIPLSDHVGITVTVSFETVVKLPPTAPTRN<br>FSLANWDIINAQIACHDWTRAISNKTATEAYCYFSNFLNSLLDDFVPLKPSKSSSGFPKFLSILHDRLQRLHSVAPNSDSTHSLRARFNKALKTFEIMRENNAINSGNANLFFQYCKTRFKPSTSSLP<br>GIVDCSGAVLLTDQDKATAFSKFFSKVQTPPMISPLPRSLPSNSSFDIPYISIEQILSAISQLVPKVNGSPDRIPNLVYTKSYKIYIRPLLESYTVIWNPTAIGLVNKLESVQLRDTGILTCIKRHLPCPGYL<br>TVIIYDEPLHLSSLSPKLNCKHGLPQNPIPKSSPEPEYHLLPPYRP |
| ***Ppa*-UAR-34** | MRLALALLVLLCLTEAQQRQFTTRVQTSRGTVRGFRVDYGSDRNQLFYGAADIFMGIPYAQPPVGPLRFQLPRPICQYQGEVGELDFKPKCAQAESPFSPGGTSAEDCLYLNVFTPSTAVPRAYP<br>VMFWIHGGAFEIGGANEYDYQGAVRNLVSNGVVVVTVQYRLGPFGFFTTYTEEFPPNRGLYDQIMALRWVQQEIAAFGGNPDMVTMFGESAGGVSVSALSLSPLARGLFHRIIPQSGSIQVLFQ<br>SPNDVRGAIERDRAARWCNVTADPITNRDARRTLQTCLANITVARLVDAPPPPPGTGVFAISINWNPVRDGAMFPEDPEVLAMSRPRYDALLLDTRDEHALFSAPFFSGNVSAQGPGIIRDTFLRR<br>GYGYLTDEQLDRVVSLIVQTYQTRQLAANDNLGWYRLAVEAFSGEQFQRQMRQDANWLKASGSRTYLTTFTYEPRICGPISLTTGYNPIPHFAEICYLWFTSANWQSAANSSTIKPADLAVANNM<br>GLAWTNFAKTGQTGWPESGPNFSFVNFADTLVVGNDWHGKEDRLFIKTLPSLIGNYPPTKVSPIIDSVLKSNGAKILAAWQQSLSSCRR |
| ***Ppa*-UAR-35** | MRLALALLVLLCLTEAQQRQFTTRVQTSRGTVRGFRVDYGSDRNQLFYGAADIFMGIPYAQPPVGPLRFQLPRPICQYQGEVGELDFKPKCAQAESPFSPGGTSAEDCLYLNVFTPS |
| ***Ppa*-UAR-36** | MPRRSLLIIFLSFFIILIACISVVVPIVIVNATKGEDNIPDEFTNRVKTTRGTVRGFHVDYGSDRNQLFFGAAEIFLGLPFAQPPVDELRFQVSLCILSYVKYVIYLPRPVCKYYGEVGELEYKMSCAQP<br>PQSPTNPKPSSEDCLYLNVFSPSTATPGKYPVMIWIHGGSFLFSGADEFHYKGAVRNFVSNGVVVVTIQYRLGPFGFFTTFTPEFPPNRGIYDQIMALRWVKEEISAFGGDPDRITLFGESAGAMS<br>VSALSLTPFAQGLFHRIILESGAANAVFRQPNDVRGTMERQRAAQLCDVIADPITNRAARQTLQDCMANKTTFELTKLDITWFMPGTLSWAIVRDGAMFPETPEVLARSRQPFDALLMDMRDEQA<br>LFVEEYYTGSVNGDGPNTIREWLMNRGYQYLTDAS |
| ***Ppa*-UAR-37** | MTPAKVMIILTSIAAAFFLMCVIITTISTAQEKSRAETVKFTKRVYSKRGIVKGFHVDYGSNRNELFFGSADIFLGIPFAQPPVGALRFKLPQPVHKYTREEGELVYKPKCAQANDLGQVTNGSSEDC<br>LYLNVFTPNVTGKYPVIVSIHGGSFVARGADDFHYKGAVRNFVSNGVVVVTIQYRLGPFGFFTTFTPDFPANLGLFDQILALKWVKQEILAFGGDPNQITLYGQSAGAISASGLSLSLLTRGLFNRLI<br>MNSGSVLVSFFVPNDPRGSIQQLRAAQWCNVTEDLMTSPTAPATLTSCLANLSTEEILKYDAALSLPRALRWAPVRDGAIFPDDPEVLALSRPAYNALLMDMPVEQAMFDPVYKSGNVSGFGPN<br>SLKKALLNRHYGYLSDAQLDRMVDILIANYKTSELDDNDHLAWFKQVMHIISAERYTNHGRLEAQWLKAGGSRTYLSIFTYEKRICGMHSAIPGYDPTTHYTDVCYIWFTSAEWEKAASDGKVTA<br>RDRAVADNFARVYSQFVKNGTTGWSESGNDYQWMHFNDSLDQMGKNWRAHDNYVFTEALPIVIGANPPIKLAEGIVSALKLNGELVLNSWEQLLNSRPP |
| ***Ppa*-UAR-38** | MSPIKCMVILTSIALTLKFTKRVYTSRGIVRGFHVDYGANRKELFHGCAVIFLGIPFAQPPVGALRFKLPMAVDKYSDEEGQLNHNEILRSRCAQANETGKPGQPSSEDCLYLNVFTPNVTGRYPV<br>MVWIHGGAHFKGGANDFHYKKFRVQWRGFFTTFTPEFPANLGLFDQILALRWVNEEISAFGGDPDRITLFGQSAGAMSISGLSLSPLTKGLFNRMILDSGSVLVSFFVPNDKRGSIQRIRAAQWC<br>KIKDDPMSSETALAELRKCLSNLSTEEIVKDDVTLSHPGSLRWAPVRDGAVFPDDPEVLARSRPAYDALLLDMPVEQAMFAPAYKSGNVSTFGIRSLKKSLLNRHYGYLSDVQLKRMLDILMENYK<br>TKEIDKNDHLEWFKLNMDIISAERYTNYGRLEAEWLKSQGSRVFLATFTYGERKQHTPMTSAICGSLLQNGKRHGWKGKLLPEIWQWPTILVETSGWPESGSDYKFMNFDDSLDQMSNNWRA<br>MDHYVFTEAMPPVIGKNPSINLTQDVESLLKYYVYSTPGTNQSILALDNK |
| ***Ppa*-UAR-39** | MAEDCMYLNVITPNVTGKYPVMVYIHGGTFTTGGADIYHWKGAIRNLVSHGVVVVTFQYRLGMIGFFTTFTERFPPNRGMYDQILALRWVNEEIVNFGGDSSK |
| ***Ppa*-UAR-40** | ETKNAIMFHSLLYALFLHLVTSESITVETSRGTVQGFDQDFGSDTTQRFYGYGQMFLGVPFAKPPLGELRFKLPEDICKYTDSGDVHNATFYRPRCWQNEPGHELSEDCLCLNVMTPNITGNYPV<br>MVYIPGGTFTTGGAEKYHWKGAIRNLLSRGVVVVTIQYRVGVIGFFTTYTDNFPPNRGIYDQILALRWVNEEIASFGGDASRITIFGQSAGASSASVLSLSPLARGLFHQIIQESGSVEQELETITDA<br>RGSIHQDRAQKICQINSTDWGSAAKDEELLNCLVQASPQDLVAYDKTDNLNWNVEIDGAVLPDYPEILAKSRPTYPALIGDMLEEFAPFIPGVTDGNLTNITSASNPFFVQLAWPNYDDATAKNISDI<br>MIAGFCDGAIPPDSDHLAWTRLITDMFTGLQFSSTMIRDVQWHLNAGNKNVWTFTFAYSSPLSTSIIFDDWRPVVHSSELPFLWFYPSTWETANATDADFAVADYMGQLWTDFAKNGELSLPRVG<br>EDMNYVEIGEILKTKSDWRSTANTVYNEELPSYLGEFPPLVMSDESWVQLKDLGKKVLAKWNPNTCPSTTSTPTSESTTTTKESASHHSIFVIVLCFLCLN |
| ***Ppa*-UAR-41** | LQMIPILLCVVFLSISCSSTRIIVNTSRGSVQGFDHDFGNDTTQLFYGYGQVFLGIPFSQPPVGERKFSLPEPLCQYNERGEVHDATYYRPHCTQTLLTPEPANEMSDDCLYLNVITPNISGSYPVM<br>VWIHGGTFTTGGAEIYHWKGAIRNLVSRGIVLVKIQYRLGLAGFFTTFSERFPPNRGIYDQILALRWVNEEIAQFGGDPSRITIYGQSAGAKSVSDLSLSPLSRGLFHQLIQSSGVDLMEVIDDPRGS<br>IHQRRAEQICGINSTDWGSEEKDQALLDCFLRATPEELVAYDGKDRFNWNVAIDGAFLSDEPENLAKSRPQYPALIGDMLEDYAVFIKGVSFGLLSNISSLTALDILREKWPNYDEDSLKILTDSLIDG<br>HSNGVRPAKDDHMGWARLLSEIFTGLFFDNYAIRDARWHRNTGNDNVWLFTYTHRSLLPFQMEIDGWIPISHSSELPYVWFYPQNWETFNASKADFTVADYLGETWANFVKHGELSLPKAKES<br>LNVVEFGDSITTKSSWRSVSNKIRIAEQLPKYKGIFTPLKTSEHNWRRLRDLGEKVLSKWNSMKCLKSETKSLRTTTRGAPSTSRMALLMLLLVQCLQVIRNMH |
| ***Ppa*-UAR-42** | MIRPSLCVVFLPLCICEFFTVETSRGSVQGFDHDFGSDKSQRFYGYGQMFLGIPYATAPLREKRFTLPEDICHYNDRGEVHNATYYRPRCWQTYDYLQPANEMDEDCLYLNVLTPNVTGKYPVM<br>VYIHGGTFTTGGADIYHWKGAVRNLVSRGVVVVTIQYRLGMIGFFTTYTENFPPNRGLYDQIVALRWVNEEIANFGGDPSRITIFGQSAGAASVSDLSLSPLSRGLFHQLIQTSGSSIQEIETIESPG<br>GSIHQDRAKQICDVDSSDWGSPDKDQALRNCLVKATPQQLISYDFTDGNSWNAAIDGAFLPDYPENLGKSRPKYPVLIGDMLEEIALSIPDDISNVTEQTAFDKMKMMYSFYDEETINNITETMIAG<br>YSNGSVPADNDHMGWTQLMTQMFTGLTFDSNMIRDVQWHRNAGNEAVWLFTLSHRSLLPLNKQIDDWIPVGHCSDLPYLWFYPDIWETYNASNADFAVTDYMGGLWTDFAKNGELPQPRAG<br>KVMNYIDIGDLPSLKSNWRSVSNTVYNEELPAYLGEFPPLKISDDSWNKLKELGEKIVTKWNSMKCPDISTTTRLQTTTSKGAFQGTLAAAVIIILLKSCANVGM |
| ***Ppa*-UAR-43** | MLILLLLPLATASSITVYTSRGSVQGFDHNFGSDKSKPFYGYGQVFLGIPYAKAPLGERRFTLPEDICQYNDKDEVHDASYYRPRCYQVQDDLQPADNMSEDCLYLNVVTPNVTGSYPVMVYIHG<br>GGLQTGGADVYHWKGTIRNLVSRGVVVVTIQYRVGLIGFFTTYTEKFPPNRGFYDQILSLQWVQDEIKHFGGDANKVTIFGQSAGGSSVSDLSLSPLARGLFHRLIQTSGTALLEVETPEDPRGSI<br>HKERARQICNIDDSNWSTAEKDDAIMKCLLAATPEQLIEYDGSTSKGWNPTLDGAFWPDYPPSLANSRPKYPLLMTDMLDEYAYFLPGLHSNDITGIGPGTSMKLFSDQWPNYDQNTVQRLNDL |

| | |
|---|---|
| | FISSYDNGKVPADNDHLGWTKLTSDIWTGFFFNAFMVRDVEWHKANDNNDIWLFTFTHANKLGIPIDVEGWIPVGHCSELPYLWFYPDVWDNASVTLTDDDFTIADHMGRIYTDFAKNGELPYDRAGANRNYLEIDKTLTKKTNWREKENDVFNHQMLQILGEYPPLTISQKSWDMLNDLGKKVLKTWNSMECSYPNLTTTTKSGQFISIVSAVIASVIVIFLL |
| ***Ppa*-UAR-44** | MNRSLPCVLIVPLVCLCDLITVDTSRGPVQGFDHDFGNDRSQRFFGYGQMFLGIPYAKAPLGERRFTLPEDIRRYNDSGELHNATYYRPRCWQPQGTIQKAGQMDEDCLYLNVMTPNITGKYPVMVYIHGGKFTSGGADVYHWKGAVRNLVSRGVVVVTIQFRVGLIGFFTTFTERFPPNRGFYDKMLALRWVNEEIANFGGDANRVTIFGQSSGASSVFDLSLSPLARGLFHQVIQSSGPTLMQIGTIENPLGSIHQDRAMQSDSKNWNPVIDGAVLPDYPADLSKYRPRYPTMIVDMLEEGAYLIPGVNDGDVCNVTRETAFELMRSLFPYYHEVTMTNMTEMMFEGFAKGHFPEEEDRLGWAQLLSAMFSGLSFNSMMVRDALWHRNAENEDVWLFTLAHRSLLPFNKQVEGWIPVDHCYDLPYLWFYPSVWETHNATDVDIAVADNMGAIYTDFAKNGKLSFPRAGKSMNYVEIREFLTEKSNWRSDANKVFNEEFPAYLGEFPKLKMSDESCTLGTIEVEFDEPDISHEIDEIYSTNSSKYVENGVTYLPEDIGRYNDRGEIHNATYYRPRCWQARVDFQKGYNFGEDCLYLNVMTPSVTGSYPVMFYVHGGTFTTGGADVYQWKILALRWVNEEIANFGGDPNRITISGQSSGATAVSHLSLSPLTRGLFHRVIQTSGSALTQIETMENPMGSTHQARAMQICQIDSSDWGTPAKDQALYDCFIKATPQEIIAFDLDNHLWNPVIDGAFLPDYPDILAKSRPPYPTMIVDMIEEAAYLLPGERIGDASTVSRQTAFDLMRDFFPYYDQDTVTAMTEVMYDGYAKGNIPADEDHLGWAQLESGMFTGLVFNHKMLRDAQWHLKAGNEDLWFFTLAFRTIDHCYELPYIWFYPSIWETYNASAADFVFADKMGEIYTEFVKHGKFPFPRAGKSMNYVEIREDYLLKTDWHPEANKVFNEIFPSFLGEFPKLKMSNEVDWARAMDTGKKVLSKWNSMSELKWKTKAMIISAHKAPSTARLALLMLLLAQSLKVIRTTRYSLFCDVF |
| ***Ppa*-UAR-45** | AHFPCCFRSMLSLLVFLLCSFNMNADPITVQTSRGAVLGFDQDLGNDKSQTFYGYGQMFLGIPYMKPPLGERRFTLPEDICQYNDNGEVHNATYYRPRCWQIRDGLQPADNMDEDCLYLNVYSPNVTGNYPVMFYIPGGGFTTGGGDVYHWKGAVRNLVSRGVVVVTINYRVGVIGFFTTFTEAFPPNRGMYDMMMALKWVKEEIKNFGGNTSRITIFGQSAGASAVSHLSFSPMARGLFTQTIQTSGTALLEITSPEPAKGDINKDRASELCNITVDAWGSTDYDQPLMDCLLAATPQELIAFDVSPGPWAPALDGSFLPDYPENLAKIRPHLPAIAIDMMEEAAPPSPYNEIVLPFTGPKTIPNMFELLWHDKDPEAVANLTDYAINAFSKGNPPADKDHMGWLKLVTDVATGMFFDTLFLRDVKWQTQYGNDNVWLFTLAHRSNLPFYIQLEDWIAVAHCADLPYLWFYPDIWETYNATESDFATADQFGVIWTDFAKNGKLSFDRAGTSRNYVEIDEELTMKNNWRETTDDVFNRKSIELLGDWPVLTISDDAWTMLNALGEKIKAKWAADTCSAAPVDVQTTTESDAQTTPDNLPTTSSSVPTTSTTPETTTSLCSRSLSLLSFVLTYIALSW |
| ***Ppa*-UAR-46** | MLLVLVLLLSSAISNAKLITVKTSRGSVQGFDHDFGDDKTQTFYGYGQLFLGIPYAKAPLGKKRFTLTEDICQYSDTGEVHNATYYRPRCWQFRDSLQPADNMDEDCLYLNVYSPDVTGSYPVMFYIPGGSFTTGGGDVYDWRMMALRWVNEEISHFGGDTTRITIFGQSAGASAVSHLGMSPMARGLFHQQIQNSGSIMLEILTPEPERGSIDKERAKQICNMTDTSNWQAPESDAILMDCLVNASPQEIIKYDMTTGKYWAPSIDGAILPDYPENLAKIRPHYPLIAIDMMEEAASTPPGSTDRLPMTGPNTALLMWEGFWYFRDHKAVNELNEYAINVFSNGTSPDKDDHLGWDKLDVDMNTGMLFNAMFLRDIKWQTQYGNEKVWLFTLAHRSELPFPIAIEDWIPVRHCADLPYIWFYADIWSTYNASAGDFATADHFGRIWTDFAKNGKIDYPTAGVERNYIEIDEELTMKSNWRETTDDVFNRKGVELFGEAPNLAISPDGWAMLNELGPKIKAKWEEAKAAGCPAPEVPTTTATSTTAKAVSTTTKTTKMTKMTPTSTTSEATTTTKVAITIKFFELVALICIILLKIARNSLIILFVVHNLQPNSGQLARFHEEGRAEGHAYLSGGKEVHAKLPPIDIAVNGVFGNYVFFKTREQTFKATFVNGKIKVFALRQMLQTIHHGTMCSNVFPGAESVYRLGEDLKDGLLVDVPKEKLSGLHLKDIHRKRLVYLSQVKDSIRPTICDLSENAILIEVVFCEVCLSTRSLCAASIFTNLFSQHQIYANLQIASDPVKTRIYLLEMHHLFCHLFTLDTNRRMCLRTLKLRPFCDIVGFYNGILTVGSYDNGAFEKKACYNGKEISPTETSDDERKELMQKLKNKEELNEELQRTKLTKFGEEVSRGKGALVEAESTDIRSEENASNADVRIASLMAEMDSMRQRLDNLQHDLPAAAAADPEPAADASGLTGPNTIRMRECTIFYHDYFNSPHRLFVKWNGKEIKADMPQDEFEVMGAVGDAVYIRTQRKDETIHCSILCSRKYYGNKYIYRVSEDFRHGIIFDVLDKKLTLLGMHRKKLIYVSNQKYCPTGPRLSQLTENAIVIEPPVEHCRGSFYASDSSRIIYFTTFTRKKFAGHLFALDTEEKTFLPNLSFADGTLSLLGVCNGVITATSFICVSEFNLATARLPRGYIQQSISQTGNQLLKVPDGRKEVHQTNGELLEQISQLRLQLHRLTFENEQLSEALKRSEENVFNAEQRIAELKSEMLEQSCEDPMHRSNSFSDLAMAQVGSTIKLVDKGCWEHYVQYQQLDDGTLVYLEMSTPQKFYAKFNGKRIDAVLPEGKFKMIDAHSNSVYLMKTKELLTQPFVKHYADGFLDYSNQIFEAVFTPPSALAISLLKNRRQPYEIFGTSNSVMCTTMRNCKQYVHKIWQVAESDDILNNVKETFQLMSIHRKQLYFFNMQEFLRRESVNKLRENCYAINILAKSHWISIHTRDTTSFYYLTDGMCLYTLNTETKKFLPPLQFSRHEYDAIELIGIFNGLLTLRGTHNGLNYLLTAQLPTRYYDVMDAKDVASQQDEESVNKKMDVMRSIISELQEKNAKLSAELERLSIVNKEMEISTADVFEENKELKNQLDLIRNMRSAEGTSRDFCGDVTLSVIMTMPGSPSLPGVASSSK |
| ***Ppa*-UAR-47** | QMGRQRVISSFILLFSLALLCSADPTVKTEYGEVVGFEFEEADVFLGVPYAAPPLGELRFTNPIRPTPWSEPREAKTFADACVPHSREAVTWTASEDCLYLNVIAPRRDSSSSAELAPVLFYIHGGGFEIGNAKIYGYEDFARSYAAQGVIVVSIQYRIGVLGFFTLTASDKMSGNYGLFDQVAALQFVHRNIERFGGDPKRITVFGISAGGSSASMLTLSPLSRHLIAGSIEVSGTAHAGWAIDNRVETHSEDLVDAVGCWGKRSVADIEACMRKVSVNDLYSGVEYIFEAAFSFNMLKFAPRIDGIFAPQRYEDLAQESPKIPVLTGINALESAFFILMHKSPTIHRSTIFKGEMPVFDADKFDMKTRILLNEFLDPSETDEAVNDVMTFYLGENYLEKYSKPTKEQIPWMLRQYTTFWSDVYFNLPAKYRAEERKKIGADSYVYLWEHYNEDLFKDDDPVPAAVHINEMPYITGLQALGQFEWTPEELKLKERAIQMMISFVRNGKPSVDGVFWPSYTNSSNSPYVKIDSPEWKTEEGFWDKNTRFWEEVMSKYSYDFVRFKKRTVPQMQIPPPPTQKNHEEL |
| ***Ppa*-UAR-48** | MLAMRSLLVIFSTIGLAQSATFPRKTLVNGVVEGFCLIDSPVQVNAFLGVPFAEPPVRFEKPLPKNNWSGVLATKTQPNMCTQVTGAANSEDCLYLNVYAPAVAPSAALNKDSCTHGLPVFVVVHGGAFATGSAQEGAPEHIARYLAAKGIVVVAIQYRVGPLGFCTTKDSAMPGNYGMWDAKIAFEWVRDNIAAFGGNPNDVTAYGGSAGAALIDGMHLSPLTTSDDHLFHKMVLFSGAARDMWDAHTTEHCEERATAIGLSWTDSASFKSALLSANAADLAGGWQVIGEENFDYLLCGEHGISSTIGNADSKKSTHSDWAPVLDGDFFPSTPAAMRAATQPKPSIFGNYFASCFFGVDGAATQHKPSFFGISFLEGAGMSGAITIDHTTVEKIVDFMVPATIANRSLFQSYFLESYRNQGQILEPTAIDKHAMMAVNVFKHFNPAPIGGLYPFISYATHSFDQWYSLGLPGFTLNTADQIVIDIYTTALVNFAKTGNPNGSSASALPVPWVAATAQNPSLNYVIETTPSMDAQFFYGRPNLNNILNKIGGTFKPV |
| ***Ppa*-UAR-49** | MRSLLCIAATVGLALSACSTAAPQVTVANGIVEGFCVNDDVVQVNGFFGIPFAEPPVRFEANTAKKPVPKTNWTGVKATKTMPAMCTQATPGDSEDCLYLNVYAPDGAASGSLPVFVIVHGGAFAFGSANDGTPAHIAKYLAAKGLVIVMIQYRVGPLGFCTTKDSVMPGNYGMWDAKMAFEWVRDNIAAFGGNPNDVTAYGGSSGAALIDAMHLSPLATNLFHKMALFSGAATDIWDAPTKQKCEERAASLGLTWTDSASFKAAMLSAPASGLGSGWLFHGESTFDTIYADWSPVYDGDFFPTTGAAMRAATQPKPSIYGISFLEGAGTSGALNITSETVEKIVNYMVPATVGNRSLFQSYLIESYRNQALVLEPTAIDKHAAQAALGERNRAAGMDAVLRRNFELFGTNQTYYRYVFKHFNPATIGTRYPFISFASHAFDQYYSIGKPGFTFTGDDLTVVNLYTTPRPGHNPNGAGPVSSLAVPWVASTPANPSLNYVIESTPSMDPQFFFGRPHLNNVLNKIGGTFRPV |
| ***Ppa*-UAR-50** | MPLSLFSKMGNSRSAPSTPVIATRDGKIQGKRLVDEPNFKADAYLGIPFAQPPLGDLRFRKPVAPNKWADTRECFEHPKKCIQVPFELLSQNDKDWPASEDCLYLNLFCPGDYTLKGQFLIILRLLS |

| | |
|---|---|
| | VWFPWLSHDNYNKYPVMLFIHGGGYACGSVKAYGEQGICDGLVRQGVVVVTIQYRLGIFGFFSTGDEVCPGNLGLWDQIEALKWVKDNIEHFGGDKARIYFLIISLSTYDNVTILGQSAGGASVDL<br>LSLSPHTKGLIHRVIPMAGNAHAPWAHRKTTADYCREYAETKLGIKADSSSELISQLRRVSADRLATELHMETALAETFLGFCPVIDGDLLPRPVEELRKVAPVLPVMAGVTSLECGLFIPNKDLTEE<br>NIKKMAEAMLPDSAIKESTDALVKMYREVALRKQPKNELWRAMVEIAGDRMFNISTVETLKKCSQRGAATYFYVFDYYNPKCLGPFSEMAPSSDASHCSELAYVFNAGVAAPFAFCDHDKQMAD<br>MTMKAFANFAKRGNPNDEGEKCWLPFDFEHPGRHMVIDLKPRMEEEYLGGRCTRWMELQKHE |
| ***Ppa*-UAR-51** | MTASSPTITTCKGTIRGKRLIDEPDFQADAYLGVPYARPPLGELRFKKPLPSLEWAGTRECVHHPAKCVQQLSELLSQNDKDWPASEDCLYLNVFSPGDYPANNKRPVLVFIHGGGFARGSIRAY<br>GDRAICDGLVRQGLVVVMIQYRLGTCPGNFGLWDQIAALRWIQANIEQFGGDKDNVTICGQSAGGACADLLSLSPHANGCAHFLPISQFHGRPRPPLREGGLFHRTILMSGNARAPWSHRPSTA<br>ESCRQYSETVLGVKADSNDELIAQLRLLPSAVLESRLLATTARFEFHPVIDGDLLPAPIEQLIVHAKALPVMAGATTLEGGLHLPDKDLTDAHLKKAVTALLRKDDCDESIEALSAKYKEVALRKQPE<br>HAVWRAMAEVNSDLVFNLPTVQTLQERIRRGIPPYFYVFDYYNKACIGPLAQNSPCADTPHCSELAYLFNDGVVAPFAFSEDDEQVAHLVMRAFANFAKSGNPNDDGSCPWQPCDVLLPTRHM<br>VLGTQPRMQQQFKKGRCERVIGLMK |
| ***Ppa*-UAR-52** | MDYSSSPTITTCKGEIRGKRLIDEPEFQADAYLGIPYAQPPLGDLRFKKSIPAREWTGTRECVEHPAKCVQMLSELLSQNDKDWPASEDCLYLNVFCSGEYSEKMRPVLVFIHGGGFACGSVKAY<br>GDRGICDGLVRQGLVVVMVQYRLGILGFFSTGDESCPGNFGLWDQLAALQWVQTNIGQFGGDRDHVTICGQSAGGACADLLSLSPHSKGLFHRIIPMAGNARTPWSHKSSTADWCRRYAEDTL<br>GVNAETSEELMAQLRRMPADRLAVALDAVMLRNEAELGFCPVIDGDLFPAPIEELLAAAPALPVMIGLTTLECGLLIPNEDLTDEHLEKAVKALLPKNASEGSTEALSAMYKEVALRKQPAHATWRV<br>LIEATSDRIINIATMKVLHECIRRGAPAYFYVFDYYNKACIGPTANNAPCSDAPHCAELAYVFNGGVIAPFTFTKDDHRVAQLMMRTFANFAKNSNPNDEGSSSWQPCDAAHPGRHMALGMQPRM<br>EEQFLEGRCERWIDFMQ |
| ***Ppa*-UAR-53** | MDCFGAYPPSTTIQTCNGKIEGKRMIHNGPRQVDGIPYAKPPVGELRFKRPMVLDPWEEVLSTREYRARSIQRNFYIWDNIKKGKPSEDCLYLNVFMPCWSPPPTGFPVIVFIHGGSFVMDGASN<br>YGDIEICRNIVSRDTIFVSIQYRVGYLGFWTTGDDACPGNNGLWDQCEALKWINKNIDKFEGDPTLDRLRKISADEFTVNRGDEGGDSEGGTQHITPVLDGYFLSENLGELRKKAKPKPLLTGVTR<br>LEALMFVNGSLVADANTLETFVRSLIPSQIDDQEKIYESLYRRYLPPGLLANKSTFLRGVVEVLSDRFINTPTLQLAKETLQRKEAPVYLYTFEFLNSKSLGLARFWMSIVEASHGSELPYFFGTGIS<br>WKFTFNTDDIEMTHLVAQAFTNFAKCSDPNGLRPHDALPVKWNELTAIRPTSHFVFGKRPKMSDDFFLGRPLTLLEMDEEPPQPGFS |
| ***Ppa*-UAR-54** | MGGSFLFFLLIFVSLTLKNCNADCQIGEPENFMKTFVADRCMEKKVGMEHCRFHIVHEHSSNETSPYTRLLKNEKTGVDVECGVEAHTEGTGVVVGDACSRLLTEDLKLNCKDGEGKDECLREI<br>ECVCTGNKCTTTIGMAYIAAKLQGAPLQSEELALYRFSIYESLTTDGSLNPENYAARHNSSKATTFNPRIHSKRELFGHLRKMFLAICCIIIATAAFYKKVSGWINKLNGKKRKTAKVVQPRLSREAVA<br>ESPEDARIEEYLRGDHLLFEWTRPQQLLTPFVCFPKGFGIDQERITEVEEKPVRCNQTKEYLIVSAIVILAVKQINGKSSLNQPNKCLEEDVEKEYCRFHIVHDHSPNLAKSPEVRIFKSGKTDTDGF<br>ECGKPAHTEATGTVEGDACSRITTEQLELNCKPGDDYDECLRKIQCVCTGKNCTSKVALTYIAAKLQGAPLQSDQLALFRFSIYESLTTDGSLNPENYAARHNSSKATTFNPRIHSKRELFGHLRK<br>MFLAICCIIIATAAFYKKVSRWINKLNGKKRKTAKVVQPRLSREAVAKSPEDARIEEYLRGDHLLFEWTRPQQLLTPFVCFPKGFGIDQERITEVEEKPDIPGMGTSQSSPVPDSRIVETEYGRVQGR<br>RLIAEGDRQVDAFQGIPYAKAPVGELRFKKPQPPDRWEDVLDTKKWGPRAVQKDVFTFGLWKMGPRSEDCLRLNIFAPCWTPPEGGFAVMVFVHGGGFAMGGSNVYGDRSICETLCQKDVIV<br>VSIQYRLGYLGFWTTGDDACPSNLGLWDQTAALQWIQSNIGAFGGNKDNVTVMGQSAGGVSVDLLALSPHSSGLFHKVIPMGGNATARWAFAPSMQKVCERRAKKMKIQEWTDNKHLMEQLR<br>ALPASAFEISMFGAELIDGADLECTPVIDGDFLPVPVDELRKIAAPKPMMTGVAKLEALMFLLMAKKTPGKVRKAATKAVPEHVPNREEEIENLIREYIDLDTAKGKKAVHRALHEVHSDFATNAATL<br>KMIKDTLATHPDNPVYSYVFAYLNPKSYGPLRWYLSVVEATHGMELPYLFGKSLMLKFDFNESDREMCDIFSSAFTNFAKYGNPNGPVSSSSSLPIEWERATIEHTERHYMFDKEFRNEDSYFNG<br>RPSKLLQLRESTLEAPKI |
| ***Ppa*-UAR-55** | MGAFHSHPESRVVRTHYGEVVGRRLYSKDGKSVDAFQGIPYARPPVGDLRFRKPEPPEPWEGIFQAKSFGNRAIQRDLLFFDNWRCMKDVIVVTIQYRLGYLGFWTTNLFHKVIPMAGTASAK<br>WALSPSMKLQCEKKALRLGCDWKDNNDLMAQLRDLPASAFAISLIGADTEKETDLECTPVIDGDFLPSTVDEMRKCAPPKPRMTGVAKMEGLLFLITIKSSFEKLKKLMHRALPKDMPNRESMQN<br>EFLSKYIDIESKPDTDTIYRVMNEIYSDFSMNAPVLKQVQQTIEANPDSPVYNYVFSYLNPKAWGPFRWYLSFLEASHCHELPYIFNKGIIFGFSFTDEDKRMADIFSTAFTNFAKFGDPNGPFGKT<br>ELPVQWKSATKDYPERHYNFDLQPELVDSYFRGRPAALLNYQKQSGVTTAKL |
| ***Ppa*-UAR-56** | MSFYSSSPQMGCCSSRHPSRIVNTKYGPVQGTRIVNDGELRVDAFLGIPFAEPPVGQLRFKKPLPPRHWLGIRQCTKFARRCSQKDYFLHDRIMKKSGSEDCLYLNVFTPVWQPPTEGFPVLVF<br>VHGGGFSMHDAQTYGDEGIARFMVRKGVIVVTIQYRLGYLGFFSTGDEACRGNWGLWDQTMALRWVQENIESFNGNKGNVTLFGQSAGGASVDLLSLSPHSRDLFHKVCPMAGVGTAEWTL<br>HPDLPSKCREKAKKLGVEEKNSEKMMDKLREMKPSCFAAGVEPEQGIESAIMENTLDITPIVDGDFLPDHPSVLRKTAPPKPRMTGVAKLEAVIFLMNYKVTLERLQEMVGRSLPQALPGVEKIR<br>DDLVESYLDTGKLSPNHVMLRAFAEAYGDRSMHVPTFRMVTHSLKYQPEAPVYTYVFEYNATHCTELNYLFGKGIIWPYKFTANDHQMLHLFTTAFTNFAKYGNPNGGSIEAARTPSITPNTLPVF<br>WPAATPENPETHYVFDLSPRVSTTLMYGRPRRIAQYQEQYEK |
| ***Ppa*-UAR-57** | MGGSSSKCPSSKEVSITGGKIIGRRFFHDEDKPVDAYQGIPYAKPPVEELRFRKAEVVDEWEGVKECISFGNRGMQTTNMYEKWKHGPVSEDCLYLNVFSPTWKSSSPSGFPVVFFIHGGAFV<br>SDSTVKYGDILISKYLVRKDVIVVTTQYRLGFLGFWSTGDSSCVDNLGLWDQTCALQWVQDNIESFGGDKNNVTIMGQSAGGASVDLLSLSPHSRDLFHKVIPMAGNGSADWALQENAVEKCR<br>EWAMNKMKIDDGGNSKEFIEKLRLESAQKFATSMEVKVKDNPKDGIQIGPRIDGIFFPKSLNELRKEAGNKPRMVGCCRYEGIMLMMGGKVKKDRSSFESVIASLIPESVNEFKRKRDEMLEEYL<br>KEDDMTKGLAELYSDLFLNISCQQYVLDWLEVGYDHIYLYSFDYFNEKSFGPLSYLLPIKGATHCSEIPYILGKGLIWDFTLNEKDIEMVEKVTGAWSNFAKFGNPNGENNENPLGVEWIPATKENP<br>ERCMNIDWESRMDDHFREGRPMKWIEHRKGL |
| ***Ppa*-UAR-58** | MISVMRPLKLHRDCAPAVLAEVEEGTAEGFVYSFSGKDCEVYLGLPFAAPPIGELRFNKPQRAEPWNGVRKCKRFGPRSIQRDMFWDKVVLQTPQSEDCLYLNVFAPKREEGKTYPVFFYIHGG<br>GFMMDSAAKYGYKEMCEQLITKGIIVVTIQYRLGFLGFFSLGNGSCKIAALEWTKRNARAFGGDPDRITVGGQSAGAVSADLLSLSPVSRDLFTQKIAMGGSSFCHWAVSDTDEISEFCRKKARS<br>LGWKPKGKHYISQEQEDAAMMNYLRVLPAHKFGCHMIGTKEVFSEARLPLAPVIDGEILPAPLPILRAEAPPKPSIGGVGEYESLLFVALGFIRCNGKFLERVQTILARKAKGSSLKEMQEALREIY<br>GDAKGKNKKEVARICIVLLSDLVSNYANYQYMREAERLQEKTYMYSLDFTSRNMWGWMKAVIPLCERRGTHASELLYLFKCNYFVSPLPMDANDRAVADIMPRLFANFVKTGNPNSAETRERGV<br>QWDPIERERKQILSIAPQSTMKREPFDGRMDALDRLFHRLNLDLRYNISRR |

| | |
|---|---|
| ***Ppa*-UAR-59** | GMGGVISLLPLFRRETIHPPSRIIRISTGEIQGFHLPIEEDRQVDIFLGIPFARPPVGERRFQRAEPADHWEGIRLCTRFGPRAPQSDFIWEKLSLGPKSEDCLHLNVFAPSWKKPESGFPVLVFVHG<br>GGFLIDSAVKYGDEGIAKYLCRHDIIVVTVQYRLGLLGFFSTGDSVCPGNLGLWDQTLALQWVRENIGAFGGDPRKVTVFGQSAGGASVDLLSLSPHSRDLFHQVIPMAGNAECEWATVSHDRII<br>DACTKFAERKGFVKPDNELKTSRDMIEFLRTRKGREFERGLLGRKGMDVSKVGLDLAPLVDGDFLPKSINELREEAPKKNIMVGTCEHEGLLFAVLAPNSFDEKGMDKLLAIVIPPELNPEWKKLR<br>EEAKGMYMEGVDEYDRQEVARAYVTLYSDIFVCNGTKEYAEKMSDLGHKVFLYTFDYFNPRSFGILSIRAPFKGATHCTELTYLFGQSIVFSFKFNEADHRMIDLMTRMWANFVKYGNPNGAYDD<br>SSCFTFQWEPTDSTSLKYLKISENCEMAEKYENRRADFWRKVRISKKIPS |
| ***Ppa*-UAR-60** | VFMGGASSYPASRVVDTAYGKVQGRKLVFKGDKCVDAFQGIPFAKPPVGPLRFKKPEKPDSWDGIRDTTKFGARAIQPPLFFFEKFRKSAPSEDCLYLNVFTPCWDPPKEGFAVMVYIHGGGFV<br>MGDAETYGDIGMCENLCTRDILVVTIHYRLAYLGLFTTGDAVCPGNLALWDQTEALKWVQKNIGAFGGNKDNVTVFGQSAGGASVDFLHLSPHSTGLFHKGICLAGNTEAPWALASNMVEQCRK<br>KAKLLGIEENDSSKLIERLREVPADKFGVNMMTQSDKEDFIDLETTPYIDGDFFPESIDELRKKAASKPLMTGVTAEEGIFFMVGKKNNDEGLRKFISHVLRDADNKERLEDQLRKKYVDDRNLKD<br>KNNLLRIQAEVHSDYFMNIPTLKWCKKNAENAQPVYLYLFEHFNVKTLGLGRFVLPILDATHCLELTYLFRKSIFAPFVSTDSEHAVEDNFSIAFTNFAKFGNPNGGNDFRTDLPAHWKPIDIRNATR<br>NFVFTADGGHMNENYFGGRPAEVQTMIQLGR |
| ***Ppa*-UAR-61** | MGILFSYPDSRIVGTTCGSVQGRRLIYKGDKQVDAFQGIPFAKPPVGDLRFRKPEPPEKWEGVKATKKFARRPYQAPFFYIDNLIRGCPSEDCLYLNVFTPCWEPPKEGCPVMMFIHGGGFEIGD<br>TISYGDRNICENIVTRDVIFITIQYRLGYLGFFSTEDEACPGNLGLWDQVAALKWVNENIEAFGGNKNNITVLGQSAGGVSTDMLHISPHSTGLFHKMIPMAGNAHLAVIASNRNMALHSKKKVARL<br>GISDYKNSFELLEKLRRISASKFLEPVKLFRKKNPEEPEFGMFETVPNLDDRDFFPDTVDELRKRAIPKPLMTGITREEGTLFVLTKKCTEQSLNEVISLTCCEARNKDKLAQELRILYVNETLEDSK<br>EKFSTAIVEVASDYYMNAGTLQLCRNTVATQDKPAYLYILDHFNPKVMGILAWFMPIKLATHGGELTYLFNKGFFGKYPSMTKEDQTVMDAFVTSFTNFAKYGNPNGSDPSHSELPTEWTQVTKE<br>NPGRSYVLTGEKNYMREDFFQGRTAKFIEILAKHRSTTSDE |
| ***Ppa*-UAR-62** | MYVIVDSANGRNDKEEFGNRIYSLILDISAGIARSLEMESPKESYKINVVQESTGALSAWSFVLPRMGGYFSYPNSRIVETSYGKIQGRRLVYSGDKQVDAFQKPVKPKSWNGVKETKGFGPRSI<br>HAIDSDSPAQIGSEDCLYLNVFAPVAEPMKDDGYPVMVFIHGGAFEMGDAPTYGDLNICENIVSHDVIFVTIQYRLGYMGFFTTGDDACPGNLGLWDQVAALTWIQRNIEAFGGNKYNITLLGLSA<br>GGASVDMLHCCPHSTNLFQKAIIMAGSAECRWAMNMSMTQQCLVKAERLGVKDWKSSQELLDKLREVPANKLAVIYSEEIKMEKSDFETVPVIDGDFFPASLDELRQRATPKPMMTGVTKEEGL<br>LFILNKTPTEENLRWMIELAAQDSKDKDTLSMKLRDCYVGSVTVEDKDKFMRCIANLASDYYVNAGVLEMCRKTTSIQNEPVFLYVLEHHNPGSMGENENKMFIKDCTHTSEVQFLFKRGRFATP<br>SFTETDNRVTHLFTKTFTNFAKYGNPNGPDVMKSDLPVVWKPIDQENYGRNIVIASDNTYMTNQFFDMGNLFSYPDSRIVETSYGKVQGRRLIYKGDKQVDAFQGIPFAKPPVGELRFKKPEAPE<br>KWDGVKDTKKFGPRSIVGAYFHWAIDSLLMGKTSEDCLYLNVFTPCWKAPENGFPVMVFIHGGAFELGDAISYGDIGICENIVTQGVIFVTIQYRVGYLGFMTTADSSCPGNNGLWDQTAALRWV<br>NDNIDEFGGDKKNITVLGQSAGGASVDMLHISPHSTNLFHKMIPMAGSAECRWASNKNMPQQCRKKAANLGITDYANSEEFLEKLRGVPADKFGVVFLKRDKEPEVDFETVPLIDGDFFPESFD<br>ELRKKATPKLMMTGVTKEEGLLFIPGKKATEDDLKEVITLAIHDAKDKEKLANDLKSAYFPAGIPDDKEVFLRRIANIASDYWFNAAVAEMCRKTVAIQKEPVYFYLFEHYNPKVMGFMGKQMPIQD<br>CTHAAELIYLFKKGLFANVALTEDDIRVMYLFTTAFTNFAKYGNPNGAEESKSDLLVYWEPLDKENHSRNFVITSNEPYMSDNFFEGRVAKFVETVKSHA |
| ***Ppa*-UAR-63** | MFLLIHYYSTFPIFKMANPISYPDSRIVETSYGKIQGRKLIHEGEKQCEGFQGIPFAKPPVGALRFKRNIIQKPEAPERWEGVKETKKFGPRAIVGSYDGWDLEYDAGTPSEDCLYLNVFSPCWKA<br>PQGGFPVMVWIHGGAFEVGDPMAYGDVNICENIVKRDVVFITIHYRVGYLGFFTTGDAACPGNAGLWDQVAALKWINENVAAFGGDKSNITVLGQSAGSVSVDMLHLSPHSTNLFHKRICMSGT<br>VDSRWVVNKNMPIVCKAKAARLGVTEYGSNEEMLEKLRAIPAEKFGIMNMNREKEPDVDLETVTFLDGDFFPEPFDVLRAKATPKPLMIGVNNEEGLLCFMGKKPTTEAEVDEVITWAIHDAKDK<br>QKLFDELKSLYFEGGLPEDKDVHDPVYLYTFKHNNPAIFGAVGTLLPLQDATHSTEMFYLFHKGLFGNPPLTDDDKYIMDLFTTTCTHFAKFGNPNGASNHGTDLPVQWKPLDSESHTRNFVFIAD<br>KPHMQEQFFEGRPARFAEIVKKHRAS |
| ***Ppa*-UAR-64** | MGNRQSCPDSRIVETSCGKVQGRRLIYKGERQVEAFQGIPFAKPPVGKLRFKKPEPPEKWDGIKETKKFGPRAIVTPFVNGFVENLLVTRGVVFITIQYRLGYLGFLSTGDAACPGNNGLWDQTA<br>ALRWVNDNIEAFGGNKNNITLLGQSAGGASVDLLHLSPHSTNLFHKVIPMAGNAECRWATNKNMPQQCRNKAARLGITSYETSEELLDKLRALPAEKFAVNFRNREKEPDVNLETVPILDGDFFP<br>ESIDEMRKKAKPKPLMTGVCTEEGLFFIPGKKPTEQDLNEVVNFAVHEAKDRNAIAADLKSVYLPDGIPADKDVFMRSIANIASDYFFNAGTIELCRKTVALQGLNTTEQVTPHFPPHSTPFAAPAD<br>PPLSIPKCWFGYNKPSSKEWKQLMEFDDEDAEDIMEEYLSNSTQTSDSDQTNSPRILLDHRANGERANPGEYSELIKHEKNIDRFVLAICPSEKTLTSSFGTWETKRLRHEAVYLYQFEHFNPKV<br>MGFAGRQLPIQGRTESRYTEFHPKQFFSDATHASELIYLFKKGLFTLTEVSLTDEDKHAMHLFTTAFTNFAKYGDPNGSNDDKSDLDVYWKPLDKMNHSRNFVFASVKPYMSDQFFEGRIAQFV<br>DVVNKHRAEIHASSYPSSVSMDMSHSRTHLLPPDPMLGTRILSRKPTRFRCALAEVKTLTLTCVIDRCTKCVDGVCSDYPLTQQSTTSRNRKRAAAPEERHSQSYSFAWNCSTSRYGRTACDGT<br>GCTN |
| ***Ppa*-UAR-65** | MGDRQSCPDSRIVETSCGKVQGRRLIYKEERQVEAFQGIPFAKPPVGKLRFKKPEPPEKWDGIKETKKFGPRAIVTPFVNGFVENLLRGTPSEDCLYLNVFTPCWKAPQGGLPVMIFVHGGAFE<br>LGDAQIFGDLNICDNIVTRGVVFITIQYRLGYLGFLSTGDAACPGNNGLWDQTAALRWVNDNIEAFGGNKNNITLLGQSAGGASVDLLHLSPHSTNLFHKVIPMAGNAECRWATNKNMPQQCRNK<br>AARLGITAYETSEELLDKLRALPAEKFAVNFRNREKEPDVNLETVPILDGDFFPESIDEMRKKAKPKPLMTGVCTEEGLFFIPGKKPTEQDLNEVVNFAVHEAEDKNAIAADLKSVYLPDGIPADKDV<br>FMRSIANIASDYFFNAGTIELCRKTVALQVEFDDEDAEDIMEENPYPCTTKQRTKSFLELIKHEKNIDRFVLAICPSEKTLTSSFGTWETKQLRPKATAKRGDEAVYLYQFEHFNPKVMGFAGRQLPI<br>QDATHASELIYLFKKGLFTLTEVSLTDEDKHAMHLFTTAFTNFAKYG |
| ***Ppa*-UAR-66** | PLHNYPLAMHEKQFPNSRVVQVAQGKVQGRRLIYEGEKQVDAFQGIPFAAPPVGELRFKKPQPLFHWDGVKETKAFRDRAIQAPKYNGDFEANGIPSEDCLYLNVFTPCWKPPVEGFPVMIFIH<br>GGGFETGEARTYGDENICENIVTRGVVFVTIQYRVGYLGFFTTGDSVCPGNLGLWDQTEALRWIQINIGSFGGNKDNITVIGQSAGGASTDFLHLSPHSTGLFHKMICMAGNAECRWASNQKMP<br>LHCRSKARRLGINWDSSEELISELRKIPADNFGVNFFKKEKEDDVDFETVVYNDGDFFPESFDKLRLKAKPKPTITGVTKEEGILMMMFMKINKRTAEWIVKAASQSASDKEKLARILSARFDGLW<br>EDEEKLGRAIANIASDYYFNAGTLEQCRKTVAIQKEPVYLYTFEHWNPESLGALLSIVPIEDVTHACELFYFFKHSLFGASNPEVTEEQQRVIDRFTTAFTNFAKCGNPNGMGSSTLPARWDPITRE<br>NYSKNYVFETETCSMRDDFFEGRTAEFIRIVKEHSAERHRSSL |
| ***Ppa*-UAR-67** | MAPGKQFPNSRIVQTAQGKVQGRRLIYEGEKQVDAFQGIPFAAPPIGELRFKKPQPHPSWNGVRETKAFSARPIQAKKYPQDYDMNGVPSEDCLYLNVFSPCWEPPETGFPVMIFIPGGGFEC |

| | |
|---|---|
| | GEAKTYGDDNICENIVTRDIVFITIQYRVGYLGFFTTGDSVCPGNLGLWDQTEALKWVQANIEAFGGNKNNVTVVGQSAGGASADFLHISPHSTGLFHKVICMAGTADCRWTSYDKMPLHCRNK<br>ARRLGITSDSSEEIIAELRKVPAEQLVVNITRREKEDDAFMETVVYNDEDFFPASFDELRARAKPKPMITGVTKEEGLLMMLSMKLNKKTALYFTTLASHSAKDNKLLEKDFSRRFDGLVEYSDQFG<br>VTLANFVSDYFFNAGTLELCRKSVEIQKEPVYLYTFEHWNPEVMGFMVDMLPYKDVTHTCDLYYLFKIGTLGDFRPEISENEQRLIDEFTTAFTNFAKFGNPNGSNIVATELSSSWEPVTRENHSR<br>NYVFKSENCAMNEQFFGGRTEEFIRIVNTHNANQPRSSL |
| ***Ppa*-UAR-68** | TLSRATRRTFLTSQMTADCPPSPIVETSYGKVQGRRLIHAGDRQVDAFQGIPFAAPPVRELRFKKPQPPASWEGVKETKKFSARNIQVPFPGLPEDELHGVMSEDCLYLNVFTPCWEAPEKGFP<br>VMVFIHGGAFVFGEASSYGDIRICENIVSRGVVFVTIQY |
| ***Ppa*-UAR-69** | MPADCPPPSRIVMTTHGKVQGRRLVHAGDRQVDAFQGIPFAAPPIGELRFKKPQPPAPWAGVRETKKFAPRCIQVPFPGLLEDELHGEMSEDCLYLNVFTPCWEAPEEGFPVMVFIHGGAFVFG<br>EASSYGDIGICENIASRGIVFVTIQYRLGYLGFLSTECGQRKCGSAASQSAFDNSKVDTVVHFTLYLVDRAGERVCNMKRRWYHISSSTSDLFHKAICMAGTAECRSATRSCSSMAAQSIRKAACL<br>GVTEFANTQELLDKLRQIPAEKFAVSLFEAPTRGDRIDFGANFIIKETCPCLDGDFLPESLDALRAKATPKPLLNGVTKEEGLFLMPGRRSTLEGLQETLQYVTLDCTKQDAMKKELCSCFVGDAK<br>PEDPACIRAQAGMVADSIFVAGHLELCRKTVAIQTEPIYLYVFEHFTGPSILGAFNPNGWDDSVNDLSVPWKPITKENPALNYVFTSNEPMMSNDLFEGRTAAFIEIHRRHK |
| ***Ppa*-UAR-70** | MPADYPPSRIVTTSHGNVQGRRLVNAGDRQVDAFQGIPFAAPPIGELRFKVGRLFVPEFDTRSKPQPPSPWDGVRETKKFASRNIQVPFPGLPEDELHGEMSEDCLYLNVFTPCWEAPEGGFP<br>VMVFIHGGAFVFGEASSYGDIGICENIVSRGIVFVTIQYRLGYLGFLSTGDSVCPGNIALWDQTEALRWVQSNIGAFGGSKHNVTLLGQSAGSASVDLLHLSPHSTNLFHKEICMAGTAECRSSTR<br>SCALKHGRAEHPESGDFGRDRLFNKLREIPAEKFAVSLFEAPTPGNRTDFVGNLRNMNAQLVMITVCLTDVTHTNELFYLFKKGFFSDPEITESDSKLIDVFTTALTNFAKYGNPNGSDDSVYNLSV<br>PWKPITKENPALNYVFTSDEPKMSNDLFEGRTAAFIEIHRRHK |
| ***Ppa*-UAR-71** | MPDIVYPPSRIVETAYGKVEGRRLISEGERKVDAFQGIPFAAPPVGDLRFKKPQQPESWEGVRETKQFAAPQIQYLRAYQDNKDKGVPSEDSLYLNVFTPRWEAPEGGFPVMVFIHWGAFAFGE<br>AISYGDIGICENIVTRGIVVVTIQYRLGYLGFFSTGDAICPGNFGLWDQIEALKWVQLNIGAFGGDKNNVTVAGLSAGSVSVDLLHLSPHSTNLFHKVFCMAGTAECRWAIHPDMVAQSRRKAKRL<br>GVEKFSNDEELLAQLRLLPAENFAVDEDNQDKEDKAHFEVGPCLDGDLFPESIESLRRKATPKPILTGITKDEGLLMMPGRKSTSEGLEETLLEATRDCRRKEEMKAELISRFVGDTKPDGPVYMR<br>AQAGMVADTYFVAGVVELCRKTVQLQTNSIYLYVFEHFNPSVLGKLGEAMAFQGPTHGCEIFYLFKKGLFGNPELSETEKRVMDIYTTACTNFVKYGNPNGSNDSGSDLPRHWKPITIENPALNY<br>VITSDEPKMNNLLFEGRTDAFINIHNMYN |
| ***Ppa*-UAR-72** | MGESDKRSKYQGIPFAAPPIGEMRFRKPVRPAAWKGVRQATAFSARCIQHPKNPQDYSINGIPSENSLYLNVFAPCWKAPEGGFPVMMFIHGGLYLNGEASSYGDIGICENIVSRGVVFVTIQYRL<br>GYLGFLSTGDAVCPGNFGLWDQVAALEWVQQNNNVTLVGQSAGAASVDLLHLSPHSTGLFHKAICMAGSAECRWALHPEMEAQTRQKAKRMGVQYDTNEELLEKLRKLPAVEFGVGIFSQVK<br>ADHLDMEVGPCLDGDFFPESLDVLRKRAKPKPFLVGVTQEEGLYAMAGRTSSTTGLADTLKEATADCVSREAMKASLRSHFIGYASPTNPSYIRAMAGMASDALFVAGIAGHCAKTVAIQKEHMY<br>LYVFEHFNPAIMGYITPLLPFHSATHTCELFYLFKKGVLGEPPMTETEQRVMDTFTTALTNFAKYGDPNGAVECETDLPTRWDPITKQNPHLNYVITSGEPVMSDELFEGRTEDFIDIRDKYN |
| ***Ppa*-UAR-73** | CQMAPTSYPQSRIVETAYGKVQGRRLVHEGERKVDAFQGIPFAASPVGELRFRKPVPPSRWEGLKQTSEFAPRSIQHAKNPQDYDINGIPSEDSLYLNVFTPCWNAPEEGFPVMVFIHGGGYIN<br>GEAKSYGDIGICENVVTRGIVFATIQYRLGYLGFLSTGDTVCPGNFGLWDQVEALKWIQMNITAFGGNKNNVTLVGQSAGAASVDMLHLSPHSTGLFHKAILMAGSTYCRWAMHPDMAEQTRQK<br>LKRLGVENVENSEELLKKLRGLPAKEFGVGIYTQVKEADIELETAPCFDGDFFPEPLEVLRRKATPKPFLIGVTEEEGLFPMAGRTSCAASLSDTLRDVTGECRNKEQMKDALLKRFIGDATPNDS<br>SYLQGMAGMMSDSFFVAGSADLCKKIVEIQEEPVYLYVFEHFNPGMMGYLTPFMPLQKATHTCELFYLFKKALLVDLPLTETEMKVTNIFTTAFTNFAKFGNPNGTEAETELSVRFEPVTKKNPHL<br>NYVITLKDAVMNDELFEGRTAAFIDIRNEFK |
| ***Ppa*-UAR-74** | MGCRWSSYPDSTIVETKYGKVQGRRLIREGEKQVDAFQGILFAKPPTGELRFKKPEPPEWWHGVKETKKF |
| ***Ppa*-UAR-75** | MAHIVYPPSRIVKTAYGEVRGRRLIYEGERQVDAFQEYPSPLHQLAIFDSKVDETKRNSEIPKNFNYLHFSGKRKPQPPARWEGVKVTTEFAARSMQGPKNPQDYVNGIPSEDSLYLNVFTPSWD<br>VPEGGFPVLVFIHGGAFAMGDASWKSKMFCGECGQINLQVTRGVIFITIQYRLGYLGFLTTVEALKWIQLNIGAFGGDTMNVTLMGKSAGAASVDLLHLSPHSTNLFHKAITMSGTAECRWALLAK<br>LRLLPAEAFGVKMFQQEKEDNADFEVGPCLDGDLFPEPLDVLRARATAKPFITGVTKEEGLLMMVGRQSTPEGLEETLMDATRDCQEIDAMKCELRGRFIGDTKQDDPAYMRAQAGMISDAVFV<br>AGTLEHCRRTVAIQKLPVCLYVFEHFTPSILGYLEGALPIQDTTHACEIFYLFKKGAFGDPEINVTEKRVMNIFTSAFTNFAKFGNPNGSDDATSALPVQWDSITEENPALSYVFTSEEPKMCTDLAE<br>LLSSSIFATNTSEVLNHE |
| ***Ppa*-UAR-76** | MPNAVYPPSRIVDTAYGKVEGRRLINEGEKQVDAFQGIPFAAPPIGELRFKKPQPPVSWDGVKETKKFAARSIQGPHDPHDYELNGIPSEDSLYLNVFTPCWDAPEGGFPVMVFIHGGAFVMGD<br>ASSYGDIGICENIVSRGIVFVTIQYRLGYLGFLSTGDSICPGNFGLWDQVEALKWIQLNIGAFGGDKTNVTLLGQSAGAASVDLLHLSPHSTNLFHKAITMAGTAECRWAVHPNMHLRSVRKACDLI<br>SGEFTGSEEAKAGPCLDNDFFPESLDVLRAKATPKPFITGVTKEEGLLMMTGRQATPEGLEETLVDATRDCKNIEGMKAELVSRFIGDTKQDDPAYMRAQAGMISDAFFIAGTLEHCRKTLSIQKQ<br>PVFLYVFEHFTPSILGFLGGAMPFQEVTHACEIFYLFKKGVFGTPEINETEKKVIDIYTTACTNFAKYGNPNSSKDSALPIHWDSIIEENPTLNYVFTSDEPKMTSKLFEMSDAVYLPSRIVETAYGKV<br>EGRRLINEGDRQVDAFQGIPFAAPPIGELRFKKPQTPACWKGLKETKKFAARCIQEPLDSQDYDLRGIPSEDSLYLNVFTPCWEAPEQGTNSFTDHFFSGFVMGETSSYGDIGICENIVSRGIVFIT<br>IQYRLGYVGFLSTGDGSCPGNFGLWDQIEALKWIQLNIGAFAGDQSNVTLLGQSAGGASVDLLHLSPHSTNLFHKAVTMAGTAECRWALPPQLPIESIRKACKASVGEFTDNEEMIAKLRLLPAVE<br>IAEIFKMVKEEKVVYDNVETGPCLDGDLFPESLDALRAKATPKPFITGVTKEEGLLMMTGPRGIPEGLEETLSYVTKDCSRQAEMKVEHIRRPFSIILAIIYIGFISIFRSLPYRSKQFHLRWKYATASL<br>ETYNPMIQPTFERKPG |
